# Forest biodiversity benefits from warmer microclimate, but only in colder regions

**DOI:** 10.64898/2026.10.20.760414

**Authors:** Kerstin Pierick, Roman M. Link, Orsi Decker, Johanna Asch, Petr Baldrian, Claus Bässler, Julian Bittermann, Vendula Brabcová, Pia M. Bradler, Medha Bulusu, Marc W. Cadotte, Lena Carlson, Simone Cesarz, Anne Chao, Alice Claßen, Nico Daume, Benjamin M. Delory, Sebastian Dittrich, Bronwyn Lira Dyson, Martin Ehbrecht, Nico Eisenhauer, Heike Feldhaar, Joshua Ferenczy, Andreas Fichtner, Ronja Hausmann, Csenge V. Horváth, Michael Junginger, Patrick Kacic, Lisa Köstler-Albert, Daniel Kraus, Ludwig Lettenmaier, Clàudia Massó Estaje, Oliver Mitesser, Akira S. Mori, Jörg Müller, Goddert von Oheimb, Marcell K. Peters, Ruth Pickert, Fons van der Plas, Jeisson Rodriguez Valenzuela, Julia Rothacher, Kai Sattler, Michael Scherer-Lorenzen, Jens Schlüter, Rike Schwarz, Ingolf Steffan-Dewenter, Jean-Léonard Stör, Leah Vogelfänger, Clara Wild, Dominik Seidel, Bernhard Schuldt, Christian Ammer

## Abstract

The diversity of many forest-dwelling organisms depends on forest structure, but it remains unknown to which degree this is mediated by microclimatic temperature, and whether microclimatic buffering is universally beneficial for forest biodiversity.

In 234 broad-leaved forest plots across nine climatically differing regions in Germany, we measured micro-and macroclimatic temperatures, canopy cover, and deadwood volume after experimental manipulations of forest structure through thinning operations and gap cuts. We surveyed 21 taxonomic groups, representing plants, bacteria, fungi, and animals, to assess how their diversity depended on the predictors.

Multi-taxon diversity, and diversity of ten separate taxonomic groups, increased from closed to open canopies. Only necrophagous beetles were more diverse in closed stands. These effects of canopy cover on diversity were partly mediated by microclimatic temperatures. Thereby, diversity responses to microclimatic temperature depended on the macroclimatic context. In regions with colder summers, locally warmer microclimates in canopy gaps had strong positive effects on biodiversity, while these effects were not found in regions with warmer summers.

Consequently, management strategies aimed at multi-taxon forest diversity conservation should be chosen based on the regional climate, favoring canopy gaps primarily in colder regions, and taking future climate warming into consideration.

---

In the context of the ongoing biodiversity change^1–4^, forest management approaches in Central Europe and their impact on forest biodiversity need to be re-evaluated with the aim to better integrate biodiversity conservation in production forest landscapes^5–8^. Through its modification of forest structure, including canopy structure and deadwood availability, forest management impacts biodiversity across a broad array of forest-dwelling organisms^9–15^. Deadwood as a resource is essential for saproxylic species across many taxonomic groups^16–18^. Canopy structure may affect diversity directly, for example in communities of flying organisms of larger body size like bats and birds where species can have specific requirements for flight space and perches^19–21^. However, indirect effects mediated by microclimate are often assumed to dominate the diversity responses to canopy structure of most taxonomic groups^14,22^. Mediation in this context means that canopy structure has a causal effect on microclimate, which in turn affects diversity^23–25^. Among the many microclimatic factors that can potentially mediate forest structure effects on diversity, temperature is of particular interest, since it is not only a known driver of biodiversity at larger scales, but also predicted to change markedly in the future^26,27^. As below-canopy temperatures in forests depend to a high degree on macroclimatic weather conditions, the same forest structure may in the future result in markedly different below-canopy temperature regimes compared to today^28,29^. In order to predict forest biodiversity responses to climate change, it is therefore especially important to quantify the extent to which the effects of canopy structure on forest biodiversity are mediated by microclimatic temperature.

To date, there is no consensus whether open canopy structures such as larger canopy gaps featuring less buffered microclimatic conditions tend to be beneficial or detrimental for biodiversity in temperate forests. On the one hand, it is often argued that forest management should aim at achieving maximized temperature buffering through dense, undisturbed canopies to provide cool microrefugia for forest organisms that would otherwise suffer from heat stress^30–35^. On the other hand, diversity metrics of many taxonomic groups of forest species seems to be higher in canopy openings than in closed stands^36–39^, and respond positively to reduced canopy cover in the context of natural disturbances^40,41^ or successional stages^42^. Typically, biodiversity responses to forest structure vary widely between taxonomic groups, underlining that there is no single forest structure meeting the requirements of all taxa^14,43–46^. Diversity responses to microclimatic temperature have especially been observed for mobile, ectothermic animals, though heterogeneous in direction^47^. In addition to this taxon-specific variation, recent evidence also suggests that biodiversity responses to microclimatic conditions can depend on the larger thermal context. In a Central European mountain range, effects of vegetation structure and microclimate on biodiversity depended on altitude, i.e., the macroclimatic temperature (with directions of the effects differing between taxonomic groups)^47–50^. For saproxylic beetles, similar interacting effects of microclimatic and macroclimatic temperatures on taxonomic diversity were observed along a latitudinal gradient in Northern European boreal forests^51^. While it seems plausible that such a dependency of microclimate-diversity-relationships on the macroclimatic context could be a common pattern, this has rarely been investigated so far.

Here, we present evidence from a Germany-wide manipulative field experiment set up in nine broad-leaved production forest regions along a climatic gradient shaped by elevation (lowland – submontane) and continentality. In 234 plots, canopy cover and deadwood availability were manipulated experimentally, creating structural gradients from closed stands over moderately open stands to canopy gaps 30 m in diameter^52^. We measured canopy cover using ground-based LiDAR, assessed the total volume of available deadwood, and derived both micro-and macroclimatic temperatures from air temperature measurements in the plots (Fig. 1). Specifically, we measured average daily maximum temperatures in summer, of which we used regional means under closed canopies as a comparable regional baseline (hereafter termed “macroclimatic temperature”), and local deviations from this baseline to describe smaller-scale variation (“microclimatic temperature”, Fig. 2). Taxonomic α-diversity of 21 taxonomic groups, covering bacteria, fungi, plants, vertebrates, and invertebrates (Fig. 1) was measured in each plot and quantified as coverage-standardized Hill numbers^53^ of orders *q* = 0, 1, and 2 (no weighting, moderate weighting, and strong weighting by abundance or incidence frequency), and additionally aggregated into a multi-taxon diversity index^54^. Using Bayesian generalized linear mixed models, we tested whether multi-taxon diversity and diversity of the taxonomic groups depended on canopy cover, microclimatic and macroclimatic temperature, and deadwood volume (Fig. 1). We used a causal inference framework^23^ to test whether the causal diagram in Fig. 1 is consistent with the observed data.

**Figure 1.**
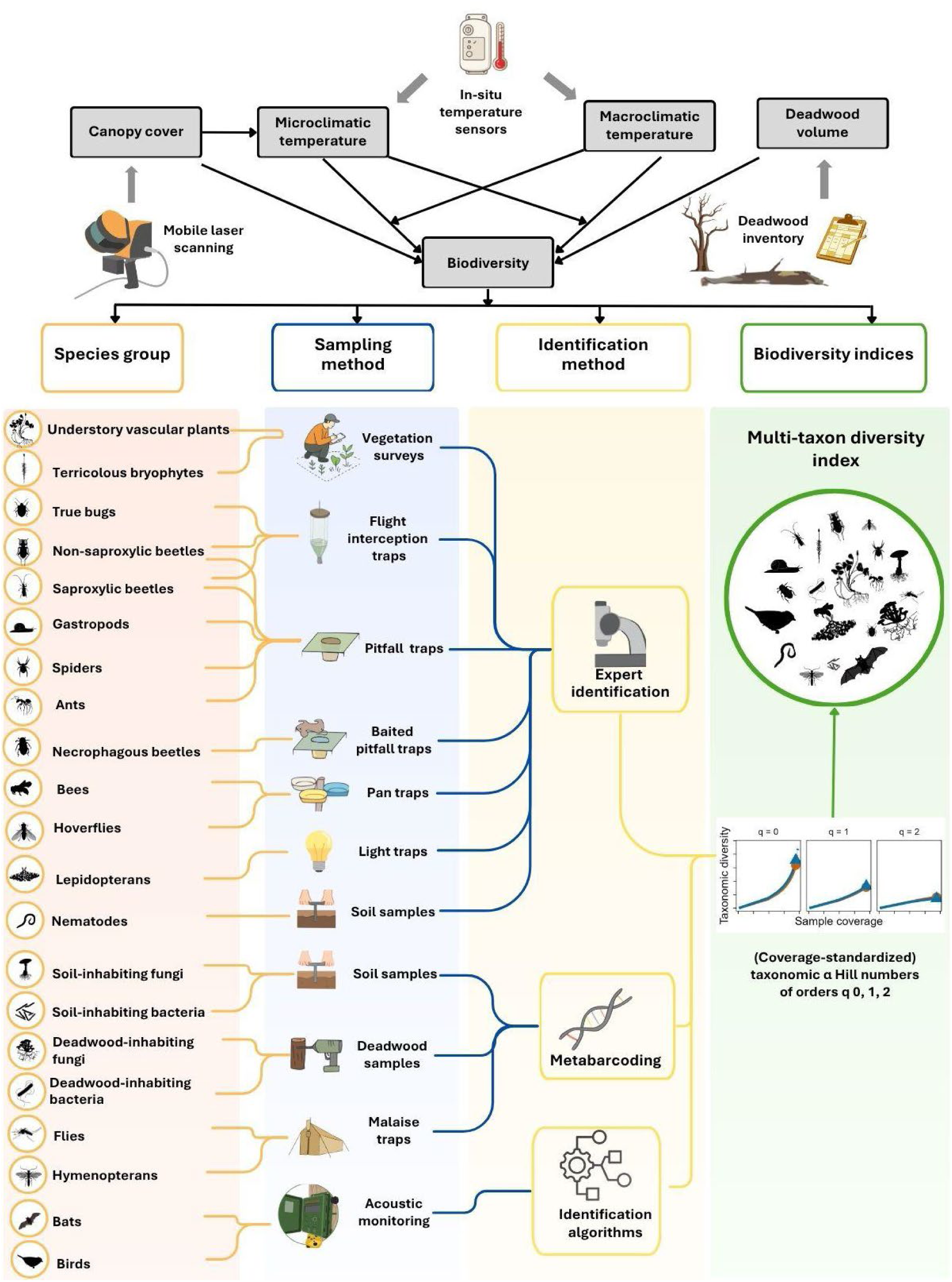
Conceptual scheme of the methodological approach: The top shows a causal diagram representing the tested relationships in this study, and the methods to obtain the variables involved. Canopy cover was assessed with mobile laser scanning, micro-and macroclimatic temperatures were derived from in-situ temperature measurements, and deadwood was manually inventoried. Taxonomic α-diversity of 21 groups was measured through various sampling and taxon identification methods and quantified as coverage-standardized Hill numbers, from which a multi-taxon diversity index was calculated.

**Figure 2.**
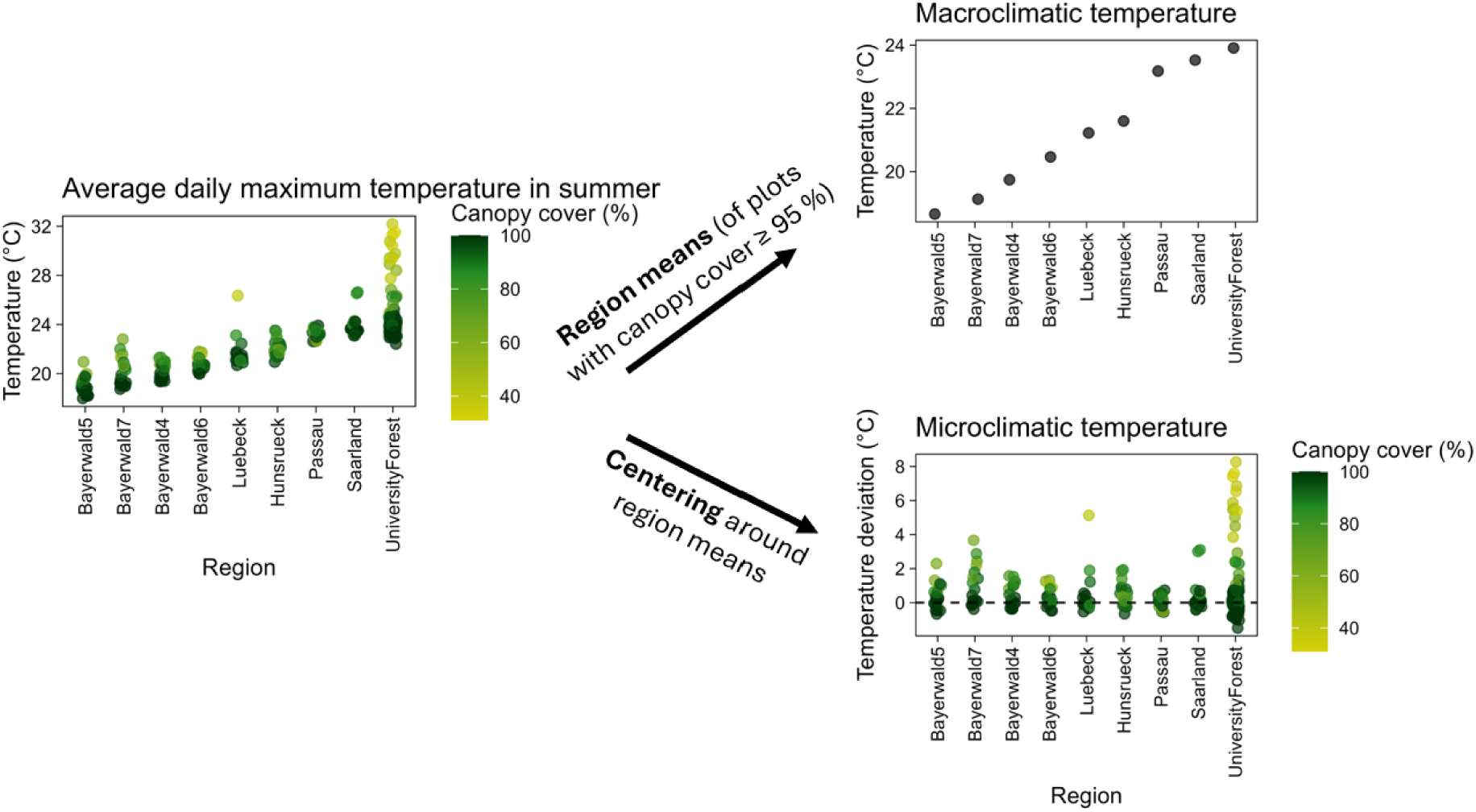
Derivation of microclimatic and macroclimatic temperatures from in-situ measurements. In a first step, time series of half-hourly air temperature measurements from June to August 2023 were aggregated to the average daily maximum temperatures during this period. These data contain both a macroclimatic signal (general differences between regions, caused mainly by elevation and continentality) and a microclimatic signal (within-region differences caused by local canopy cover, topography, and other local factors). We extracted the macroclimatic signal by calculating regional means based on all plots with a canopy cover of at least 95%, in order to base the regional means on comparable conditions. To obtain microclimatic conditions, we then centered the data around these regional means. This way, the microclimatic temperatures can be interpreted as the deviation in °C from the regional average for closed stands.

We hypothesized that (i) multi-taxon diversity responds only weakly or not at all to gradients of canopy cover, deadwood volume, microclimatic and macroclimatic temperature because of specific, often contrasting responses of different taxonomic groups, (ii) for many taxonomic groups, especially mobile, ectothermic animals, canopy cover effects on biodiversity are partially mediated by microclimatic temperatures, and (iii) in colder regions, warmer microclimates tend to affect biodiversity more positively than in warmer regions.

## Results

### Canopy cover effect on microclimate

Plots with a lower canopy cover had a higher average microclimatic temperature (i.e., within-region centered average daily maximum temperature in summer), with 21.7% (95% credible interval (CI): 11.9-31.8%) of the variance in microclimatic temperature explained by canopy cover. Increasing canopy cover from 80% to 100% reduced the predicted microclimatic temperature by 2.06°C (95% CI 1.68-2.42°C; Extended Data Fig. 1, Supplementary Table S1).

### Multi-taxon diversity

Multi-taxon diversity (i.e., plot averages of re-scaled diversity per taxonomic group^54^) was higher under more open canopies. It furthermore increased with microclimatic temperatures, but the strength of this relationship depended on the macroclimatic temperature: in the coldest region, an increase of microclimatic temperature by 1 standard deviation (SD) above the regional average increased predicted multi-taxon diversity (*q* = 1) by 0.029 (0.47 standard deviations of multi-taxon diversity), while an analogous change in temperature led to a predicted increase of only 0.013 (0.20 SD) in an average region and 0.004 (0.06 SD) in the warmest region. The increase of diversity with microclimate was statistically credible on a 0.05 level for all but the two warmest regions, which is, in combination with the credible effect of canopy cover on microclimate, consistent with a temperature-mediated canopy-cover effect in all but these two regions (Supplementary Table S2). While warmer regions tended to feature higher multi-taxon diversity, this effect was not credibly different from zero. Deadwood volume did not credibly affect multi-taxon diversity (Fig. 3a, Supplementary Tables S3, S4). These outcomes were consistent across Hill numbers of different orders (Fig. 3b, Extended Data Fig. 2). The models explained 60.2 – 81.3% of the total variance of multi-taxon diversity (conditional R^2^, fixed effects plus random effects accounting for the nested plot design), with the fixed effects alone explaining 29.5 – 34.4% (marginal R^2^). After accounting for residual correlations between deadwood volume and both macroclimatic temperature and canopy cover not present in the initial causal diagram in Fig. 1, our structural model was consistent with the data (Fisher’s C test, χ^2^ = 1.362, df = 6, p = 0.968) and indicated both a direct negative effect of canopy cover on multi-taxon diversity and an indirect pathway mediated by microclimatic temperatures (Fig. 4; Supplementary Table S1, S2).

**Figure 3.**
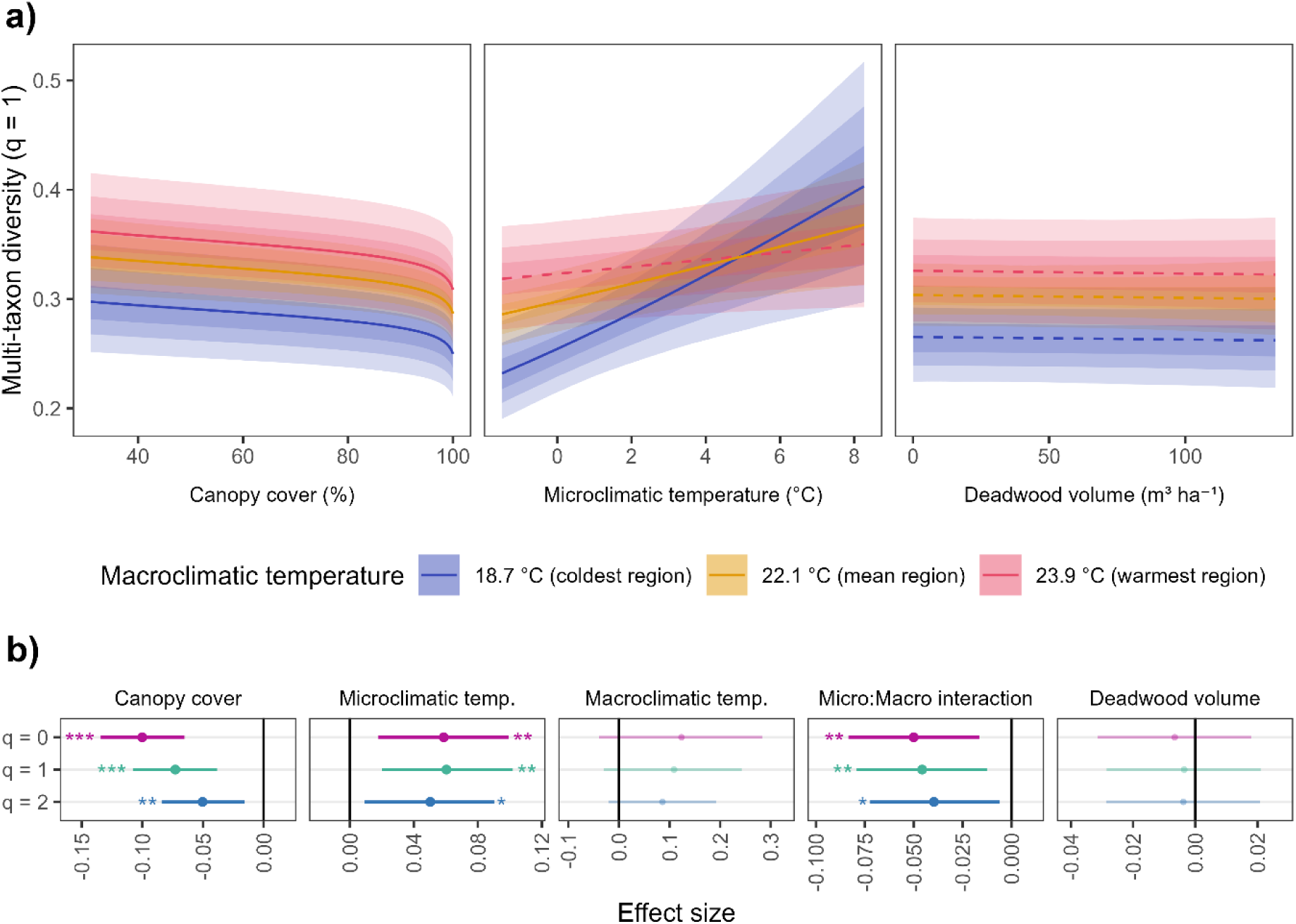
Results for multi-taxon diversity. a) Conditional predictions from the model for multi-taxon diversity of order q = 1 for three levels of macroclimatic temperature: the coldest and warmest region in the dataset, and a hypothetical region of average macroclimatic temperature. Solid lines represent the posterior mean for relationships credibly different from zero, dashed lines represent relationships that do not differ credibly from zero (Supplementary Tables S2, S3). The ribbons show, in order of increasing transparency, 50%, 80%, and 95% credible intervals. The variables were back-transformed from the scale used for model fitting onto their original scales, causing the non-linear relationship for canopy cover. b) Effect sizes (change in linear predictor after increasing variable by 1 standard deviation) from all models for multi-taxon diversity (posterior mean ± 95% credible intervals). The effect sizes are given for canopy cover, microclimatic temperature, macroclimatic temperature, the interaction between micro-and macroclimatic temperature (with negative effect sizes meaning that effects of microclimate are less positive under warmer macroclimates), and deadwood volume.

**Figure 4.**
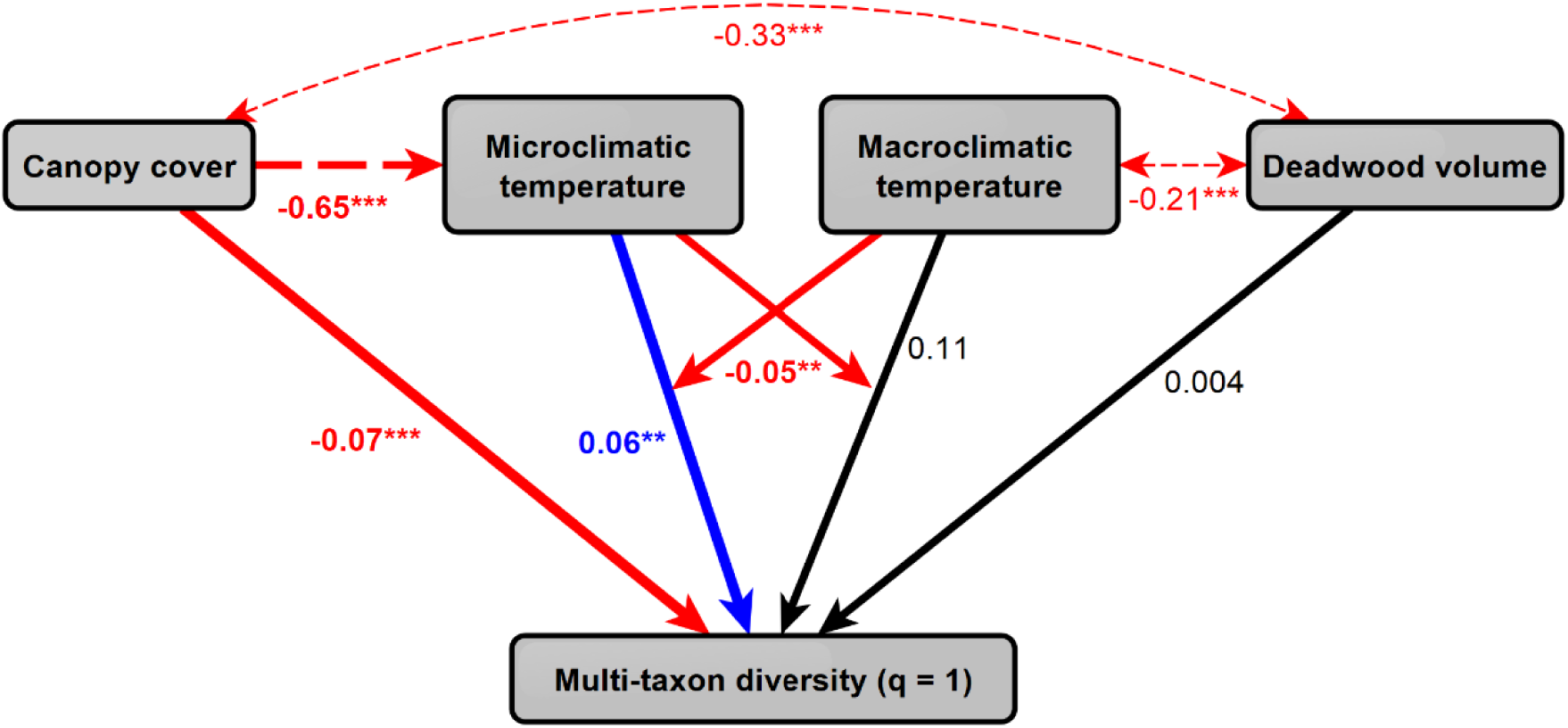
Path diagram of the estimated relationships between the analyzed variables. Relationships displayed with solid lines were assessed in a beta GLMM with multi-taxon diversity as a response (see Fig. 3), relationships displayed with dashed lines were assessed with multi-and bivariate linear models with Gaussian error distributions. Because of the nonlinear relationship enforced by the logit link function, the effect sizes from the beta GLMM for diversity, the Gaussian linear model for microclimatic temperature and the residual correlations are not directly comparable. Fat lines indicate relationships that were part of the initial path diagram (Fig. 1), thin lines are credible correlations between variables that were not included in our initial conceptual model. Positive effects are highlighted in blue, negative effects in red, and effects not credibly different from zero as black arrows. Asterisks indicate parameter estimates credibly different from 0 on the 0.05, 0.01 and 0.001 level, respectively. The overall structural model in this form was consistent with the observed data (Fisher’s C test, χ^2^ =1.362, df = 6, p = 0.968).

### Individual taxonomic groups

For the Hill number of order *q* = 1 (moderate weighting by abundance or incidence frequency), the diversity of 10 out of 21 taxonomic groups was credibly negatively impacted by higher canopy cover, including understory vascular plants, several insect groups (e.g. true bugs, bees, hoverflies), and also soil fungi, soil bacteria, and birds, resulting in a negative overall effect of canopy cover. Necrophagous beetles were the only taxonomic group that was more diverse under higher canopy cover. The diversity of true bugs, hoverflies, and hymenopterans increased with microclimatic temperatures, while true bugs, ants, non-saproxylic beetles, and gastropods were more diverse at higher macroclimatic temperatures than in cold regions, resulting in positive overall effects of microclimatic and macroclimatic temperature (Fig. 5, Supplementary Tables S5, S6). There was a negative overall interaction between micro-and macroclimatic temperatures, with credible group-level effects for hymenopterans, flies, non-saproxylic beetles, spiders, and necrophagous beetles (Fig. 5). Accordingly, for all of these groups, diversity increased strongly with microclimatic temperature in the coldest region, but microclimatic temperature had no credible effect in the warmest region (Extended Data Fig. 3, 4, 5). The total amount of deadwood did not affect diversity of any of the organism groups. Results for other orders of *q* showed largely similar tendencies (Extended Data Fig. 6).

**Figure 5.**
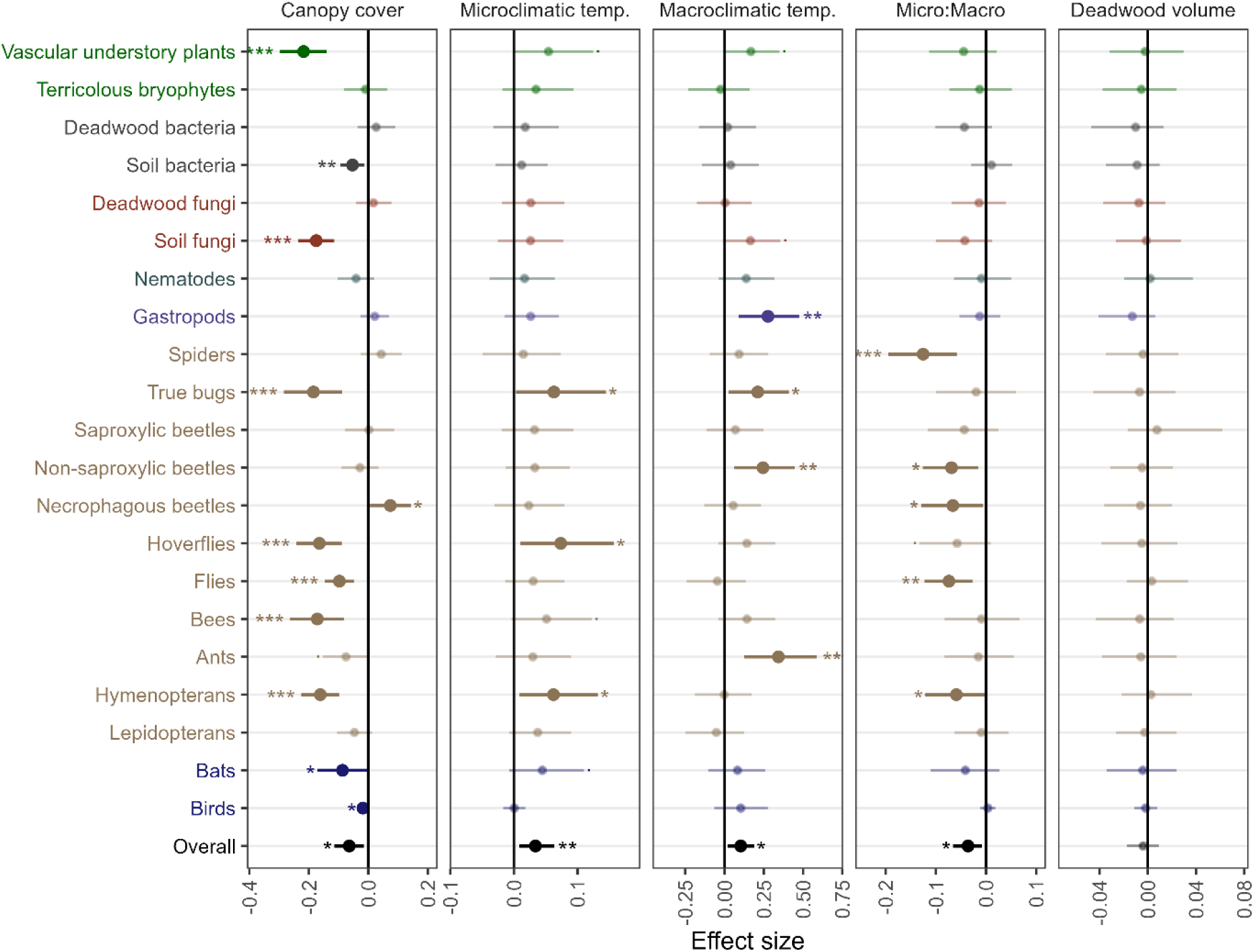
Results for the taxonomic groups. Shown are effect sizes from the model (posterior mean ± 95% credible intervals) for q = 1 from the conditional estimates for all 21 organism groups, as well as the overall effects from the marginal estimates of the model (in black). The effect sizes are given for canopy cover, microclimatic temperature, macroclimatic temperature, the interaction between micro-and macroclimatic temperature (with negative effect sizes meaning that effects of microclimate are less positive under warmer macroclimates), and deadwood volume. Colors highlight different classes of organisms (plants, bacteria, fungi, nematodes, gastropods, arthropods, vertebrates), symbols indicate parameter estimates credibly different from 0 on the 0.1, 0.05, 0.01 and 0.001 level (., *, **, and ***, respectively).

## Discussion

We generally found that lower canopy cover, i.e. more open forest stands, favored higher diversity. This was true for multi-taxon diversity and for nearly half of the studied taxonomic groups. The highly synchronized response of many different taxonomic groups to canopy structure was unanticipated, since other studies found more heterogeneous responses of different taxonomic groups to forest structural features^14,43–46^. For multi-taxon diversity, as well as diversity of true bugs, hoverflies, and hymenopterans (for *q* = 0 also understory vascular plants, bryophytes, non-saproxylic beetles, and bees), this effect of canopy cover was partially mediated by microclimatic daily maximum temperatures in summer. As we hypothesized, most of the taxonomic groups where this mediation was found are mobile ectothermic animals. Ectotherms rely on keeping their body temperature within an optimal range through thermoregulatory behavior, with severe impairment of their performance when their body temperature deviates too much from the optimum^55,56^. Likely, the higher daytime temperatures in the gaps met the thermal niches of many species in these groups better than the surrounding forest, increasing the diversity in gaps possibly by addition of more thermophilic species to the community. In addition to the summer daytime temperatures we tested, temperatures during other times of the day and year also depend on canopy cover in forests^57^, which could have driven diversity responses to canopy cover. For example, nighttime temperatures can also affect arthropod communities^58^, and insect populations respond to weather during different phases of their life cycles in complex ways^59^. While such temperature-related variables are typically highly correlated along canopy cover gradients, which makes it statistically impossible to separate their effects within this study, much evidence points towards temperature maxima being mechanistically relevant for individual-and population-level processes driving the diversity of arthropod communities^60,61^. However, testing how average thermal niches of forest-dwelling communities shift along gradients of microclimatic buffering, and whether diversity changes are driven by thermophilic or psychrophilic, and diurnal or nocturnal species, remains a research priority.

Beyond temperature effects, many other factors can potentially mediate canopy structure effects on biodiversity. Understory plant diversity increased in more open stands most likely because of the increased light availability^62–64^. Also for insects, light availability can be a direct driver of habitat use^65^. Several taxonomic groups that were more diverse in open than in closed forests feature a high number of species that are trophically directly dependent on plants, e.g. folivores (true bugs) or pollinators (hoverflies, bees). It is likely that these groups positively responded to increased understory plant diversity under more open canopies by bottom-up trophic effects, i.e. more plant species sustaining more specialized herbivores^25,66,67^. Likewise, the more developed understory vegetation in the gaps^64,68,69^ possibly also contributed to the increased diversity of these groups via increased resource availability and habitat complexity, independent of species interaction effects^14,70,71^. Remarkably, soil fungi and bacteria were also more diverse under lower canopy cover, suggesting that increased plant diversity could have altered belowground communities through plant-microbe interactions or more indirect trophic pathways^72,73^. Other contributing factors to the responses of plants and belowground organisms could be altered soil moisture regimes and soil chemical properties due to accelerated mineralization in canopy gaps^74,75^. Furthermore, canopy gaps feature higher horizontal microclimatic heterogeneity than closed stands^24^, which, following the habitat heterogeneity hypothesis^76^, could have additionally contributed to the observed responses to canopy cover. At the upper end of trophic cascades, bats and birds potentially profited from increased prey availability in open stands, in addition to direct structural effects^20,21,77^.

Necrophagous beetles stand out as the only group negatively affected in their diversity by more open canopies. This effect could either directly result from many necrophagous beetle species being more cold-adapted and therefore filtered out of communities in canopy gaps^78,79^, or from faster resource degradation of carrion in warmer daytime microclimates, similar to dung degradation effects on dung beetle communities in canopy gaps^80–82^. Since canopy cover did not affect carrion removal rates in several studies^83,84^, the former option appears more plausible.

The response of diversity to microclimatic temperatures depended on the macroclimate. Multi-taxon diversity, and the diversity of hymenopterans, flies, non-saproxylic and necrophagous beetles, and spiders, increased substantially in response to locally higher temperatures in regions with colder summers, but only slightly or not at all in regions with warmer summers. This implies that the thermal landscape context modulates how communities respond to locally warmer habitats. Due to our approach of calculating microclimatic temperatures as the deviation from a regional mean, for a given microclimatic temperature value, the absolute temperature in a warmer region is higher than for the same microclimatic temperature value in a colder region. We therefore assume that in colder regions, many species find conditions that match their thermal optimum best in canopy gaps, while in warmer regions, gaps heat up so much during summer days that heat stress becomes limiting^85,86^. Furthermore, habitats matching the niches of more thermophilic species are likely widely available in warmer regions, but rare in colder regions^87^. Similar interactions of micro-and macroclimatic temperature effects on biodiversity were recently found in a mountain landscape^47–50^, and for saproxylic beetles in boreal forests^51^, but are otherwise understudied. Remarkably, though, certain ecosystem functions in the same set of plots as this study, i.e. dung removal rates, folivory, and leaf asymmetry, also responded differently to microclimate, depending on the macroclimate^82,88^. Future studies exploring whether macroclimate-dependent reactions of biodiversity and ecosystem functions to microclimate are a common phenomenon could improve our mechanistic understanding of ecological patterns.

Diversity, not even of typically saproxylic taxonomic groups, did not depend on total deadwood volume, which seemingly contradicts well-established evidence that deadwood availability is paramount for forest biodiversity^16–18^. Similar findings of weak or absent deadwood effects are known from other studies^14,37,39^. In our case, diversity of deadwood types^89^, saproxylic community development over time along different decay stages of deadwood^90^, spatial scale in combination with high mobility of saproxylic organisms^91^, or deadwood availability levels in the surrounding landscapes^18^ might have blurred potential treatment effects. Therefore, the lack of deadwood effects in this study does not allow the conclusion that deadwood does not matter for forest biodiversity conservation.

Our findings hold key implications for the management of temperate forests. From a biodiversity conservation perspective, they imply that the most beneficial canopy structure will depend on regional temperatures, calling for regionally adapted management strategies. In regions with warm summers (mean summer temperature > 17.4°C, estimated based on the climate in our study regions, Table S7), promoting microclimatic buffering in broad-leaved forests by managing towards stands with dense canopies, i.e. through single-tree selection harvest in uneven-aged management systems, could be a valid strategy that allows avoiding negative effects of canopy gaps on tree vitality^92^ without affecting multi-taxon diversity. In colder regions (mean summer temperature < 15.6°C), however, silvicultural practices that increase the frequency of local canopy openings, e.g. gap cuts and irregular shelterwood systems^5,93^, will likely provide valuable habitats for many taxa, which allows prioritizing biodiversity conservation in regions less threatened by high temperatures. However, due to climate change, it is likely that many regions that currently feature a colder macroclimate will most probably change towards warmer macroclimates, where canopy openings may become less beneficial or even detrimental to forest biodiversity^43,94^. Furthermore, while early-and late-successional stages are underrepresented in forest landscapes managed by the continuous cover approach when no large-scale disturbances occur^42^, this situation may change under intensifying disturbance regimes in the upcoming decades^95^. If so, it will likely become unnecessary to provide additional human-made canopy openings.

Diversity indices can per definition only provide a limited perspective on the effects of forest structure and microclimate on forest-dwelling communities, since they inform us how many, but not which species were sampled. Community turnover, e.g. thermophilization in the context of climate warming, can take place without net changes in diversity^31,96–98^. Additional analyses of community composition shifts and of functional traits, thermal niches, habitat associations, and conservation relevance of the sampled species are beyond the scope of this study, but should be a research priority in the future. Particularly, it needs to be studied whether biodiversity increases in gaps are driven primarily by forest specialists, or by species associated with non-forest habitats, generalists, and disturbance indicators. While in some studies, diversity increases under more open canopies were partly driven by generalists and open-land species^37,99^, other studies found such patterns were mostly driven by characteristic species of early-successional forest stages and forest edge habitats^12,36,100^. For understory plants in our experiment, diversity increases were driven both by forest specialists and generalists^64^. For arthropods, another study from our experiment found a low proportion of forest specialists both in gaps and in closed stands^58^. Still, for many taxonomic groups, this remains a research gap. Nevertheless, even though α-diversity is higher in more open forests, deep-shade specialists still require highly buffered forest habitats^9,22,33,82^, which should be accounted for in the context of landscape-scale heterogeneity^12,52^. Future studies on the interplay of open and closed forests and its impact on β-and γ-diversity in different macroclimatic contexts should transfer the insights gained here on the α-diversity level to the landscape scale^101–104^.

In conclusion, we showed that a) many taxonomic groups are on average more diverse in more open forest stands, b) these effects are partly mediated by microclimatic temperature maxima, and c) positive effects of warmer microclimates are stronger in regions with colder summers. These findings imply that conservation-oriented forest management can increase multi-taxon diversity by providing warmer habitats in the form of canopy gaps in regions with colder summers, while the same measures will likely not affect biodiversity positively in regions with warmer summers. Moreover, our results illustrate that management schemes that have been demonstrated to have a beneficial effect on biodiversity under the current climate may no longer be effective after climate-change-driven rises in macroclimatic temperature.

### Online methods

#### Study regions

The study was conducted in the BETA-FOR experiment^52^, which comprises nine study regions distributed over Germany (Extended Data Fig. 7, Supplementary Tables S7, S8). The regions range from lowland to submontane altitudes (41 m – 1072 m a.s.l.) and cover a continentality gradient, leading to a range in mean annual temperature (1991-2010) from 5.73°C to 10.2°C.

In each region, the plots are located within mature broad-leaved forest stands representing typical production forest structures. The tree species composition ranges from European beech (*Fagus sylvatica*)-dominated stands to beech forests with high admixed proportions of common ash (*Fraxinus excelsior*), common hornbeam (*Carpinus betulus*), maple (*Acer* sp.) and oak (*Quercus* sp.).

#### Experimental design

The research was conducted in 234 permanent plots of 0.25 ha (50 m × 50 m). The plots were grouped into districts (10-20 ha), with each district containing nine plots, and each region featuring two districts. The only exception is the region University Forest, which features six districts, each of which contains 15 plots. In half of the districts (within each region, and over the whole dataset), experimental silvicultural interventions took place between winter 2015/16 and winter 2018/19 (Supplementary Table S9), with the goal to enhance between-plot structural heterogeneity^52^. The treatments combined tree removals that altered the canopy structure with deadwood retention. The tree removals were applied to ca. 30 % of the stand basal area and were executed either regularly distributed over the whole plot, and thereby resembling single-tree selection-based thinning operations, or aggregated around the plot center, resulting in canopy gaps with diameters of approximately 30 m. Deadwood from the harvested trees was retained as standing structures, lying structures, none, or both (Supplementary Table S10). Due to the structural variation of plots with the same treatments, we describe forest structure with continuous variables (canopy cover and deadwood volume) instead of comparing the treatments with each other.

#### Predictors

Canopy cover was measured during the vegetation periods of 2023, 2024, and 2025 using ground-based LiDAR. A hand-held mobile laser scanning (MLS) device of the type ZEB Horizon (GeoSLAM Ltd., Nottingham, UK) was carried through the plots in outward spirals starting from the plot center. The resulting point clouds were processed using the Faro Connect software (Faro Technologies Inc., Lake Marry, USA), and in subsequent steps noise filtered and normalized with regard to terrain. Then, we voxelized the point clouds to a resolution of 20 cm and calculated canopy cover in percent as the fraction of plot area vertically covered by canopy, considering all structures above 5 m as canopy^24^.

Both micro-and macroclimatic temperatures were derived from in-situ measurements of air temperatures at 2 m height in the plot centers. At each plot, we installed one EL-USB-2 temperature logger (Lascar Electronics, Salisbury, UK) within a TX Cover sun shield (Technoline, Bernburg, Germany) on a pole and measured air temperatures in half-hourly intervals during June to August 2023. For details on the temperature data processing, see Note S1. The diurnal maximum was chosen over other summary statistics due to its known biological relevance^60,61^. We summarized the data to the average daily maximum temperatures over this period. To separate within-from across-region signal (i.e., micro-vs. macroclimatic temperatures), we calculated region means of all plots with a canopy cover of at least 95 %, which we used as macroclimatic temperatures. To obtain microclimatic temperatures, we centered the temperatures around the macroclimatic region means (Fig. 2).

The total volume of coarse woody debris (deadwood), including stumps, snags, logs and crown deadwood with a diameter > 7 cm, was inventoried in all patches after the interventions.

#### Biodiversity sampling

We sampled biodiversity data for 21 taxonomic groups (Fig. 1). Understory vascular plants, terricolous bryophytes, nematodes and necrophagous beetles were sampled in 2023 in all regions. All other groups were sampled in 2022 in six regions, and in 2023 in the remaining three regions. The sampling methods ranged from vegetation surveys for plants over pitfall traps, flight interception traps, pan traps, light traps, soil cores and malaise traps for invertebrates, deadwood and soil samples for fungi and bacteria, to acoustic monitoring for bats and birds (Fig. 1). All details on the sampling campaigns are given in Note S2. All raw biodiversity datasets are published (Table S11). Data were generally aggregated to the plot level by pooling observations from all sampling units (e.g., traps) and sampling periods per plot whenever applicable. The term “taxonomic groups” is used throughout the paper for the sake of simplicity and adherence to established terminology, even though it is not entirely correct in a strict sense. The groups are not entirely mutually exclusive (e.g., we sampled Hymenopterans with malaise traps, but also bees with pan traps) and not only defined through taxonomic classification, but also sampling methods and ecological strategies (e.g., by distinguishing between saproxylic, non-saproxylic, and necrophagous beetles, or deadwood-inhabiting and soil-inhabiting microbial communities).

#### Biodiversity indices

Biodiversity was quantified as (coverage-standardized) taxonomic α Hill numbers of orders *q* = 0, 1, and 2 (no weighing, moderate weighing, and strong weighing by abundance or incidence frequency)^105^. These indices are equivalent to species richness (*q* = 0), the exponential of the Shannon Index (*q* = 1), and the inverse of the Simpson Index (*q* = 2). Coverage standardization is a technique to account for differences in sample completeness (coverage) by estimating the observed coverage of samples, and then standardizing all samples that are compared to each other to the same coverage, either by extra-or interpolating their estimated diversity using rarefaction curves^106^. This can either be done based on abundance data (number of individuals sampled, or pseudo-abundances such as the number of reads during metabarcoding) or incidence frequency data (presence/absence of species in several samples from the same assemblage). We calculated coverage-standardized Hill numbers for most taxonomic groups using the R package `iNEXT.3D` v. 1.0.11^53^ and standardized to group-specific coverages, following the default setting in `iNEXT.3D` that is oriented on the assemblage with the lowest observed coverage to avoid extreme extrapolation. See Supplementary Table S12 for group-specific information on the use of abundance, pseudo-abundance or incidence frequency data, and coverages to which we standardized the estimated Hill numbers.

For groups where taxa were identified via metabarcoding (fungi, bacteria, and insects from Malaise traps), the observational units were not species, but operational taxonomic units (OTUs). Data processing followed established workflows^39,107^. For each taxonomic group, we constructed a plot-by-OTU read matrix. To avoid the bias of an inflated number of singletons (i.e., single reads) in metabarcoding data, which hampers coverage estimation, we estimated the number of true singletons by applying an extended approach of Chiu & Chao, 2016^108^. This approach infers the expected number of true singletons based on the relationship between singleton frequencies and the observed frequencies of doubletons, tripletons, and quadrupletons. Surplus singletons were randomly removed from the dataset^39,107^. The number of reads during metabarcoding of the singleton-filtered matrices were used as pseudo-abundance data to estimate coverage-standardized OTU diversity with `iNEXT.3D`.

For understory vascular plants and terricolous bryophytes, we did not apply coverage standardization, because their abundance was recorded as percent cover in vegetation surveys, not as number of individuals, which would make no sense for clonal organisms. Since detectability in sessile organisms like plants is not as problematic as in mobile organism groups, we assumed coverage to be sufficiently comparable between plots for understory vascular plants and terricolous bryophytes, and calculated non-standardized Hill numbers using the R package `hillR` v. 0.5.2^109^. The resulting diversity data are shown in Extended Data Fig. 8.

From these data, we calculated multi-taxon diversity following Allan et al., 2014^54^ for all three orders of *q*. First, the diversity within all groups was re-scaled to the interval from 0 (no species sampled) to 1 (diversity of the most diverse assemblage in the dataset). Then, multi-taxon diversity of each plot was calculated as the mean of these re-scaled values of all taxonomic groups (Extended Data Fig. 9).

#### Statistical modeling

All statistical modelling was performed in R v.4.5.2^110^ in a Bayesian context using R package `brms` v. 2.23.0^111^.

For all statistical models, canopy cover was transformed with a scaled logit transformation in the form *x_trans_* = *logit*(0.99*x*/100 + 0.005) to reduce the high leverage of low canopy cover values resulting from the clustering of values around 100% (cf. Extended Data Fig 1). Additionally, the transformed canopy cover, as well as micro-and macroclimatic temperature and deadwood volume were centered and scaled to facilitate prior selection and the interpretation of coefficients.

To assess whether canopy cover effects on biodiversity are mediated by microclimatic temperature, we employed a causal inference framework based on the causal diagram in Fig. 1^112^. To achieve this, first, we tested whether microclimatic temperatures depended on canopy cover using a Bayesian linear model with canopy cover as the sole predictor, using canopy cover as a variance covariate to account for the higher variance for more open canopies. Next, we tested whether multi-taxon diversity depended on canopy cover, micro-and macroclimatic temperatures as well as their interaction, and deadwood volume. This was done in three separate models for multi-taxon diversity at *q* = 0, 1, and 2. Since multi-taxon diversity is restricted to the range of 0 – 1, we modeled it in a Bayesian generalized linear mixed model following a Beta distribution with random intercepts for districts nested in regions. To test whether the data are consistent with the causal diagram in Fig. 1, we tested for directed separation by evaluating the conditional independence claims implicit in Fig. 1^23^ using Bayesian correlation tests based on a set of bivariate and multivariate normal models (Note S3). Analogous to a piecewise structural equation model, we further performed Fisher’s C test based on these conditional independence claims^23^ using approximate “Bayesian p-values” based on the probability of direction^113^. We found credible correlations between canopy cover and deadwood volume as well as between macroclimatic temperature and deadwood volume which were not accounted for in the original causal diagram (Fig. 1), which was hence updated for the final model.

Analogous to the model for multi-diversity, we tested whether diversity of the separate taxonomic groups depended on the same set of predictors. We again fitted three separate models for the different orders of *q*, but modeled all 21 taxonomic groups together in a hierarchical modelling context. These models followed the same fixed effects structure as the models for multi-taxon diversity, but with taxonomic-group level random intercepts and slopes for all predictor variables to permit differences in response between groups. In addition, the model contained random intercepts for districts nested in regions that were nested within the random species-group intercepts. To account for the distribution of Hill numbers, which are continuous and positive, have a non-constant variance-to-mean relationship, and can take values of exactly zero, we developed a custom distribution for `brms` based on the Gamma distribution that allows for flexible handling of excess zeros and non-standard variance-to-mean relationships (see Note S3). For all models, we used moderately informative priors and ran the models on 4 chains with 10000 iterations each, half of which were discarded as warmup. Model convergence was assessed by inspection of trace plots and the potential scale reduction factor R̂. To test model adequacy, we performed prior and posterior predictive checks^115^, leave-one-out cross-validation via `R` package `loo` v. 2.9.0^116,117^ and assessed residual diagnostics based on `R` package `DHARMa` v.0.4.7^117^ via `DHARMA.helpers` v.0.0.2^119^. Parameter estimates were assumed to be credibly different from zero if their 95 % credible intervals excluded zero. For more information on model structure, prior specification, model fitting and validation, see Note S3.

## Acknowledgements

We thankfully acknowledge the contributions of our cooperation partners at the Bavarian Forest National Park, Bistumswald Passau, National Park Hunsrück-Hochwald, Saarforst, Forstamt Sailershausen, and Stadtwald Lübeck. Furthermore, we are grateful for the invaluable efforts of all technicians and students involved in data acquisition. KP and RML gratefully acknowledge the kind support of BS (R.I.P.) and G. We would like to sincerely thank the DFG for funding this project.

## Author contributions

**Conceptualization**: Kerstin Pierick, Roman M. Link, Orsi Decker, Martin Ehbrecht, Nico Eisenhauer, Csenge V. Horváth, Ludwig Lettenmaier, Jörg Müller, Julia Rothacher, Michael Scherer-Lorenzen, Dominik Seidel, Bernhard Schuldt, Christian Ammer

**Methodology:** Kerstin Pierick, Roman M. Link, Anne Chao, Csenge V. Horváth, Jörg Müller, Bernhard Schuldt, Christian Ammer

**Investigation**: Kerstin Pierick, Orsi Decker, Johanna Asch, Petr Baldrian, Claus Bässler, Julian Bittermann, Vendula Brabcová, Pia M. Bradler, Medha Bulusu, Lena Carlson, Simone Cesarz, Alice Claßen, Nico Daume, Benjamin M. Delory, Sebastian Dittrich, Bronwyn Lira Dyson, Nico Eisenhauer, Heike Feldhaar, Joshua Ferenczy, Andreas Fichtner, Ronja Hausmann, Michael Junginger, Lisa Köstler-Albert, Claudia Massó Estaje, Oliver Mitesser, Jörg Müller, Goddert von Oheimb, Marcell K. Peters, Ruth Pickert, Julia Rothacher, Kai Sattler, Michael Scherer-Lorenzen, Jens Schlüter, Rike Schwarz, Ingolf Steffan-Dewenter, Jean-Léonard Stör, Leah Vogelfänger, Clara Wild

**Data curation**: Kerstin Pierick, Dominik Seidel

**Formal analysis**: Kerstin Pierick, Roman M. Link

**Visualization**: Kerstin Pierick, Roman M. Link, Medha Bulusu, Lena Carlson, Jeisson Rodriguez Valenzuela, Julia Rothacher

**Writing – original draft**: Kerstin Pierick

**Writing – review and editing**: Kerstin Pierick, Roman M. Link, Orsi Decker, Johanna Asch, Petr Baldrian, Claus Bässler, Vendula Brabcová, Pia M. Bradler, Medha Bulusu, Marc Cadotte, Lena Carlson, Simone Cesarz, Alice Claßen, Benjamin M. Delory, Sebastian Dittrich, Bronwyn Lira Dyson, Martin Ehbrecht, Nico Eisenhauer, Heike Feldhaar, Andreas Fichtner, Csenge V. Horváth, Patrick Kacic, Ludwig Lettenmaier, Claudia Massó Estaje, Oliver Mitesser, Akira Mori, Jörg Müller, Goddert von Oheimb, Marcell K. Peters, Fons van der Plas, Julia Rothacher, Michael Scherer-Lorenzen, Rike Schwarz, Ingolf Steffan-Dewenter, Leah Vogelfänger, Clara Wild, Dominik Seidel, Bernhard Schuldt, Christian Ammer

**Supervision**: Kerstin Pierick, Dominik Seidel, Bernhard Schuldt, Christian Ammer

**Funding acquisition:** Claus Bässler, Simone Cesarz, Alice Claßen, Nico Eisenhauer, Heike Feldhaar, Andreas Fichtner, Jörg Müller, Goddert von Oheimb, Marcell K. Peters, Michael Scherer-Lorenzen, Ingolf Steffan-Dewenter, Dominik Seidel, Bernhard Schuldt, Christian Ammer

All authors read and approved of the submitted version.

## Data and code availability

The data and code will be made publicly available upon acceptance of the article. The underlying raw biodiversity data are already available as separate public datasets (Table S10).

## Extended data figures

**Extended Data Figure 1:**
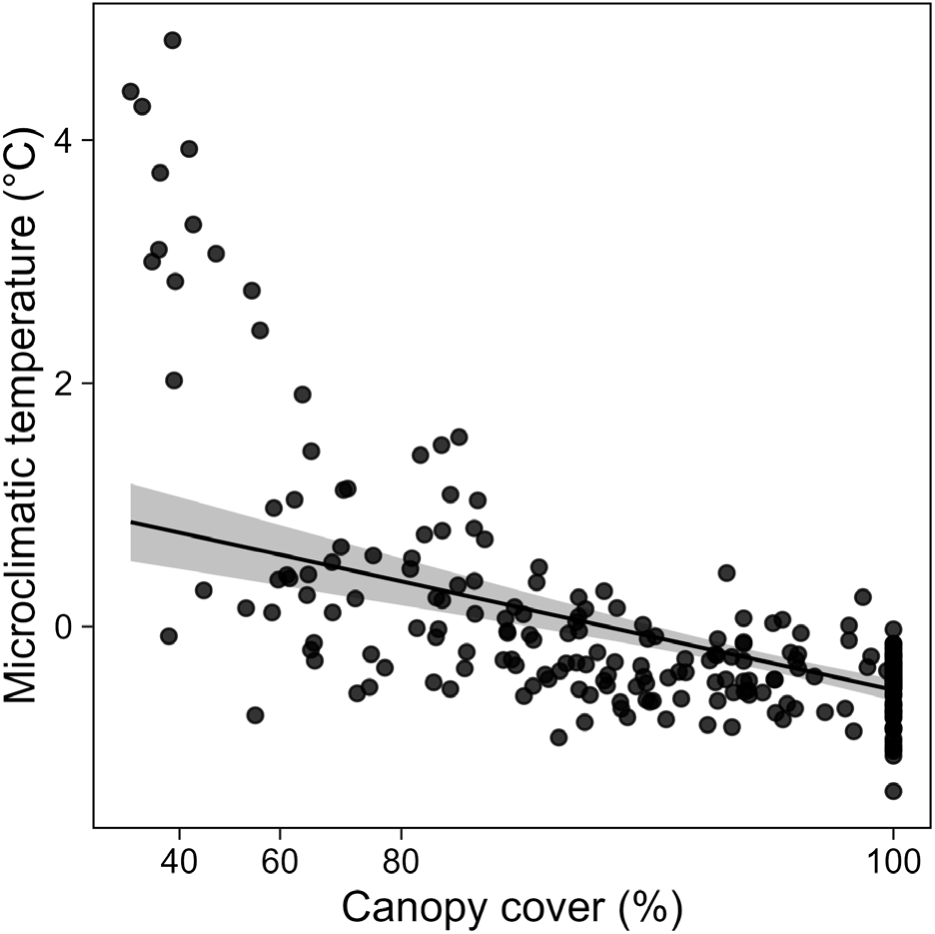
Relationship between canopy cover and microclimatic temperature. The trend line depicts predictions from a Bayesian linear regression of microclimatic temperature in dependency of canopy cover with 95% credible intervals. To account for heteroscedasticity, a regression term that lets the residual standard deviation depend on canopy cover was added.

**Extended Data Figure 2:**
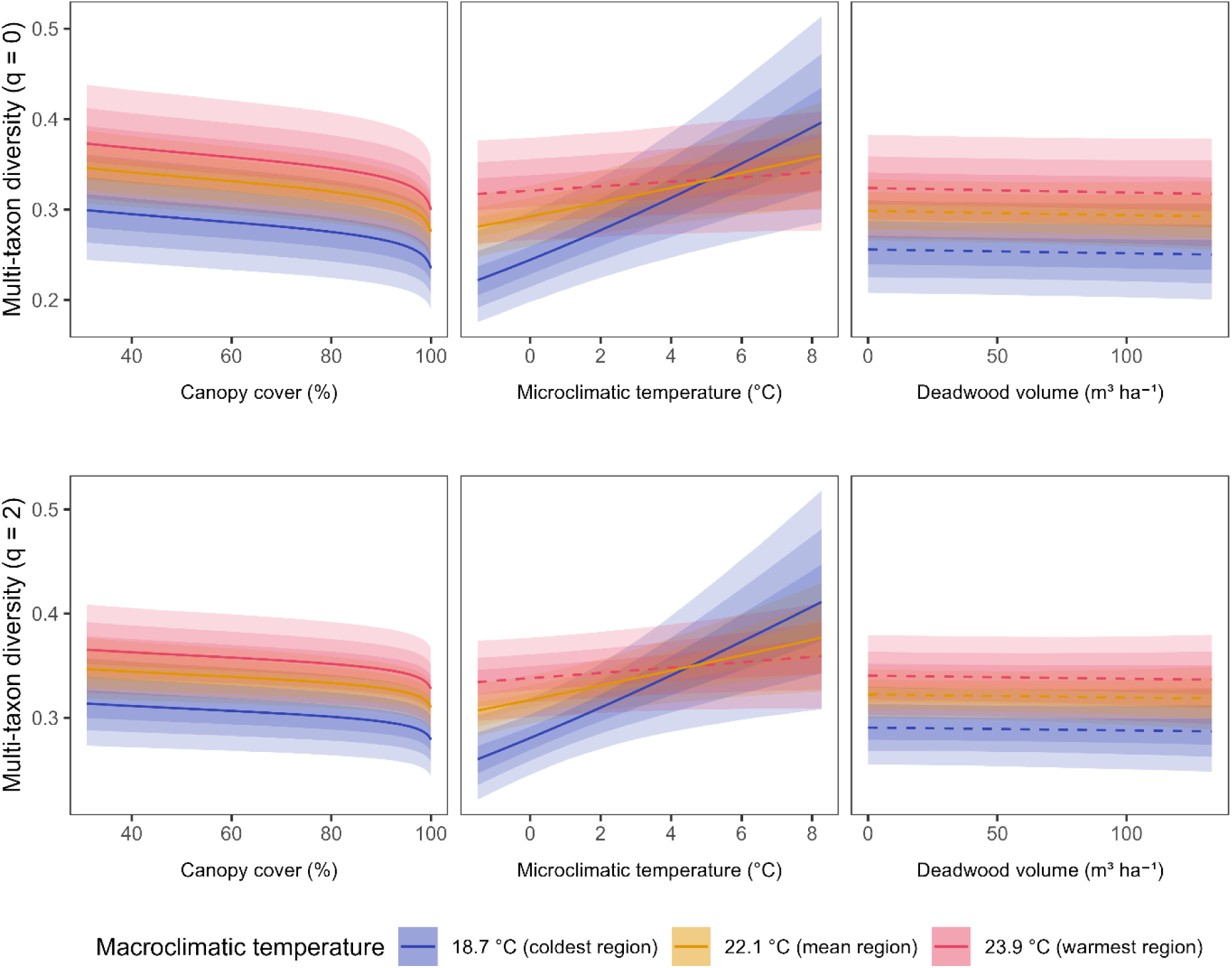
Conditional predictions for multi-taxon diversity of orders q = 0 and 2, analogous to Fig. 3a) from the main text. Shown are posterior means with their 50%, 80%, and 95% credible intervals for the coldest and warmest region in the dataset, and a hypothetical average region. Solid lines represent relationships credibly different from zero, dashed lines represent relationships that do not differ credibly from zero.

**Extended Data Figure 3:**
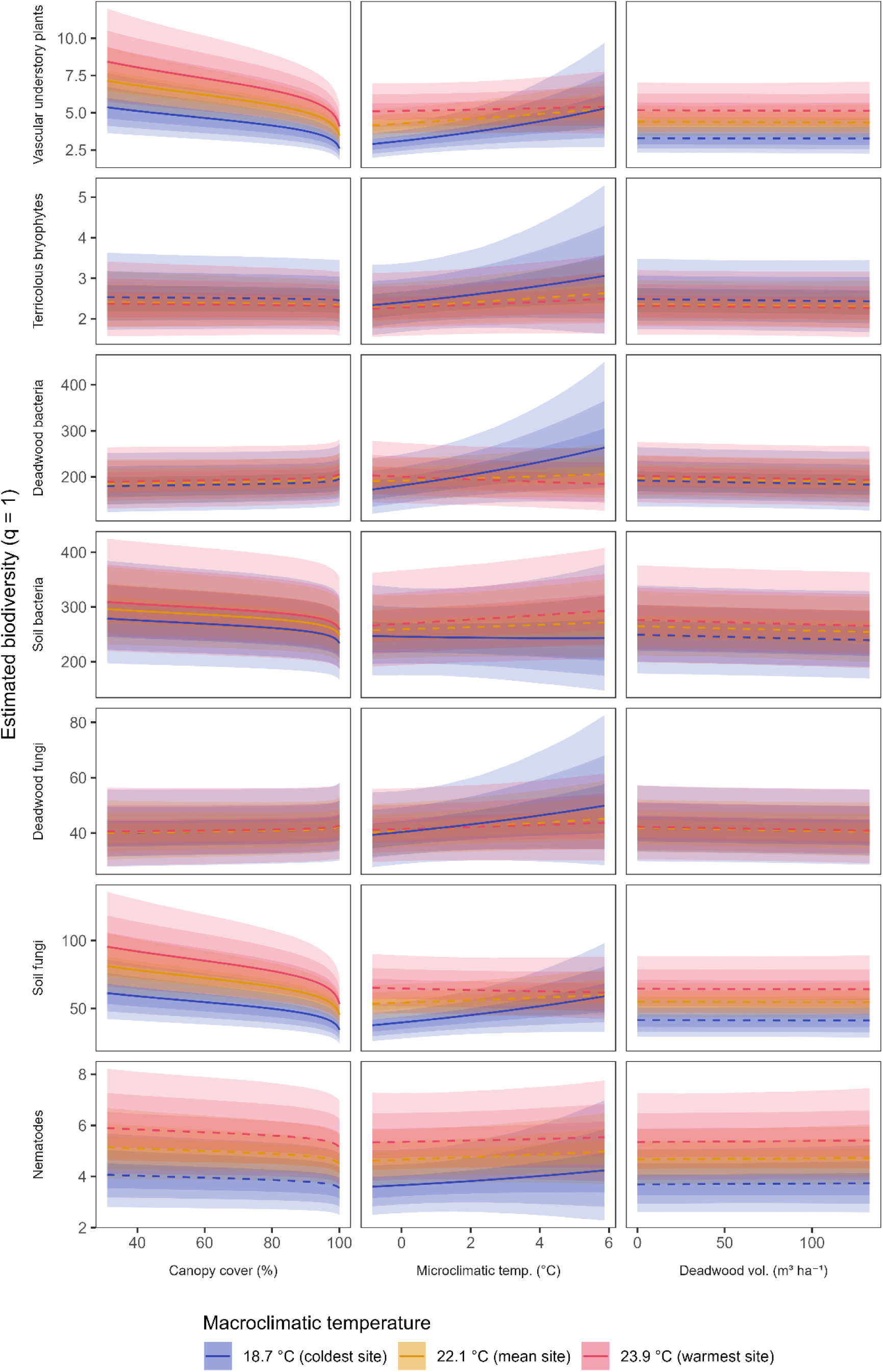
Conditional predictions for the taxonomic groups (part 1). For each analyzed species group, shown are the posterior means with their 50%, 80%, and 95% credible intervals for the coldest and warmest region in the dataset, and a hypothetical average region. Solid lines represent relationships credibly different from zero, dashed lines represent relationships that do not differ credibly from zero.

**Extended Data Figure 4:**
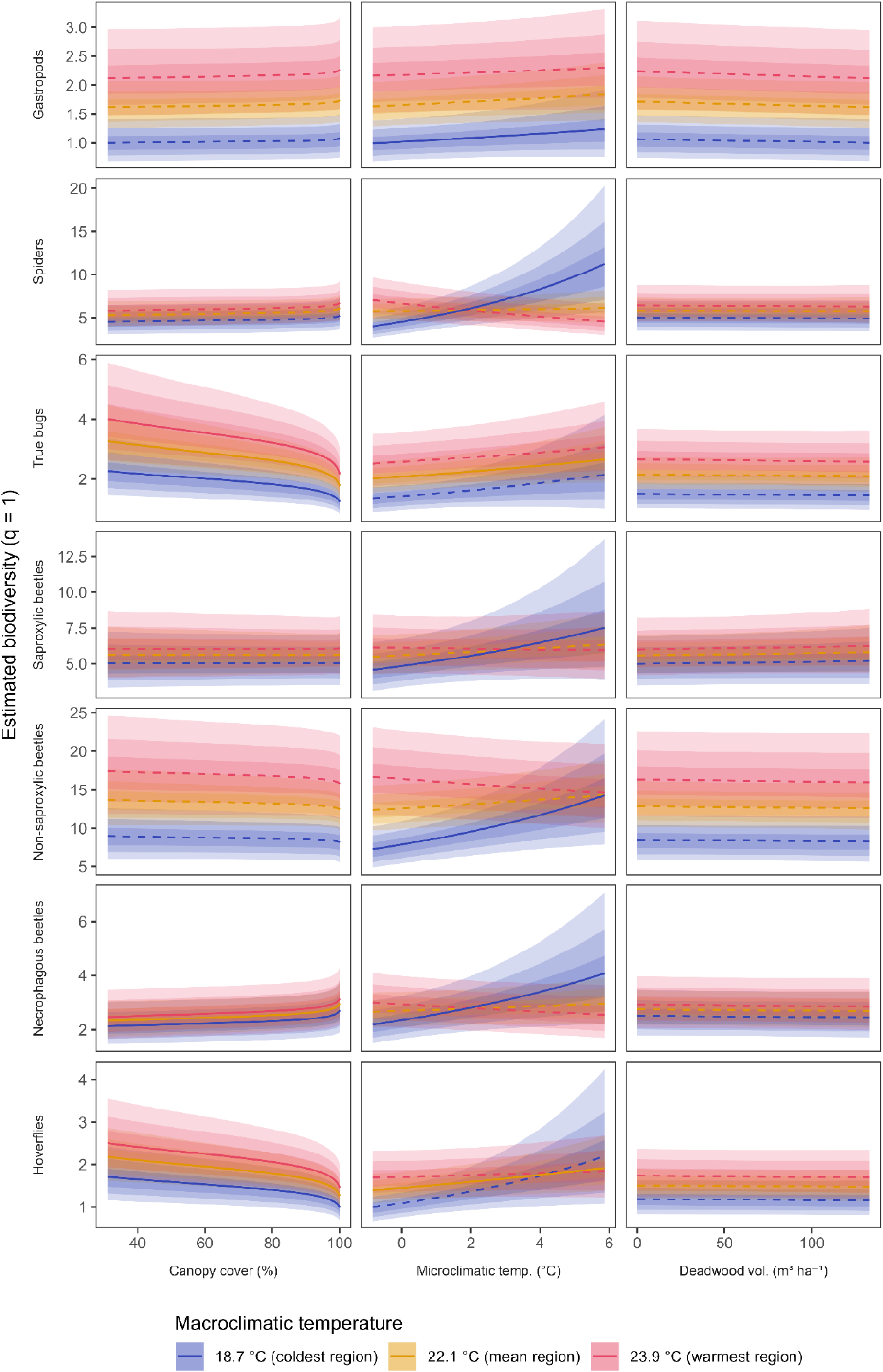
**Conditional predictions for the taxonomic groups (part 2).**

**Extended Data Figure 5:**
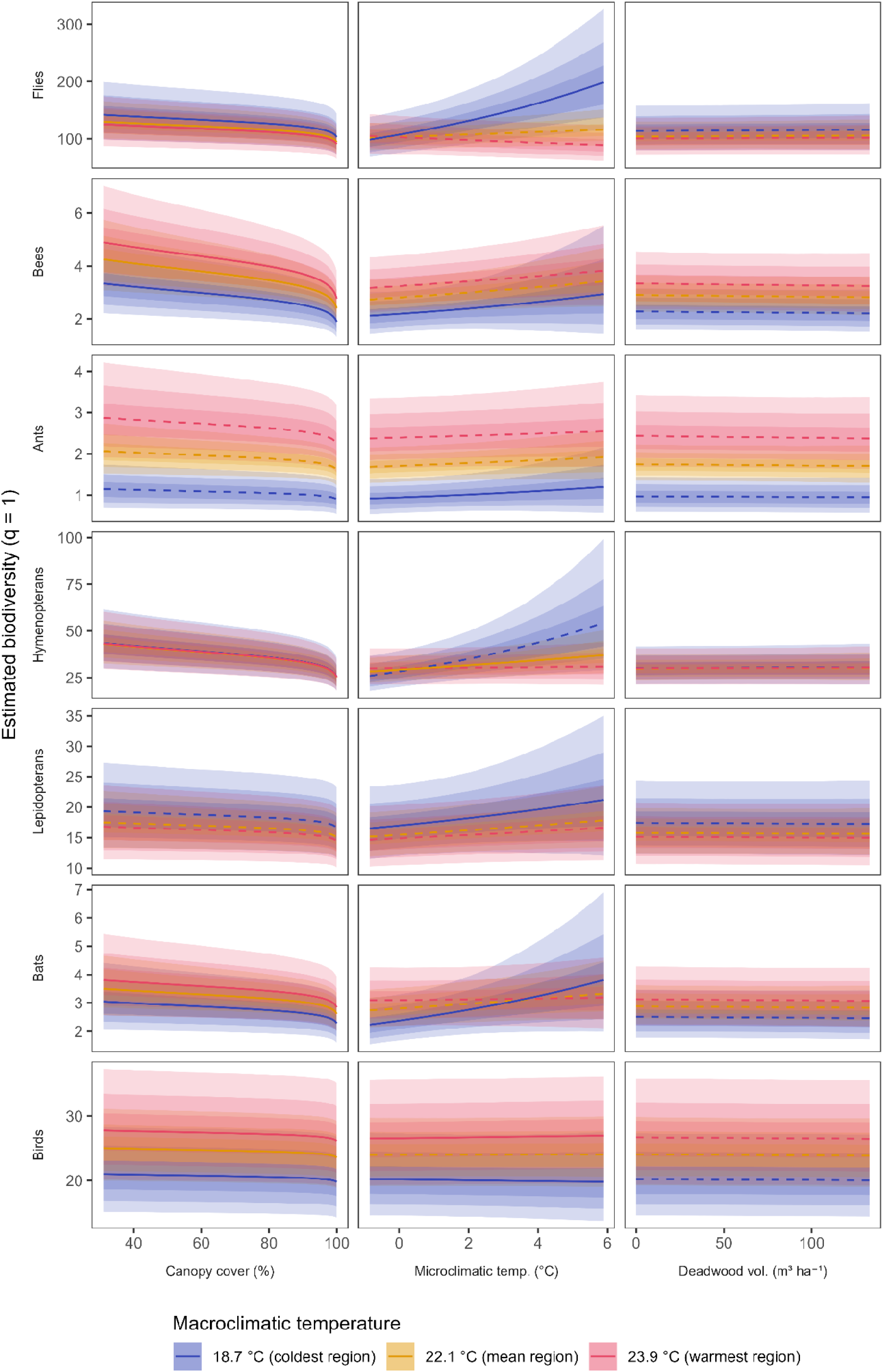
**Conditional predictions for the taxonomic groups (part 3).**

**Extended Data Figure 6:**
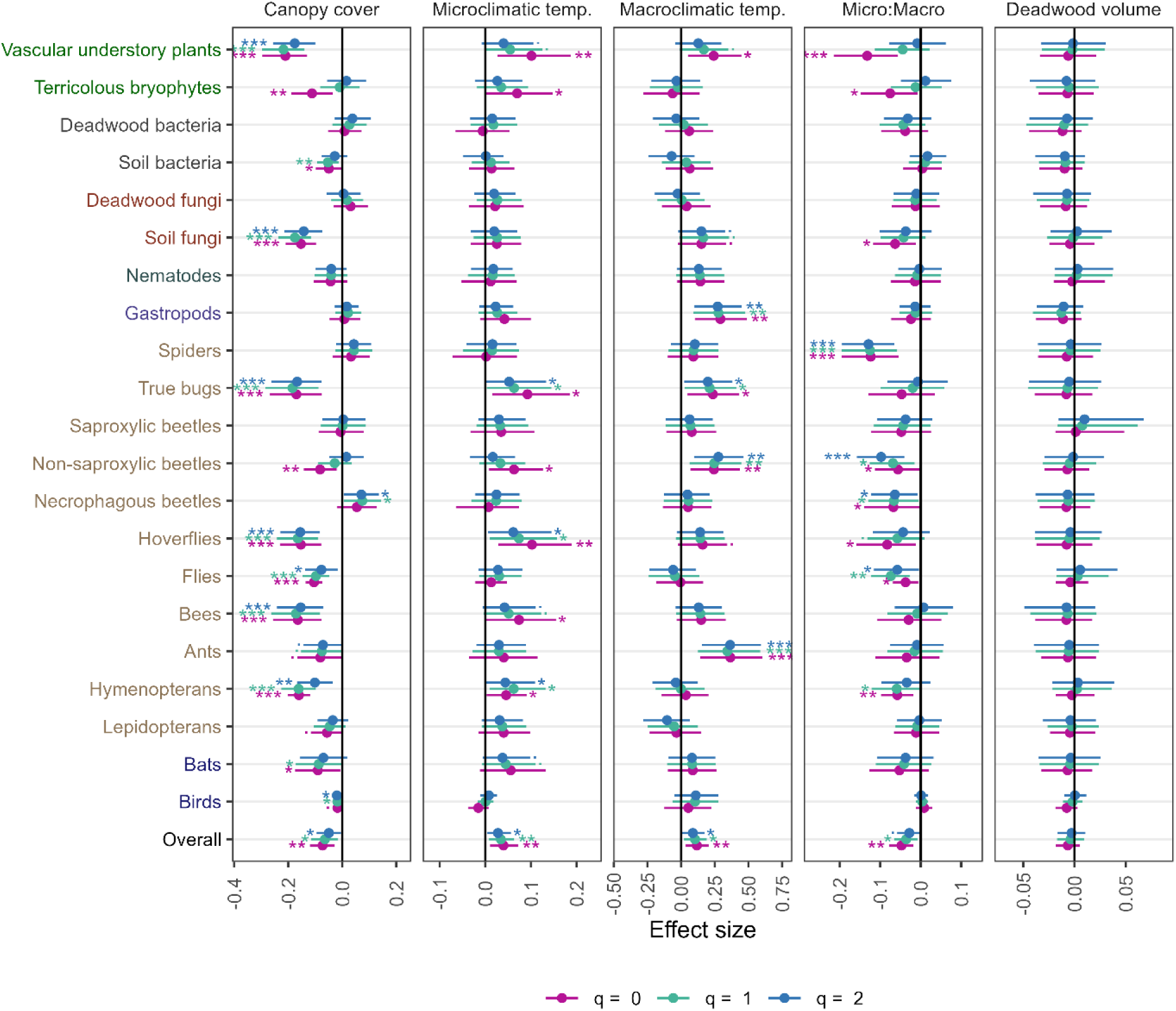
Effect sizes for all species groups for Hill numbers of order q = 0, 1, 2. Shown are estimates from the model (posterior mean ± 95% credible intervals) from the conditional estimates for all 21 taxonomic groups, as well as the overall effects from the marginal estimates of the model. Colors of the y axis labels highlight different classes of organisms (plants, bacteria, fungi, nematodes, gastropods, arthropods, vertebrates), symbols indicate parameter estimates credibly different from 0 on the 0.1, 0.05, 0.01 and 0.001 level (., *, **, and ***, respectively).

**Extended Data Fig. 7:**
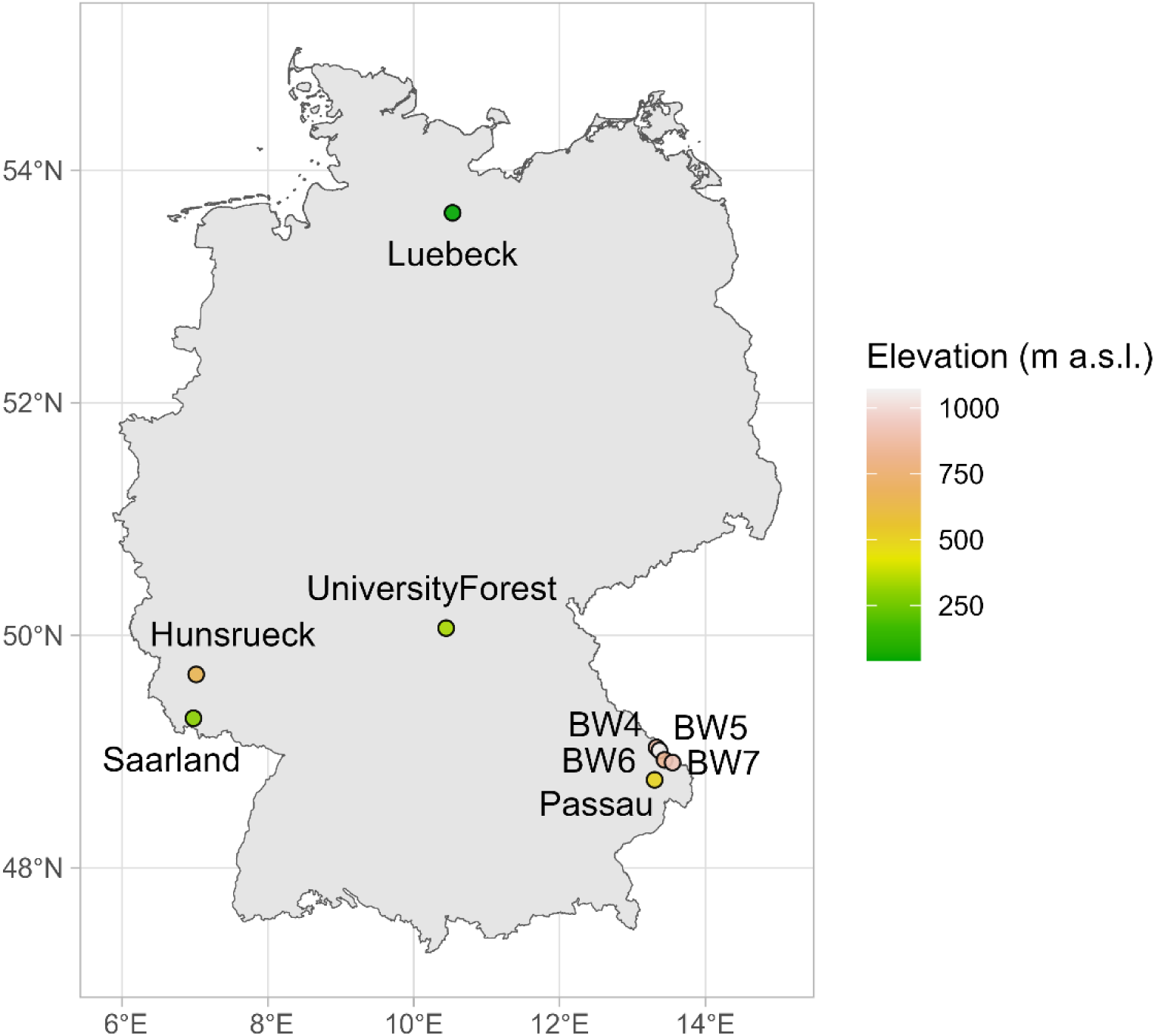
**Locations of the study regions within Germany. “BW” is short for Bayerwald.**

**Extended Data Fig. 8.**
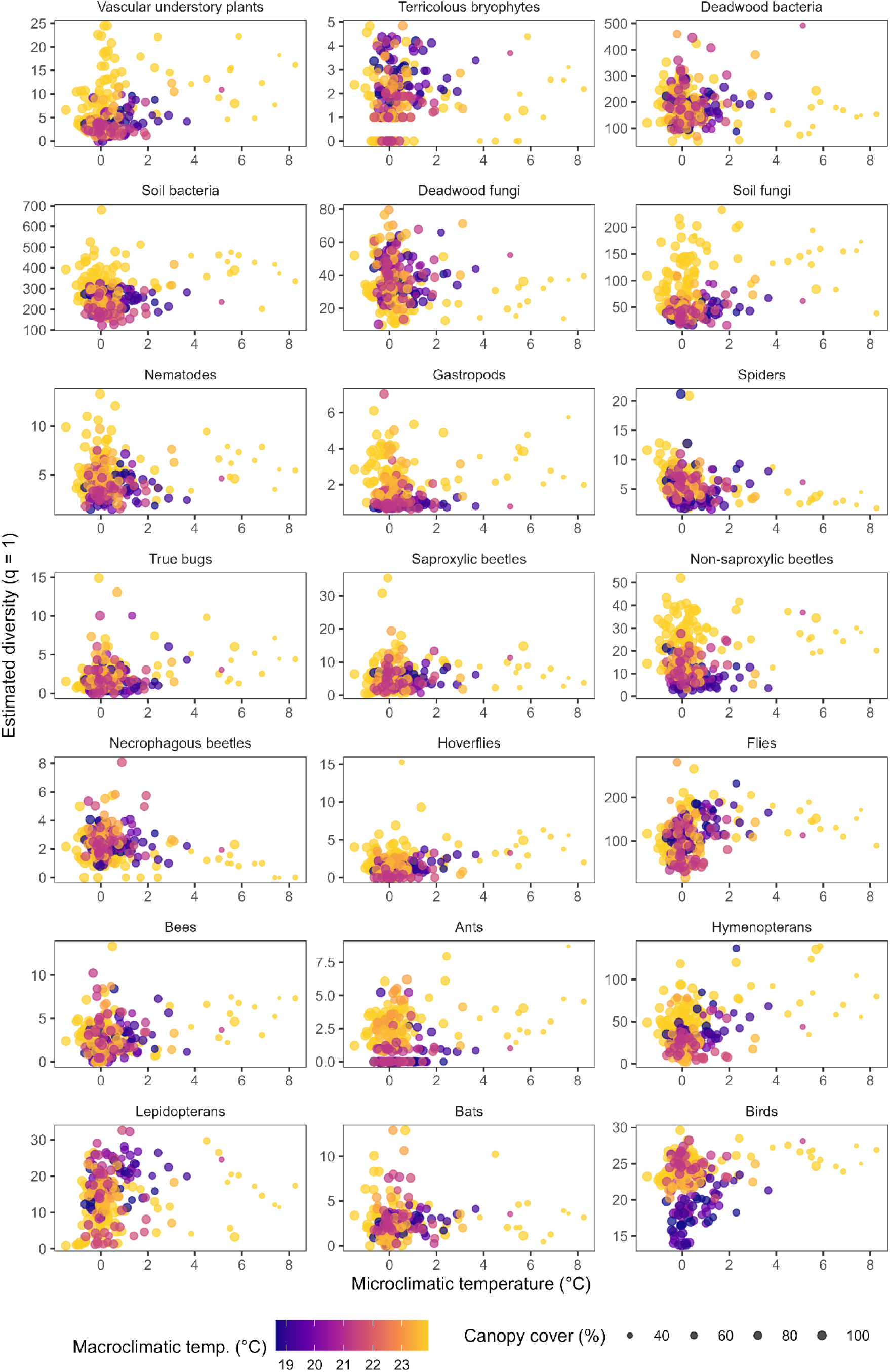
Estimated diversity (q = 1) of all taxonomic groups in dependency of microclimatic temperature, macroclimatic temperature, and canopy cover.

**Extended Data Fig. 9.**
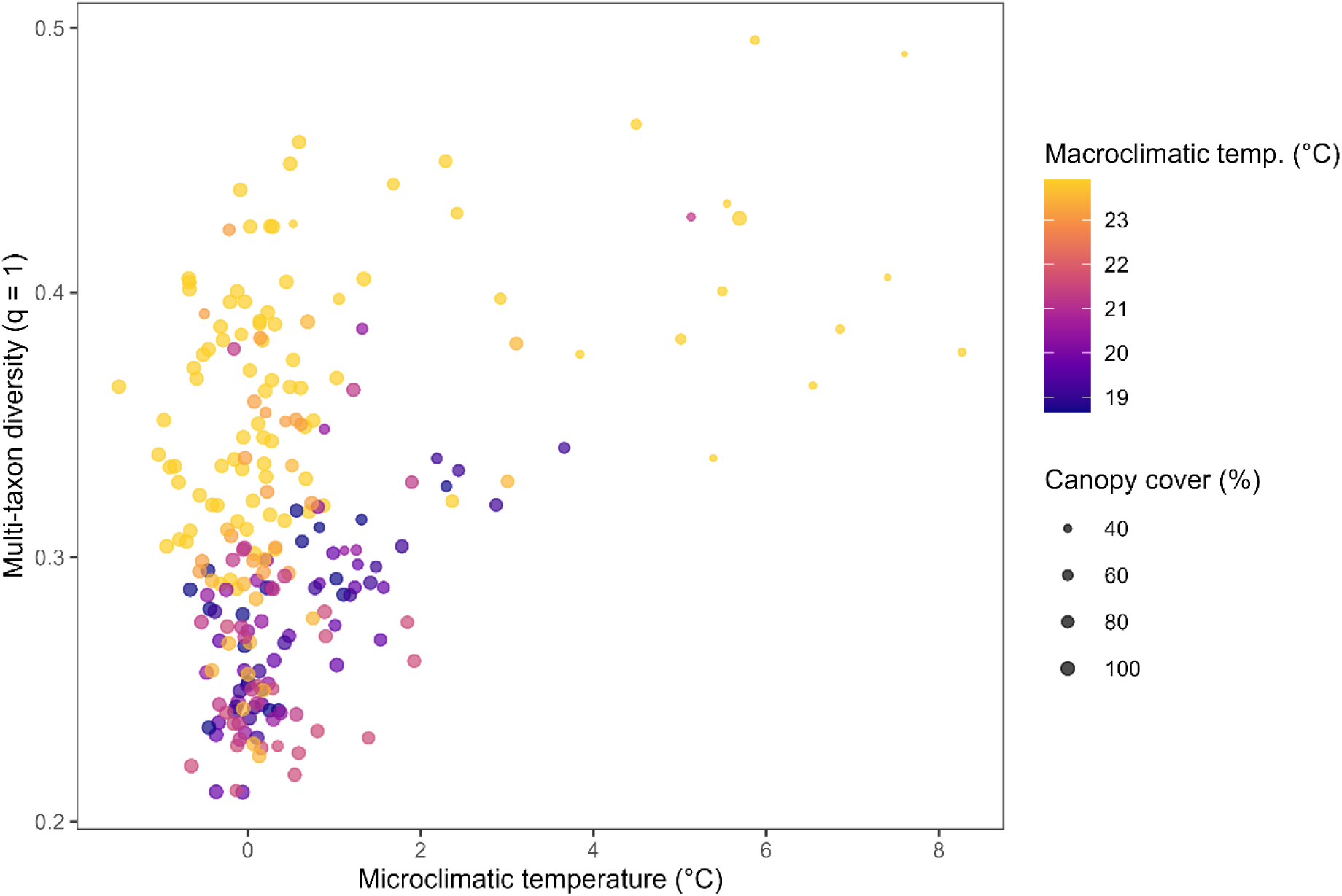
Multi-taxon diversity (q = 1) in dependency of microclimatic temperature, macroclimatic temperature, and canopy cover.

## Supplementary material

**Table S1:** Inference from the model of microclimatic temperature in dependency of canopy cover. For all parameters, the estimated value and its error, lower and upper 95% credible intervals, as well as the R^ metric and bulk and tail effective sample sizes are given. The parameters are intercept and slope for the dependency of the predicted microclimatic temperature on canopy cover, and intercept and slope for the change of residual standard deviation σ along canopy cover.

| Parameter | Estimate | Est. Error | l-95% CI | u-95% CI | $\hat{R}$ | Bulk ESS | Tail ESS |
| --- | --- | --- | --- | --- | --- | --- | --- |
| Intercept | -0.071 | 0.048 | -0.166 | 0.023 | 1.000 | 15290 | 15645 |
| Slope | -0.418 | 0.057 | -0.527 | -0.305 | 1.001 | 13535 | 15212 |
| σ- intercept | -0.489 | 0.047 | -0.579 | -0.394 | 1.000 | 28600 | 15861 |
| σ-slope | -0.391 | 0.039 | -0.467 | -0.315 | 1.000 | 22252 | 15650 |

**Table S2:** Results of the test of mediation of the microclimate effect on multi-taxon-biodiversity. Shown are the nine regions ordered by their average macroclimatic temperature, the effect size of the total microclimate effect on biodiversity for that corresponding site (posterior mean ± 95% credible intervals), and the size of the indirect effect of canopy cover mediated by microclimatic temperature (product of canopy cover effect on microclimate and microclimate effect on biodiversity; posterior mean ± 95% credible intervals).

| Region | Macroclimatic temperature | Microclimatic effect | Microclim. eff. l-95% CI | Microclim. eff. u-95% CI | Mediation | Mediation l-95% CI | Mediation u-95% CI |
| --- | --- | --- | --- | --- | --- | --- | --- |
| Bayerwald5 | 18.7 | 0.141 | 0.048 | 0.233 | 0.059 | 0.019 | 0.103 |
| Bayerwald7 | 19.1 | 0.130 | 0.044 | 0.214 | 0.054 | 0.018 | 0.095 |
| Bayerwald4 | 19.7 | 0.115 | 0.040 | 0.190 | 0.048 | 0.016 | 0.084 |
| Bayerwald6 | 20.5 | 0.098 | 0.034 | 0.162 | 0.041 | 0.013 | 0.072 |
| Luebeck | 21.2 | 0.080 | 0.027 | 0.132 | 0.034 | 0.011 | 0.059 |
| Hunsrueck | 21.6 | 0.072 | 0.024 | 0.118 | 0.030 | 0.009 | 0.052 |
| Passau | 23.2 | 0.035 | 0.003 | 0.065 | 0.014 | 0.001 | 0.028 |
| Saarland | 23.5 | 0.026 | -0.003 | 0.056 | 0.011 | -0.001 | 0.024 |
| UniversityForest | 23.9 | 0.017 | -0.012 | 0.047 | 0.007 | -0.005 | 0.020 |

**Table S3:** Inference from the models of multi-taxon diversity of orders q = 0, 1, and 2. For all parameters, the estimated value and its error, lower and upper 95% credible intervals, as well as the R^ metric and bulk and tail effective sample sizes are given. R^2^_marginal_ and R^2^_conditional_ were computed using the performance::r2_bayes() function.

| Parameter | Estimate | Est. Error | l-95% CI | u-95% CI | $\hat{R}$ | Bulk ESS | Tail ESS |
| --- | --- | --- | --- | --- | --- | --- | --- |
| **Multi-taxon diversity $q = 0$, $R^2_{\text{marginal}} = 0.344$, $R^2_{\text{conditional}} = 0.813$** |  |  |  |  |  |  |  |
| Intercept | -0.862 | 0.081 | -1.022 | -0.701 | 1.000 | 7968 | 10544 |
| Canopy cover | -0.100 | 0.018 | -0.134 | -0.065 | 1.000 | 18372 | 16461 |
| Deadwood | -0.007 | 0.013 | -0.031 | 0.018 | 1.000 | 19548 | 15513 |
| Microclimatic temperature | 0.059 | 0.021 | 0.018 | 0.099 | 1.000 | 14759 | 15088 |
| Macroclimatic temperature | 0.124 | 0.081 | -0.039 | 0.283 | 1.000 | 7551 | 9805 |
| Micro:macroclimatic interaction | -0.050 | 0.017 | -0.083 | -0.016 | 1.001 | 17386 | 16069 |
| sd(District intercepts) | 0.076 | 0.024 | 0.038 | 0.131 | 1.000 | 6920 | 11155 |
| sd(Region intercepts) | 0.214 | 0.069 | 0.116 | 0.381 | 1.000 | 7960 | 12424 |
| Phi | 217.41 | 21.285 | 177.629 | 261.227 | 1.000 | 23961 | 14934 |
| **Multi-taxon diversity $q = 1$, $R^2_{\text{marginal}} = 0.343$, $R^2_{\text{conditional}} = 0.736$** |  |  |  |  |  |  |  |
| Intercept | -0.834 | 0.067 | -0.969 | -0.702 | 1.000 | 7653 | 10005 |
| Canopy cover | -0.073 | 0.018 | -0.108 | -0.038 | 1.001 | 17715 | 16099 |
| Deadwood | -0.004 | 0.013 | -0.029 | 0.021 | 1.000 | 20793 | 15519 |
| Microclimatic temperature | 0.060 | 0.021 | 0.020 | 0.102 | 1.000 | 14550 | 14935 |
| Macroclimatic temperature | 0.109 | 0.067 | -0.030 | 0.243 | 1.000 | 7804 | 8828 |
| Micro:macroclimatic interaction | -0.046 | 0.017 | -0.079 | -0.012 | 1.000 | 15936 | 15334 |
| sd(District intercepts) | 0.079 | 0.024 | 0.042 | 0.135 | 1.001 | 6455 | 10480 |
| sd(Region intercepts) | 0.170 | 0.059 | 0.084 | 0.309 | 1.000 | 7443 | 8779 |
| Phi | 214.793 | 21.357 | 174.561 | 258.902 | 1.000 | 22473 | 14369 |
| **Multi-taxon diversity $q = 2$, $R^2_{\text{marginal}} = 0.295$, $R^2_{\text{conditional}} = 0.602$** |  |  |  |  |  |  |  |
| Intercept | -0.747 | 0.053 | -0.853 | -0.640 | 1.000 | 10755 | 11973 |
| Canopy cover | -0.050 | 0.017 | -0.084 | -0.016 | 1.000 | 19457 | 15422 |
| Deadwood | -0.004 | 0.013 | -0.029 | 0.021 | 1.000 | 23931 | 15613 |
| Microclimatic temperature | 0.050 | 0.021 | 0.009 | 0.090 | 1.000 | 17665 | 15031 |
| Macroclimatic temperature | 0.086 | 0.053 | -0.021 | 0.192 | 1.000 | 10486 | 10761 |
| Micro:macroclimatic interaction | -0.040 | 0.017 | -0.073 | -0.006 | 1.000 | 19621 | 16012 |
| sd(District intercepts) | 0.072 | 0.024 | 0.035 | 0.128 | 1.000 | 6354 | 9815 |
| sd(Region intercepts) | 0.131 | 0.050 | 0.053 | 0.252 | 1.001 | 7164 | 6963 |
| Phi | 210.241 | 20.672 | 171.694 | 252.398 | 1.000 | 23436 | 14745 |

**Table S4:** Results from evidence ratio tests on the multi-taxon diversity models. The tests were conducted with the brms::hypothesis() function to test whether slopes for microclimatic temperature were credibly different from 0 in the coldest and warmest region in the dataset, since these contrasts could not be directly derived from the estimated parameters alone. Specifically, the tested hypotheses were microclimatic temperature + micro:macroclimatic interaction * (-1.75) = 0 for the coldest, and microclimatic temperature + micro:macroclimatic interaction * 0.94 = 0 for the warmest region, with-1.75 and 0.94 corresponding to the macroclimatic temperatures of the coldest and warmest region on the scale used in the model.

| Model | Slope | Estimate | Est. Error | l-95% CI | u-95% CI |
| --- | --- | --- | --- | --- | --- |
| Multi-taxon diversity $q = 0$ | Microclimate, coldest region | 0.146 | 0.047 | 0.053 | 0.237 |
|  | Microclimate, warmest region | 0.012 | 0.015 | -0.018 | 0.041 |
| Multi-taxon diversity $q = 1$ | Microclimate, coldest region | 0.140 | 0.047 | 0.049 | 0.233 |
|  | Microclimate, warmest region | 0.017 | 0.015 | -0.012 | 0.047 |
| Multi-taxon diversity $q = 2$ | Microclimate, coldest region | 0.120 | 0.046 | 0.027 | 0.210 |
|  | Microclimate, warmest region | 0.013 | 0.015 | -0.016 | 0.042 |

**Table S5:**
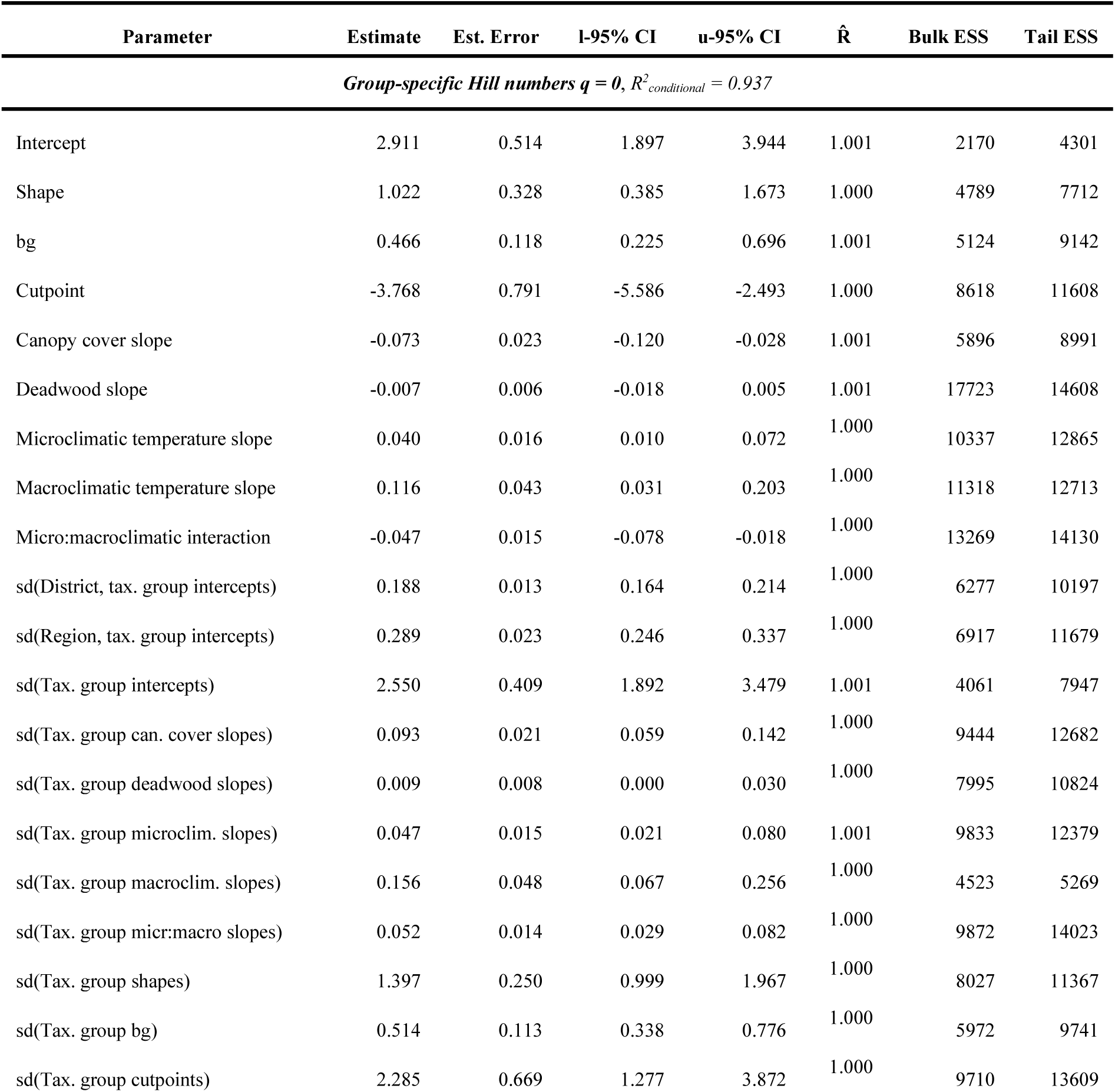
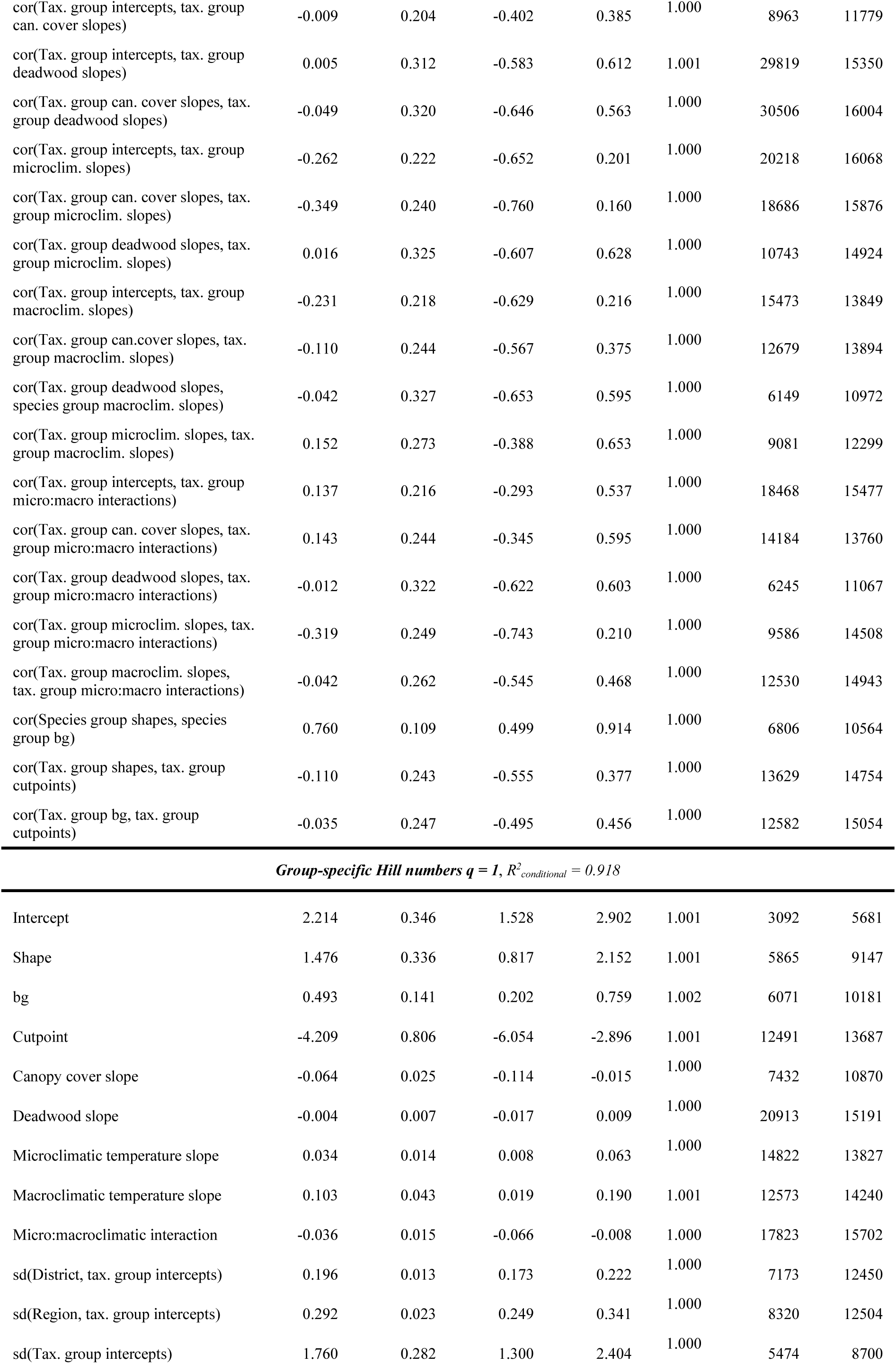
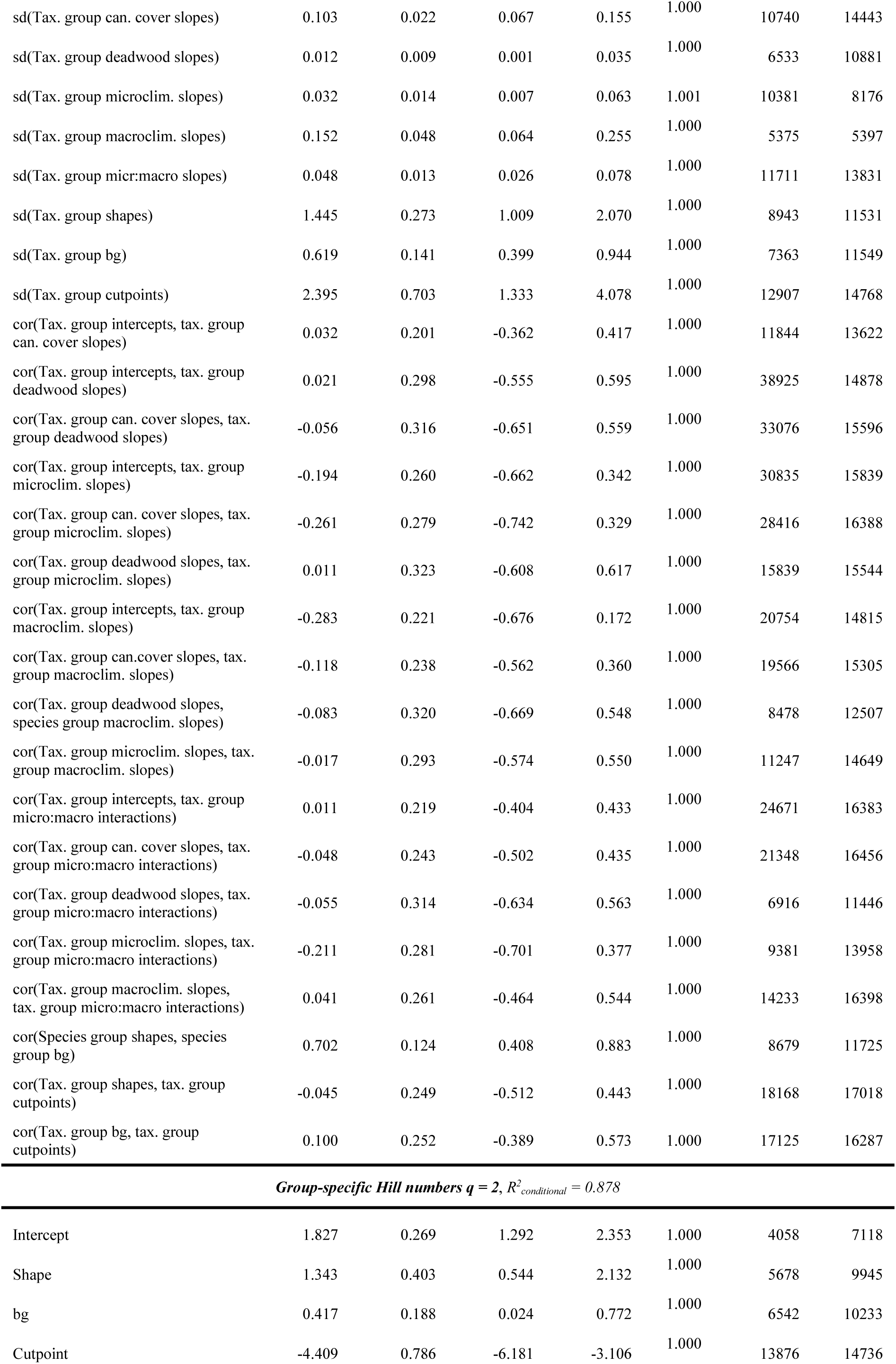
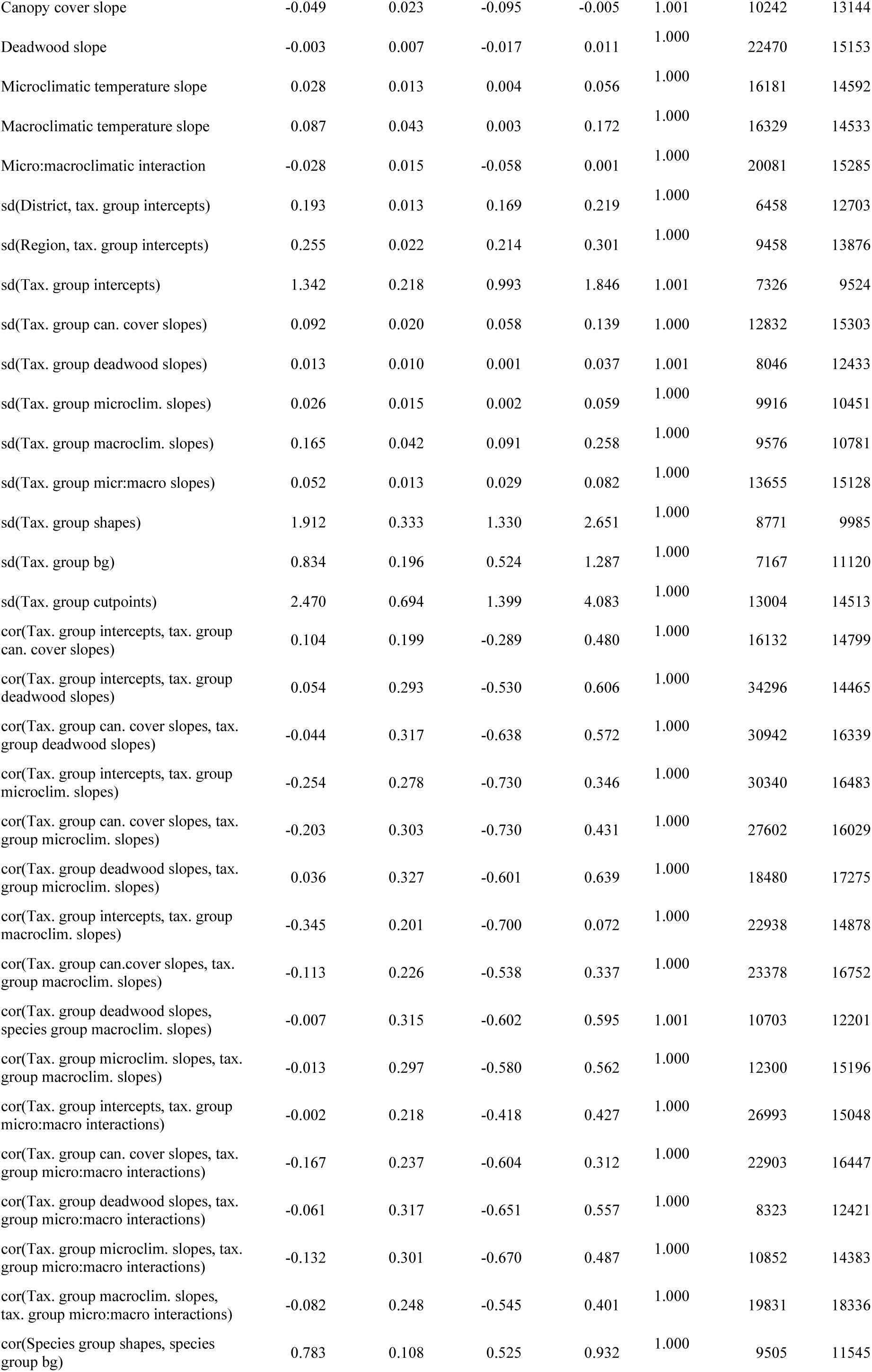
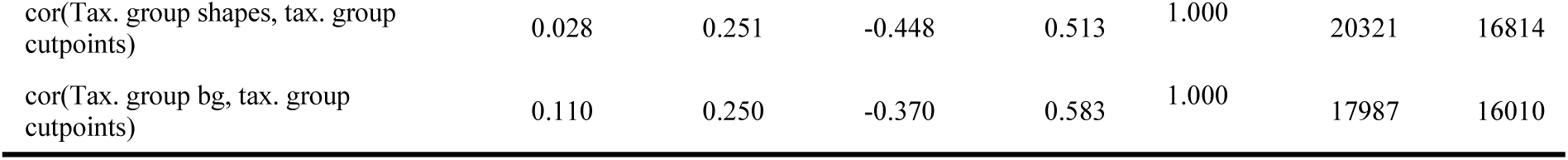
Inference from the models of taxonomic group-specific Hill numbers of orders q = 0, 1, and 2. For all parameters, the estimated value and its error, lower and upper 95% credible intervals, as well as the R^ metric and bulk and tail effective sample sizes are given. Here we only provide R^2^_conditional_, computed with the brms::bayes_R2() function, because all taxonomic group-specific effects are encoded in the random part of the model, rendering R^2^_marginal_ not meaningfully interpretable. For detailed descriptions of the parameters, see Notes S3.

| Parameter | Estimate | Est. Error | l-95% CI | u-95% CI | $\hat{R}$ | Bulk ESS | Tail ESS |
| --- | --- | --- | --- | --- | --- | --- | --- |
| *Group-specific Hill numbers $q = 0$, $R^2_{\text{conditional}} = 0.937$* |  |  |  |  |  |  |  |
| Intercept | 2.911 | 0.514 | 1.897 | 3.944 | 1.001 | 2170 | 4301 |
| Shape | 1.022 | 0.328 | 0.385 | 1.673 | 1.000 | 4789 | 7712 |
| bg | 0.466 | 0.118 | 0.225 | 0.696 | 1.001 | 5124 | 9142 |
| Cutpoint | -3.768 | 0.791 | -5.586 | -2.493 | 1.000 | 8618 | 11608 |
| Canopy cover slope | -0.073 | 0.023 | -0.120 | -0.028 | 1.001 | 5896 | 8991 |
| Deadwood slope | -0.007 | 0.006 | -0.018 | 0.005 | 1.001 | 17723 | 14608 |
| Microclimatic temperature slope | 0.040 | 0.016 | 0.010 | 0.072 | 1.000 | 10337 | 12865 |
| Macroclimatic temperature slope | 0.116 | 0.043 | 0.031 | 0.203 | 1.000 | 11318 | 12713 |
| Micro:macroclimatic interaction | -0.047 | 0.015 | -0.078 | -0.018 | 1.000 | 13269 | 14130 |
| sd(District, tax. group intercepts) | 0.188 | 0.013 | 0.164 | 0.214 | 1.000 | 6277 | 10197 |
| sd(Region, tax. group intercepts) | 0.289 | 0.023 | 0.246 | 0.337 | 1.000 | 6917 | 11679 |
| sd(Tax. group intercepts) | 2.550 | 0.409 | 1.892 | 3.479 | 1.001 | 4061 | 7947 |
| sd(Tax. group can. cover slopes) | 0.093 | 0.021 | 0.059 | 0.142 | 1.000 | 9444 | 12682 |
| sd(Tax. group deadwood slopes) | 0.009 | 0.008 | 0.000 | 0.030 | 1.000 | 7995 | 10824 |
| sd(Tax. group microclim. slopes) | 0.047 | 0.015 | 0.021 | 0.080 | 1.001 | 9833 | 12379 |
| sd(Tax. group macroclim. slopes) | 0.156 | 0.048 | 0.067 | 0.256 | 1.000 | 4523 | 5269 |
| sd(Tax. group micr:macro slopes) | 0.052 | 0.014 | 0.029 | 0.082 | 1.000 | 9872 | 14023 |
| sd(Tax. group shapes) | 1.397 | 0.250 | 0.999 | 1.967 | 1.000 | 8027 | 11367 |
| sd(Tax. group bg) | 0.514 | 0.113 | 0.338 | 0.776 | 1.000 | 5972 | 9741 |
| sd(Tax. group cutpoints) | 2.285 | 0.669 | 1.277 | 3.872 | 1.000 | 9710 | 13609 |
| cor(Tax. group intercepts, tax. group can. cover slopes) | -0.009 | 0.204 | -0.402 | 0.385 | 1.000 | 8963 | 11779 |
| cor(Tax. group intercepts, tax. group deadwood slopes) | 0.005 | 0.312 | -0.583 | 0.612 | 1.001 | 29819 | 15350 |
| cor(Tax. group can. cover slopes, tax. group deadwood slopes) | -0.049 | 0.320 | -0.646 | 0.563 | 1.000 | 30506 | 16004 |
| cor(Tax. group intercepts, tax. group microclim. slopes) | -0.262 | 0.222 | -0.652 | 0.201 | 1.000 | 20218 | 16068 |
| cor(Tax. group can. cover slopes, tax. group microclim. slopes) | -0.349 | 0.240 | -0.760 | 0.160 | 1.000 | 18686 | 15876 |
| cor(Tax. group deadwood slopes, tax. group microclim. slopes) | 0.016 | 0.325 | -0.607 | 0.628 | 1.000 | 10743 | 14924 |
| cor(Tax. group intercepts, tax. group macroclim. slopes) | -0.231 | 0.218 | -0.629 | 0.216 | 1.000 | 15473 | 13849 |
| cor(Tax. group can. cover slopes, tax. group macroclim. slopes) | -0.110 | 0.244 | -0.567 | 0.375 | 1.000 | 12679 | 13894 |
| cor(Tax. group deadwood slopes, species group macroclim. slopes) | -0.042 | 0.327 | -0.653 | 0.595 | 1.000 | 6149 | 10972 |
| cor(Tax. group microclim. slopes, tax. group macroclim. slopes) | 0.152 | 0.273 | -0.388 | 0.653 | 1.000 | 9081 | 12299 |
| cor(Tax. group intercepts, tax. group micro:macro interactions) | 0.137 | 0.216 | -0.293 | 0.537 | 1.000 | 18468 | 15477 |
| cor(Tax. group can. cover slopes, tax. group micro:macro interactions) | 0.143 | 0.244 | -0.345 | 0.595 | 1.000 | 14184 | 13760 |
| cor(Tax. group deadwood slopes, tax. group micro:macro interactions) | -0.012 | 0.322 | -0.622 | 0.603 | 1.000 | 6245 | 11067 |
| cor(Tax. group microclim. slopes, tax. group micro:macro interactions) | -0.319 | 0.249 | -0.743 | 0.210 | 1.000 | 9586 | 14508 |
| cor(Tax. group macroclim. slopes, tax. group micro:macro interactions) | -0.042 | 0.262 | -0.545 | 0.468 | 1.000 | 12530 | 14943 |
| cor(Species group shapes, species group bg) | 0.760 | 0.109 | 0.499 | 0.914 | 1.000 | 6806 | 10564 |
| cor(Tax. group shapes, tax. group cutpoints) | -0.110 | 0.243 | -0.555 | 0.377 | 1.000 | 13629 | 14754 |
| cor(Tax. group bg, tax. group cutpoints) | -0.035 | 0.247 | -0.495 | 0.456 | 1.000 | 12582 | 15054 |
| *Group-specific Hill numbers $q = 1$ , $R^2_{conditional} = 0.918$* |  |  |  |  |  |  |  |
| Intercept | 2.214 | 0.346 | 1.528 | 2.902 | 1.001 | 3092 | 5681 |
| Shape | 1.476 | 0.336 | 0.817 | 2.152 | 1.001 | 5865 | 9147 |
| bg | 0.493 | 0.141 | 0.202 | 0.759 | 1.002 | 6071 | 10181 |
| Cutpoint | -4.209 | 0.806 | -6.054 | -2.896 | 1.001 | 12491 | 13687 |
| Canopy cover slope | -0.064 | 0.025 | -0.114 | -0.015 | 1.000 | 7432 | 10870 |
| Deadwood slope | -0.004 | 0.007 | -0.017 | 0.009 | 1.000 | 20913 | 15191 |
| Microclimatic temperature slope | 0.034 | 0.014 | 0.008 | 0.063 | 1.000 | 14822 | 13827 |
| Macroclimatic temperature slope | 0.103 | 0.043 | 0.019 | 0.190 | 1.001 | 12573 | 14240 |
| Micro:macroclimatic interaction | -0.036 | 0.015 | -0.066 | -0.008 | 1.000 | 17823 | 15702 |
| sd(District, tax. group intercepts) | 0.196 | 0.013 | 0.173 | 0.222 | 1.000 | 7173 | 12450 |
| sd(Region, tax. group intercepts) | 0.292 | 0.023 | 0.249 | 0.341 | 1.000 | 8320 | 12504 |
| sd(Tax. group intercepts) | 1.760 | 0.282 | 1.300 | 2.404 | 1.000 | 5474 | 8700 |
| sd(Tax. group can. cover slopes) | 0.103 | 0.022 | 0.067 | 0.155 | 1.000 | 10740 | 14443 |
| sd(Tax. group deadwood slopes) | 0.012 | 0.009 | 0.001 | 0.035 | 1.000 | 6533 | 10881 |
| sd(Tax. group microclim. slopes) | 0.032 | 0.014 | 0.007 | 0.063 | 1.001 | 10381 | 8176 |
| sd(Tax. group macroclim. slopes) | 0.152 | 0.048 | 0.064 | 0.255 | 1.000 | 5375 | 5397 |
| sd(Tax. group micr:macro slopes) | 0.048 | 0.013 | 0.026 | 0.078 | 1.000 | 11711 | 13831 |
| sd(Tax. group shapes) | 1.445 | 0.273 | 1.009 | 2.070 | 1.000 | 8943 | 11531 |
| sd(Tax. group bg) | 0.619 | 0.141 | 0.399 | 0.944 | 1.000 | 7363 | 11549 |
| sd(Tax. group cutpoints) | 2.395 | 0.703 | 1.333 | 4.078 | 1.000 | 12907 | 14768 |
| cor(Tax. group intercepts, tax. group can. cover slopes) | 0.032 | 0.201 | -0.362 | 0.417 | 1.000 | 11844 | 13622 |
| cor(Tax. group intercepts, tax. group deadwood slopes) | 0.021 | 0.298 | -0.555 | 0.595 | 1.000 | 38925 | 14878 |
| cor(Tax. group can. cover slopes, tax. group deadwood slopes) | -0.056 | 0.316 | -0.651 | 0.559 | 1.000 | 33076 | 15596 |
| cor(Tax. group intercepts, tax. group microclim. slopes) | -0.194 | 0.260 | -0.662 | 0.342 | 1.000 | 30835 | 15839 |
| cor(Tax. group can. cover slopes, tax. group microclim. slopes) | -0.261 | 0.279 | -0.742 | 0.329 | 1.000 | 28416 | 16388 |
| cor(Tax. group deadwood slopes, tax. group microclim. slopes) | 0.011 | 0.323 | -0.608 | 0.617 | 1.000 | 15839 | 15544 |
| cor(Tax. group intercepts, tax. group macroclim. slopes) | -0.283 | 0.221 | -0.676 | 0.172 | 1.000 | 20754 | 14815 |
| cor(Tax. group can.cover slopes, tax. group macroclim. slopes) | -0.118 | 0.238 | -0.562 | 0.360 | 1.000 | 19566 | 15305 |
| cor(Tax. group deadwood slopes, species group macroclim. slopes) | -0.083 | 0.320 | -0.669 | 0.548 | 1.000 | 8478 | 12507 |
| cor(Tax. group microclim. slopes, tax. group macroclim. slopes) | -0.017 | 0.293 | -0.574 | 0.550 | 1.000 | 11247 | 14649 |
| cor(Tax. group intercepts, tax. group micro:macro interactions) | 0.011 | 0.219 | -0.404 | 0.433 | 1.000 | 24671 | 16383 |
| cor(Tax. group can. cover slopes, tax. group micro:macro interactions) | -0.048 | 0.243 | -0.502 | 0.435 | 1.000 | 21348 | 16456 |
| cor(Tax. group deadwood slopes, tax. group micro:macro interactions) | -0.055 | 0.314 | -0.634 | 0.563 | 1.000 | 6916 | 11446 |
| cor(Tax. group microclim. slopes, tax. group micro:macro interactions) | -0.211 | 0.281 | -0.701 | 0.377 | 1.000 | 9381 | 13958 |
| cor(Tax. group macroclim. slopes, tax. group micro:macro interactions) | 0.041 | 0.261 | -0.464 | 0.544 | 1.000 | 14233 | 16398 |
| cor(Species group shapes, species group bg) | 0.702 | 0.124 | 0.408 | 0.883 | 1.000 | 8679 | 11725 |
| cor(Tax. group shapes, tax. group cutpoints) | -0.045 | 0.249 | -0.512 | 0.443 | 1.000 | 18168 | 17018 |
| cor(Tax. group bg, tax. group cutpoints) | 0.100 | 0.252 | -0.389 | 0.573 | 1.000 | 17125 | 16287 |
| *Group-specific Hill numbers $q = 2$ , $R^2_{conditional} = 0.878$* |  |  |  |  |  |  |  |
| Intercept | 1.827 | 0.269 | 1.292 | 2.353 | 1.000 | 4058 | 7118 |
| Shape | 1.343 | 0.403 | 0.544 | 2.132 | 1.000 | 5678 | 9945 |
| bg | 0.417 | 0.188 | 0.024 | 0.772 | 1.000 | 6542 | 10233 |
| Cutpoint | -4.409 | 0.786 | -6.181 | -3.106 | 1.000 | 13876 | 14736 |
| Canopy cover slope | -0.049 | 0.023 | -0.095 | -0.005 | 1.001 | 10242 | 13144 |
| Deadwood slope | -0.003 | 0.007 | -0.017 | 0.011 | 1.000 | 22470 | 15153 |
| Microclimatic temperature slope | 0.028 | 0.013 | 0.004 | 0.056 | 1.000 | 16181 | 14592 |
| Macroclimatic temperature slope | 0.087 | 0.043 | 0.003 | 0.172 | 1.000 | 16329 | 14533 |
| Micro:macroclimatic interaction | -0.028 | 0.015 | -0.058 | 0.001 | 1.000 | 20081 | 15285 |
| sd(District, tax. group intercepts) | 0.193 | 0.013 | 0.169 | 0.219 | 1.000 | 6458 | 12703 |
| sd(Region, tax. group intercepts) | 0.255 | 0.022 | 0.214 | 0.301 | 1.000 | 9458 | 13876 |
| sd(Tax. group intercepts) | 1.342 | 0.218 | 0.993 | 1.846 | 1.001 | 7326 | 9524 |
| sd(Tax. group can. cover slopes) | 0.092 | 0.020 | 0.058 | 0.139 | 1.000 | 12832 | 15303 |
| sd(Tax. group deadwood slopes) | 0.013 | 0.010 | 0.001 | 0.037 | 1.001 | 8046 | 12433 |
| sd(Tax. group microclim. slopes) | 0.026 | 0.015 | 0.002 | 0.059 | 1.000 | 9916 | 10451 |
| sd(Tax. group macroclim. slopes) | 0.165 | 0.042 | 0.091 | 0.258 | 1.000 | 9576 | 10781 |
| sd(Tax. group micr:macro slopes) | 0.052 | 0.013 | 0.029 | 0.082 | 1.000 | 13655 | 15128 |
| sd(Tax. group shapes) | 1.912 | 0.333 | 1.330 | 2.651 | 1.000 | 8771 | 9985 |
| sd(Tax. group bg) | 0.834 | 0.196 | 0.524 | 1.287 | 1.000 | 7167 | 11120 |
| sd(Tax. group cutpoints) | 2.470 | 0.694 | 1.399 | 4.083 | 1.000 | 13004 | 14513 |
| cor(Tax. group intercepts, tax. group can. cover slopes) | 0.104 | 0.199 | -0.289 | 0.480 | 1.000 | 16132 | 14799 |
| cor(Tax. group intercepts, tax. group deadwood slopes) | 0.054 | 0.293 | -0.530 | 0.606 | 1.000 | 34296 | 14465 |
| cor(Tax. group can. cover slopes, tax. group deadwood slopes) | -0.044 | 0.317 | -0.638 | 0.572 | 1.000 | 30942 | 16339 |
| cor(Tax. group intercepts, tax. group microclim. slopes) | -0.254 | 0.278 | -0.730 | 0.346 | 1.000 | 30340 | 16483 |
| cor(Tax. group can. cover slopes, tax. group microclim. slopes) | -0.203 | 0.303 | -0.730 | 0.431 | 1.000 | 27602 | 16029 |
| cor(Tax. group deadwood slopes, tax. group microclim. slopes) | 0.036 | 0.327 | -0.601 | 0.639 | 1.000 | 18480 | 17275 |
| cor(Tax. group intercepts, tax. group macroclim. slopes) | -0.345 | 0.201 | -0.700 | 0.072 | 1.000 | 22938 | 14878 |
| cor(Tax. group can.cover slopes, tax. group macroclim. slopes) | -0.113 | 0.226 | -0.538 | 0.337 | 1.000 | 23378 | 16752 |
| cor(Tax. group deadwood slopes, species group macroclim. slopes) | -0.007 | 0.315 | -0.602 | 0.595 | 1.001 | 10703 | 12201 |
| cor(Tax. group microclim. slopes, tax. group macroclim. slopes) | -0.013 | 0.297 | -0.580 | 0.562 | 1.000 | 12300 | 15196 |
| cor(Tax. group intercepts, tax. group micro:macro interactions) | -0.002 | 0.218 | -0.418 | 0.427 | 1.000 | 26993 | 15048 |
| cor(Tax. group can. cover slopes, tax. group micro:macro interactions) | -0.167 | 0.237 | -0.604 | 0.312 | 1.000 | 22903 | 16447 |
| cor(Tax. group deadwood slopes, tax. group micro:macro interactions) | -0.061 | 0.317 | -0.651 | 0.557 | 1.000 | 8323 | 12421 |
| cor(Tax. group microclim. slopes, tax. group micro:macro interactions) | -0.132 | 0.301 | -0.670 | 0.487 | 1.000 | 10852 | 14383 |
| cor(Tax. group macroclim. slopes, tax. group micro:macro interactions) | -0.082 | 0.248 | -0.545 | 0.401 | 1.000 | 19831 | 18336 |
| cor(Species group shapes, species group bg) | 0.783 | 0.108 | 0.525 | 0.932 | 1.000 | 9505 | 11545 |
| cor(Tax. group shapes, tax. group<br>cutpoints) | 0.028 | 0.251 | -0.448 | 0.513 | 1.000 | 20321 | 16814 |
| cor(Tax. group bg, tax. group<br>cutpoints) | 0.110 | 0.250 | -0.370 | 0.583 | 1.000 | 17987 | 16010 |

**Table S6:** Results from evidence ratio tests on the taxonomic group-specific model for q = 1. The tests were conducted with the brms::hypothesis() function to test whether slopes for specific taxonomic groups were credibly different from 0, since these contrasts could not be directly derived from the main parameters. For the slopes for canopy cover, deadwood, and microclimate in the average region, we tested the hypothesis overall slope_[predictor]_ + group-specific slope_[predictor]_ = 0. For the slopes for microclimate in the coldest region, the tested hypothesis was overall microclimate slope + group-specific microclimate slope + (overall micro:macro interaction + group-specific micro:macro interaction) *-1.75 = 0, and in the warmest region identical with 0.94 instead of-1.75. These numbers correspond to the macroclimatic temperatures of the coldest and warmest region on the scale used in the model.

| Taxonomic group | Slope | Estimate | Est. Error | l-95% CI | u-95% CI |
| --- | --- | --- | --- | --- | --- |
| Vascular plants | Canopy cover | -0.218 | 0.040 | -0.297 | -0.140 |
|  | Deadwood | -0.002 | 0.014 | -0.032 | 0.030 |
|  | Microclimate, coldest region | 0.081 | 0.039 | 0.007 | 0.162 |
|  | Microclimate, average region | 0.054 | 0.032 | -0.001 | 0.125 |
|  | Microclimate, warmest region | 0.040 | 0.057 | -0.063 | 0.164 |
| Terricolous bryophytes | Canopy cover | -0.010 | 0.037 | -0.082 | 0.064 |
|  | Deadwood | -0.005 | 0.014 | -0.037 | 0.024 |
|  | Microclimate, coldest region | 0.010 | 0.041 | 0.021 | 0.183 |
|  | Microclimate, average region | 0.035 | 0.028 | -0.018 | 0.094 |
|  | Microclimate, warmest region | 0.002 | 0.052 | -0.103 | 0.107 |
| Deadwood bacteria | Canopy cover | 0.026 | 0.032 | -0.037 | 0.090 |
|  | Deadwood | -0.010 | 0.015 | -0.047 | 0.013 |
|  | Microclimate, coldest region | 0.109 | 0.044 | 0.029 | 0.200 |
|  | Microclimate, average region | 0.018 | 0.026 | -0.033 | 0.071 |
|  | Microclimate, warmest region | -0.031 | 0.051 | -0.137 | 0.069 |
| Soil bacteria | Canopy cover | -0.053 | 0.021 | -0.093 | -0.013 |
|  | Deadwood | -0.009 | 0.011 | -0.035 | 0.010 |
|  | Microclimate, coldest region | 0.113 | 0.044 | 0.033 | 0.205 |
|  | Microclimate, average region | 0.012 | 0.021 | -0.029 | 0.053 |
|  | Microclimate, warmest region | -0.042 | 0.044 | -0.133 | 0.039 |
| Deadwood fungi | Canopy cover | 0.017 | 0.030 | -0.041 | 0.078 |
|  | Deadwood | -0.007 | 0.012 | -0.037 | 0.015 |
|  | Microclimate, coldest region | 0.102 | 0.042 | 0.025 | 0.190 |
|  | Microclimate, average region | 0.026 | 0.024 | -0.019 | 0.079 |
|  | Microclimate, warmest region | -0.014 | 0.047 | -0.110 | 0.082 |
| Soil fungi | Canopy cover | -0.175 | 0.031 | -0.236 | -0.115 |
|  | Deadwood | -0.001 | 0.013 | -0.027 | 0.027 |
|  | Microclimate, coldest region | 0.102 | 0.042 | 0.025 | 0.192 |
|  | Microclimate, average region | 0.026 | 0.026 | -0.026 | 0.078 |
|  | Microclimate, warmest region | -0.015 | 0.049 | -0.120 | 0.079 |
| Nematodes | Canopy cover | -0.041 | 0.031 | -0.103 | 0.019 |
|  | Deadwood | 0.002 | 0.014 | -0.020 | 0.037 |
|  | Microclimate, coldest region | 0.110 | 0.043 | 0.033 | 0.201 |
|  | Microclimate, average region | 0.017 | 0.025 | -0.038 | 0.064 |
|  | Microclimate, warmest region | -0.033 | 0.049 | -0.146 | 0.054 |
| Gastropods | Canopy cover | 0.022 | 0.025 | -0.026 | 0.070 |
|  | Deadwood | -0.013 | 0.012 | -0.041 | 0.006 |
|  | Microclimate, coldest region | 0.102 | 0.042 | 0.026 | 0.189 |
|  | Microclimate, average region | 0.026 | 0.021 | -0.014 | 0.071 |
|  | Microclimate, warmest region | -0.015 | 0.042 | -0.101 | 0.070 |
| Spiders | Canopy cover | 0.044 | 0.035 | -0.025 | 0.112 |
|  | Deadwood | -0.004 | 0.014 | -0.035 | 0.026 |
|  | Microclimate, coldest region | 0.111 | 0.044 | 0.029 | 0.204 |
|  | Microclimate, average region | 0.015 | 0.031 | -0.049 | 0.074 |
|  | Microclimate, warmest region | -0.037 | 0.059 | -0.168 | 0.072 |
| True bugs | Canopy cover | -0.185 | 0.050 | -0.284 | -0.088 |
|  | Deadwood | -0.007 | 0.016 | -0.045 | 0.023 |
|  | Microclimate, coldest region | 0.075 | 0.039 | -0.002 | 0.155 |
|  | Microclimate, average region | 0.063 | 0.037 | 0.003 | 0.145 |
|  | Microclimate, warmest region | 0.057 | 0.065 | -0.050 | 0.201 |
| Saproxylic beetles | Canopy cover | 0.002 | 0.043 | -0.079 | 0.087 |
|  | Deadwood | 0.008 | 0.020 | -0.017 | 0.062 |
|  | Microclimate, coldest region | 0.098 | 0.041 | 0.022 | 0.183 |
|  | Microclimate, average region | 0.032 | 0.028 | -0.020 | 0.094 |
|  | Microclimate, warmest region | -0.003 | 0.052 | -0.105 | 0.108 |
| Non-saproxylic beetles | Canopy cover | -0.028 | 0.031 | -0.090 | 0.034 |
|  | Deadwood | -0.005 | 0.012 | -0.031 | 0.021 |
|  | Microclimate, coldest region | 0.097 | 0.041 | 0.022 | 0.181 |
|  | Microclimate, average region | 0.033 | 0.026 | -0.013 | 0.088 |
|  | Microclimate, warmest region | -0.001 | 0.047 | -0.092 | 0.097 |
| Necrophagous beetles | Canopy cover | 0.074 | 0.036 | 0.006 | 0.144 |
|  | Deadwood | -0.006 | 0.013 | -0.036 | 0.020 |
|  | Microclimate, coldest region | 0.104 | 0.042 | 0.028 | 0.192 |
|  | Microclimate, average region | 0.023 | 0.027 | -0.031 | 0.079 |
|  | Microclimate, warmest region | -0.020 | 0.052 | -0.126 | 0.081 |
| Hoverflies | Canopy cover | -0.165 | 0.039 | -0.242 | -0.089 |
|  | Deadwood | -0.005 | 0.015 | -0.038 | 0.025 |
|  | Microclimate, coldest region | 0.067 | 0.038 | -0.009 | 0.142 |
|  | Microclimate, average region | 0.074 | 0.038 | 0.010 | 0.157 |
|  | Microclimate, warmest region | 0.077 | 0.066 | -0.028 | 0.225 |
| Flies | Canopy cover | -0.097 | 0.025 | -0.147 | -0.048 |
|  | Deadwood | 0.003 | 0.013 | -0.018 | 0.033 |
|  | Microclimate, coldest region | 0.099 | 0.041 | 0.023 | 0.184 |
|  | Microclimate, average region | 0.030 | 0.024 | -0.014 | 0.079 |
|  | Microclimate, warmest region | -0.007 | 0.044 | -0.096 | 0.083 |
| Bees | Canopy cover | -0.171 | 0.046 | -0.263 | -0.082 |
|  | Deadwood | -0.007 | 0.015 | -0.043 | 0.022 |
|  | Microclimate, coldest region | 0.084 | 0.040 | 0.010 | 0.167 |
|  | Microclimate, average region | 0.051 | 0.032 | -0.004 | 0.122 |
|  | Microclimate, warmest region | 0.034 | 0.058 | -0.073 | 0.157 |
| Ants | Canopy cover | -0.075 | 0.039 | -0.152 | 0.000 |
|  | Deadwood | -0.006 | 0.014 | -0.038 | 0.024 |
|  | Microclimate, coldest region | 0.100 | 0.042 | 0.024 | 0.187 |
|  | Microclimate, average region | 0.030 | 0.029 | -0.029 | 0.090 |
|  | Microclimate, warmest region | -0.008 | 0.054 | -0.122 | 0.098 |
| Hymenopterans | Canopy cover | -0.162 | 0.033 | -0.225 | -0.098 |
|  | Deadwood | 0.003 | 0.014 | -0.022 | 0.037 |
|  | Microclimate, coldest region | 0.075 | 0.039 | -0.000 | 0.152 |
|  | Microclimate, average region | 0.062 | 0.032 | 0.009 | 0.132 |
|  | Microclimate, warmest region | 0.055 | 0.056 | -0.035 | 0.178 |
| Lepidopterans | Canopy cover | -0.047 | 0.030 | -0.105 | 0.013 |
|  | Deadwood | -0.002 | 0.012 | -0.026 | 0.024 |
|  | Microclimate, coldest region | 0.094 | 0.040 | 0.020 | 0.180 |
|  | Microclimate, average region | 0.037 | 0.025 | -0.008 | 0.090 |
|  | Microclimate, warmest region | 0.007 | 0.046 | -0.083 | 0.102 |
| Bats | Canopy cover | -0.087 | 0.044 | -0.172 | -0.000 |
|  | Deadwood | -0.004 | 0.014 | -0.034 | 0.024 |
|  | Microclimate, coldest region | 0.089 | 0.039 | 0.015 | 0.170 |
|  | Microclimate, average region | 0.045 | 0.030 | -0.008 | 0.110 |
|  | Microclimate, warmest region | 0.021 | 0.053 | -0.080 | 0.135 |
| Birds | Canopy cover | -0.018 | 0.007 | -0.033 | -0.003 |
|  | Deadwood | -0.002 | 0.005 | -0.011 | 0.008 |
|  | Microclimate, coldest region | 0.122 | 0.043 | 0.040 | 0.212 |
|  | Microclimate, average region | 0.001 | 0.009 | -0.017 | 0.018 |
|  | Microclimate, warmest region | -0.065 | 0.029 | -0.123 | -0.010 |

**Table S7:**
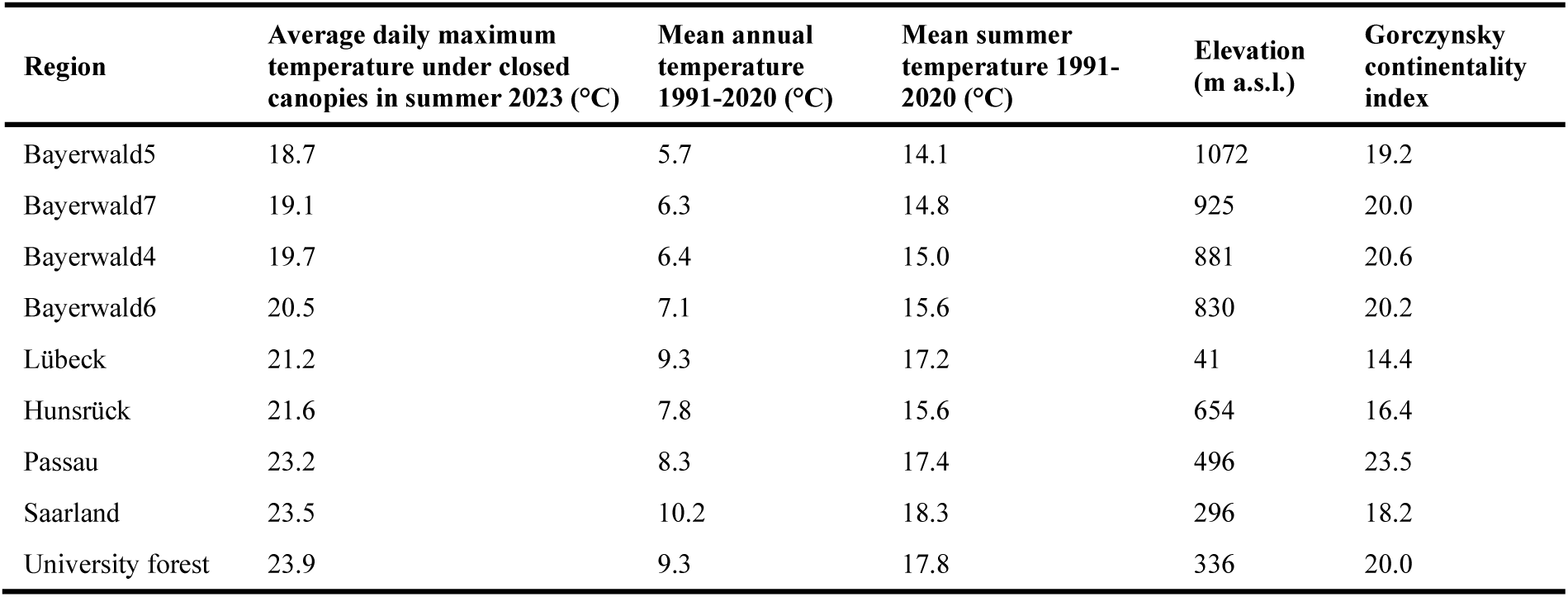
Temperature gradient of the study regions. Average daily maximum temperatures in summer 2023 are derived from in-situ measurements in the plots and are referred to as “macroclimatic temperatures” throughout the paper. Mean annual temperature and mean summer temperature for the period 1991-2020 were extracted from the “DWD Climate Data Center (CDC): Multi-annual means of grids of air temperature (2m) over Germany” dataset provided by the German Weather Service. The Gorczynsky continentality index was calculated following Szabó-Takács et al. (2015).

**Table S8:** Location and site conditions of the study regions. Longitude and latitude are given in the WGS 84 coordinate system. Soil types were determined according to Arbeitsgemeinschaft Forsteinrichtung, Arbeitskreis für Standortskartierung (2016). Tree species are given in descending order of basal area in the plots of the respective region, for all tree species contributing at least 10 % of basal area. Stand age data were provided by local forest managers.

| Site | Longitude | Latitude | Dominant soil type(s) | Tree species | Stand age (yr) |
| --- | --- | --- | --- | --- | --- |
| University forest | 10.445 | 50.062 | Parabrown earth/terra fusca | *Fraxinus excelsior*, *Fagus sylvatica*, *Acer* sp., *Carpinus betulus*, *Quercus* sp. | 64-97 |
| Bayerwald4 | 13.331 | 49.036 | Podzolic brown earth | *Fagus sylvatica* | 90 |
| Bayerwald5 | 13.371 | 49.010 | Podzolic brown earth | *Fagus sylvatica*, *Picea abies* | 80-90 |
| Bayerwald6 | 13.440 | 48.929 | Brown earth | *Fagus sylvatica* | 80 |
| Bayerwald7 | 13.551 | 48.906 | Podzolic brown earth | *Fagus sylvatica*, *Picea abies* | 90 |
| Passau | 13.305 | 48.757 | Podzolic brown earth | *Fagus sylvatica*, *Picea abies* | 70-80 |
| Saarland | 6.979 | 49.287 | Brown earth/pseudogley | *Fagus sylvatica*, *Quercus* sp. | 70-85 |
| Hunsrück | 7.017 | 49.663 | Podzolic brown earth | *Fagus sylvatica* | 125 |
| Lübeck | 10.533 | 53.634 | Podzolic brown earth/pseudogley | *Fagus sylvatica*, *Quercus* sp. | 109-127 |

**Table S9:** Timing of the interventions and measurement campaigns.

| Site(s) | Interventions | Mobile laser scanning | Temperature measurements | Plants, bryophytes, nematodes, necrophagous beetles | All other taxonomic groups |
| --- | --- | --- | --- | --- | --- |
| University forest | Winter 2018/19 | Summer 2023 | Summer 2023 | Summer 2023 | Summer 2022 |
| Bayerwald4-7, Passau | Winter 2015/16 | Summer 2024 | Summer 2023 | Summer 2023 | Summer 2022 |
| Saarland, Hunsrück, Lübeck | Winter 2016/17 | Summer 2025 | Summer 2023 | Summer 2023 | Summer 2023 |

**Table S10:**
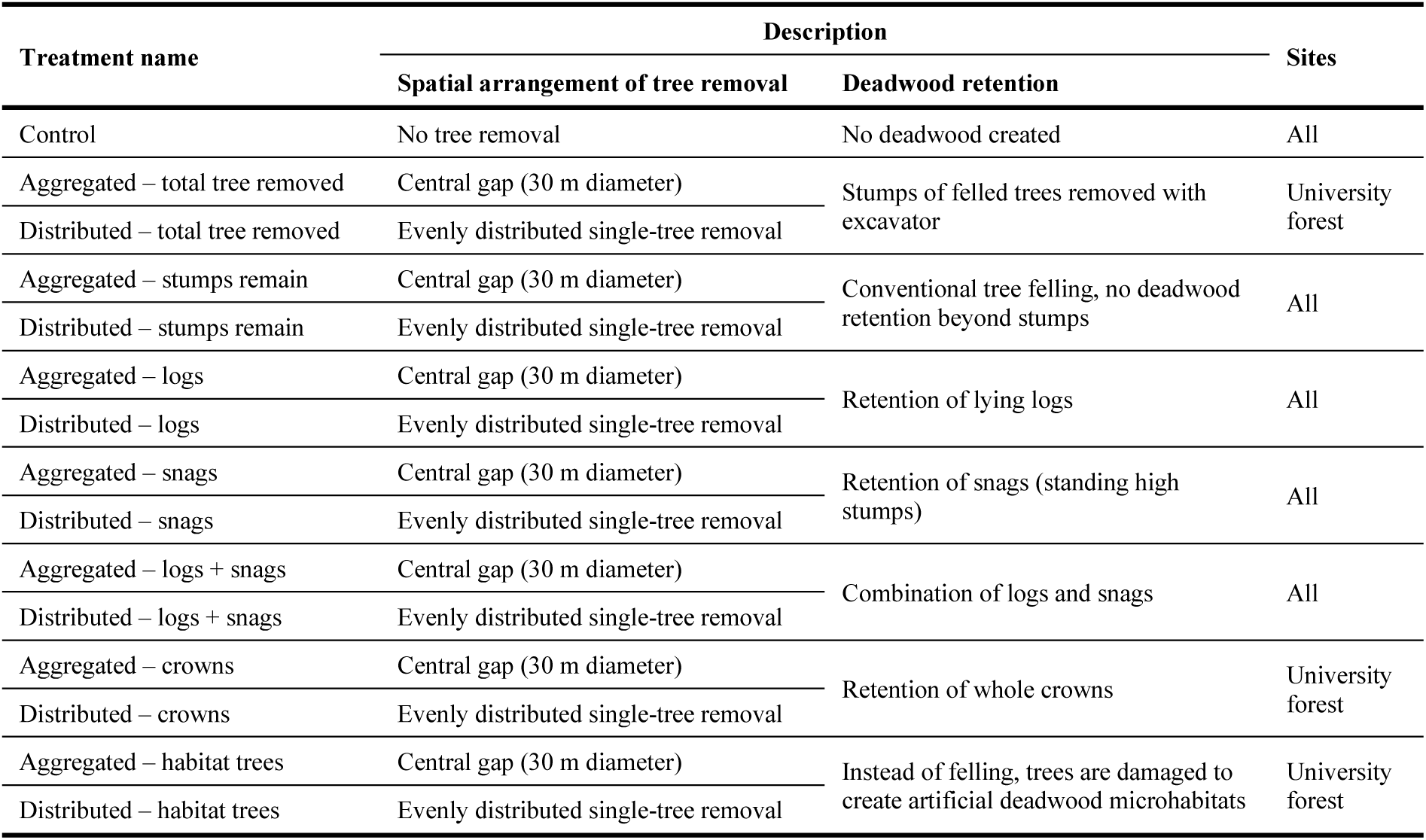
Description of the 15 experimental treatments, combining tree removal in two different spatial patterns with seven forms of deadwood retention. Three forms of deadwood retention were only applied at the University forest site. All treatments were applied to 30 % of the stand basal area.

| Treatment name | Description |  | Sites |
| --- | --- | --- | --- |
|  | Spatial arrangement of tree removal | Deadwood retention |  |
| Control | No tree removal | No deadwood created | All |
| Aggregated – total tree removed | Central gap (30 m diameter) | Stumps of felled trees removed with excavator | University forest |
| Distributed – total tree removed | Evenly distributed single-tree removal |  |  |
| Aggregated – stumps remain | Central gap (30 m diameter) | Conventional tree felling, no deadwood retention beyond stumps | All |
| Distributed – stumps remain | Evenly distributed single-tree removal |  |  |
| Aggregated – logs | Central gap (30 m diameter) | Retention of lying logs | All |
| Distributed – logs | Evenly distributed single-tree removal |  |  |
| Aggregated – snags | Central gap (30 m diameter) | Retention of snags (standing high stumps) | All |
| Distributed – snags | Evenly distributed single-tree removal |  |  |
| Aggregated – logs + snags | Central gap (30 m diameter) | Combination of logs and snags | All |
| Distributed – logs + snags | Evenly distributed single-tree removal |  |  |
| Aggregated – crowns | Central gap (30 m diameter) | Retention of whole crowns | University forest |
| Distributed – crowns | Evenly distributed single-tree removal |  |  |
| Aggregated – habitat trees | Central gap (30 m diameter) | Instead of felling, trees are damaged to create artificial deadwood microhabitats | University forest |
| Distributed – habitat trees | Evenly distributed single-tree removal |  |  |

**Table S11:** Publishing information of biodiversity raw data sets.

| **Taxonomic group** | **Dataset** | **Publication** |
| --- | --- | --- |
| Vascular understory plants | https://doi.org/10.5281/zenodo.14982724 | Bradler et al., 2026, https://doi.org/10.1111/1365-2664.70477 |
| Terricolous bryophytes | https://doi.org/10.5281/zenodo.17486171 |  |
| Deadwood bacteria | Release on SRA under ID PRJNA1435076 upon publication |  |
| Soil bacteria | https://www.ncbi.nlm.nih.gov/bioproject/?term=PRJNA1232009 |  |
| Deadwood fungi | Release on SRA under ID PRJNA1435086 upon publication |  |
| Soil fungi | https://www.ncbi.nlm.nih.gov/bioproject/PRJNA1232019 | Bässler et al., 2025, https://doi.org/10.1101/2025.08.29.673114 |
| Nematodes | https://doi.org/10.5281/zenodo.17938049 | Schwarz et al., 2026, https://doi.org/10.1016/j.soilbio.2025.110078 |
| Gastropods | https://doi.org/10.5281/zenodo.14989539 |  |
| Spiders | https://doi.org/10.5281/zenodo.14766447 | Stör et al., 2026, https://doi.org/10.1111/1365-2656.70297 |
| True bugs | https://doi.org/10.5281/zenodo.14258470 |  |
| Saproxyllic and non-saproxyllic beetles | https://doi.org/10.5281/zenodo.15396244 | Mitesser et al., 2026, https://doi.org/10.1111/ele.70355 |
| Necrophagous beetles | https://doi.org/10.5281/zenodo.17632795 |  |
| Hoverflies | https://doi.org/10.5281/zenodo.16309483 | Massó Estaje et al., 2026, https://doi.org/10.1111/1365-2664.70252 |
| Flies and hymenopterans | https://doi.org/10.5281/zenodo.15371331 | Rothacher et al., 2025, https://doi.org/10.1111/1365-2664.70086 |
| Bees | https://doi.org/10.5281/zenodo.16837614 |  |
| Ants | https://doi.org/10.5281/zenodo.14989429 |  |
| Lepidopterans | https://doi.org/10.5281/zenodo.14282139 |  |
| Bats | https://doi.org/10.5281/zenodo.14766334 | Wild et al., 2026, https://doi.org/10.1016/j.cub.2026.04.058 |
| Birds | https://doi.org/10.5281/zenodo.15835876 | Wild et al., 2026, https://doi.org/10.1016/j.cub.2026.04.058 |

**Table S12:** Details on calculation of the Hill numbers for the different taxonomic groups. Coverage_stan_ refers to the coverage to which biodiversity in all plots was standardized by inter-or extrapolation.

| Taxonomic group | Observational unit | Data type | Coverage<sub>stan</sub> |
| --- | --- | --- | --- |
| Vascular understory plants | Species | Abundance (% cover) | No standardization |
| Terricolous bryophytes | Species | Abundance (% cover) | No standardization |
| Deadwood bacteria | OTU | Abundance (number of reads during metabarcoding) | 0.97 |
| Soil bacteria | OTU | Abundance (number of reads during metabarcoding) | 0.97 |
| Deadwood fungi | OTU | Abundance (number of reads during metabarcoding) | 0.99 |
| Soil fungi | OTU | Abundance (number of reads during metabarcoding) | 0.98 |
| Nematodes | Genera/families | Abundance (number of individuals) | 0.60 |
| Gastropods | Species | Abundance (number of individuals) | 0.63 |
| Spiders | Species | Abundance (number of individuals) | 0.35 |
| True bugs | Species | Abundance (number of individuals) | 0.35 |
| Saproxyllic beetles | Species | Abundance (number of individuals) | 0.29 |
| Non-saproxyllic beetles | Species | Abundance (number of individuals) | 0.83 |
| Necrophagous beetles | Species | Abundance (number of individuals) | 0.80 |
| Hoverflies | Species | Abundance (number of individuals) | 0.41 |
| Flies | OTU | Abundance (number of reads during metabarcoding) | 1.00 |
| Bees | Species | Abundance (number of individuals) | 0.45 |
| Ants | Species | Incidence-frequency | 0.72 |
| Hymenopterans | OTU | Abundance (number of reads during metabarcoding) | 1.00 |
| Lepidopterans | Species | Abundance (number of individuals) | 0.62 |
| Bats | Species | Incidence-frequency | 0.41 |
| Birds | Species | Incidence-frequency | 0.97 |

## Notes

### Note S1: Microclimatic data processing

Air temperatures at 2 m height were measured in half-hourly intervals in each plot from June 1^st^ to August 31 in 2023. The raw data were processed using the R packages `myClim` v. 1.5.0 (Man et al., 2023) and `myClimGui` v. 1.1.5 (Man et al., 2025). Processing steps included joining time series from different files, automatic cleaning, visual inspection and manual cleaning, and linear interpolation for gaps of five or less missing observations. Larger data gaps were imputed using the R package `mtsdi` v. 0.3.7 (Junger 2025). Next, the time series were aggregated to a daily resolution by calculating the maximum temperature of each day. Finally, these values were again aggregated to one value per plot by calculating the average daily maximum temperature over the whole study period.

### Note S2: Biodiversity sampling protocols

#### Understory vascular plants and terricolous bryophytes

Occurrence and cover of understory vascular plants and terricolous bryophytes were recorded in five circular subplots per plot. Each subplot had a radius of 4 m (50.27 m²). One subplot was located at the center of the plot, while the remaining four were positioned at the corners. The corner subplots were located either 12 m from the patch center (six sites) or 21 m from the center (five sites), owing to the design of a previous study. All understory vascular plant species were recorded, including seedlings and juvenile woody individuals, taking into account all individuals up to 1 m in height.

#### True bugs, beetles, gastropods, spiders, and ants

Each plot was equipped with two pitfall traps (capturing beetles, gastropods, spiders, and ants) and two flight interception traps with yellow bait strips (capturing true bugs and beetles). Traps were positioned in the northwest and southeast, at 7 m from the patch center. They were active continuously from May to July and emptied monthly at the end of May, June, and July. A 15% sodium chloride (NaCl) solution with a drop of odourless detergent was used as trapping fluid, and collected specimens were preserved in 70 % ethanol for preserving the samples after the collections. Samples were sorted into taxonomic groups and identified to species level.

#### Necrophagous beetles

Necrophagous beetles were sampled using pitfall traps baited with rat carrion (*Rattus norvegicus)*. Traps were installed for 48h in spring (May) and in summer (August/September), respectively. The contents were stored in 70% ethanol. Necrophagous beetles of the families, Silphidae, Leiodidae, Dermestidae, Geotrupidae, Scarabaeidae, and Nitidulidae were identified to species level morphologically.

#### Bees and hoverflies

Bees (Apidae) and hoverflies (Syrphidae) were sampled using standardized pan traps. In each patch, three clusters consisting of one yellow, one white, and one blue pan trap were placed along a transect crossing the plot centre, with a minimum distance of 15 m between clusters. Traps were filled with water and a drop of detergent and exposed for 48 h per sampling round. The sampling campaigns took place during May, June and July of 2022 and 2023. Collected specimens were sorted by taxonomic group and identified to species level.

#### Lepidopterans

Lepidopterans were sampled using one light trap per patch within a maximum radius of 15 m from the patch center and suspended from a tree at a height of 1.50–1.70 m. Traps were active for one night in June and one night in July, with additional sampling nights in May and August to compensate for unsuitable sampling conditions. All collected specimens were identified to species level and counted.

#### Nematodes

Soil sampling for nematode extraction took place in October 2023. At each patch, we collected four soil cores (5 cm diameter, 10 cm depth) after roughly removing the litter layer. Sampling locations within a plot were selected to capture variation in deadwood presence and light conditions (Schwarz et al., 2026). Cores from each plot were pooled, sieved (2 mm), and kept at 4 °C prior to further processing. Nematodes were extracted from approximately 25 g of fresh soil per patch using the Baermann funnel technique (Cesarz et al., 2019). After 72 hours, the nematode suspension was collected by passing the tube contents through a 15 μm sieve, and specimens were preserved directly in 4 % formalin. Soil dry weight was determined for each sample. Nematodes were counted under 50× magnification, and a subset of individuals (target: 100 per sample) was identified to genus level (adults and most juveniles) or family level (remaining juveniles) at 400× magnification on a Leica DMI 4000B microscope, following the identification keys of Bongers (1994). Diversity analyses were performed using raw abundance data.

### Fungi and bacteria in soil and deadwood

#### Soil and deadwood sampling

Soil was sampled once at each site during the autumn period (September–October 2022 in Bavarian and University forests and 2023 at all other sites). A consistent sampling design was applied across all plots. At each plot, the approximate geometric center was used as the reference point; in the University Forest, this position was marked permanently with a plastic marker bearing the site number. One soil core was taken 1 m north of the reference point, except in the University Forest, where the core was collected 1 m to the south. Four further cores were collected 2 m from the reference point in the four cardinal directions (N, E, S, and W), resulting in a cross-shaped sampling arrangement designed to capture small-scale spatial variability within plots. Surface litter, including undecomposed leaves and needles, was removed before sampling. Soil cores were collected to approximately 15 cm depth using a plastic corer with an internal diameter of 4.5 cm.

The five cores from each plot were kept at 4 °C and processed within 48 h. Roots and remaining litter were removed, after which the samples were passed through a sterile 5-mm sieve and thoroughly homogenized. Only the upper 10 cm of the soil was retained, and the material from all five cores was combined to obtain a single composite sample per plot. Samples were frozen at −40 °C, freeze-dried, and subsequently stored at −20 °C until further processing.

Deadwood sampling comprised ten objects per plot, selected by pre-stratified random sampling to represent the respective treatment in terms of deadwood type and position. Sampling was conducted during September–October according to a standardized procedure. At each sampling location, the outermost wood layer was removed with a knife from both sides of the object, and the exposed surface was disinfected with ethanol before drilling. Wood material was then collected using an 8-mm-diameter drill to a depth corresponding to the available wood width, with a maximum drilling depth of 20 cm. At least two drillings were made from each deadwood object. Four drillings were taken from logs longer than 2 m and from standing deadwood or stumps exceeding 1 m in height. Drill shavings from all sampling points were pooled to produce one composite deadwood sample representing the ten objects within each plot. Specifically, 2 g of material per sample was taken, resulting in 10 objects x 2 g = 20 g pooled plot material. Samples were frozen at −40 °C. The pooled samples were freeze-dried, finely milled, and stored at −20 °C until DNA extraction.

#### DNA extraction and PCR amplification

DNA was extracted from 150 mg of freeze-dried soil and 100 mg of the finely milled, freeze-dried, pooled deadwood samples. Extraction followed a modified Miller protocol (Sagova-Mareckova et al., 2008), including bead beating and phenol–chloroform extraction performed using a FastPrep system (MP Biomedicals). Three parallel DNA extractions were carried out for each sample and subsequently purified with the GeneClean Turbo Kit (MP Biomedicals, Solon, OH, USA). DNA concentration and purity were evaluated using a NanoDrop ND-1000 spectrophotometer (Thermo Fisher Scientific, Waltham, USA), and the three extracts were then combined.

For each sample and primer pair, three independent PCR reactions were performed to minimize potential amplification biases. Fungal communities were assessed by targeting the ITS2 region of the fungal rRNA gene with the barcoded gITS7 and ITS4 primers (Ihrmark et al., 2012), using the protocol described by Žifčáková et al. (2016). Bacterial communities were characterized by amplification of the V4 region of the bacterial 16S rRNA gene using the barcoded 515F and 806R primers (Caporaso et al., 2011).

PCRs were conducted in a final volume of 25 µl. For the soil communities, each reaction contained 1 µl of template DNA (5 ng µl⁻¹), 5 µl of Q5 Reaction Buffer (New England Biolabs, Inc.), 1.5 µl of bovine serum albumin (10 mg ml⁻¹; GeneON), 1 µl of each primer (10 µM; Sigma-Aldrich), 0.25 µl of Q5 High-Fidelity DNA polymerase (New England Biolabs, Inc.), 5 µl of Q5 High GC Enhancer (New England Biolabs, Inc.), 0.5 µl of PCR Nucleotide Mix (10 mM; Bioline), and 9.75 µl of molecular-grade water. For the deadwood community, for fungal ITS amplification, each reaction contained 12.5 µl of KAPA HiFi HotStart ReadyMix (Sigma-Aldrich), 9.5 µl of molecular-grade water, 2 µl of a primer mixture (Sigma-Aldrich) containing 10 µM of each forward and reverse tagged primer, and 1 µl of template DNA. For bacterial 16S amplification, reactions were prepared using GoTaq® G2 Hot Start Master Mix (Promega) and contained 12.5 µl of master mix, 9.5 µl of molecular-grade water, 2 µl of a primer mixture (Sigma-Aldrich) containing 10 µM of each forward and reverse tagged primer, and 1 µl of template DNA. Template DNA concentrations were approximately 5–50 ng µl⁻¹.

For soil fungal ITS2 amplification, PCR cycling comprised an initial denaturation at 94 °C for 5 min, followed by 35 cycles of denaturation at 94 °C for 30 s, annealing at 56 °C for 30 s, and extension at 72 °C for 30 s. A final extension step was performed at 72 °C for 7 min. Amplification of the soil bacterial 16S rRNA gene consisted of an initial denaturation at 94 °C for 4 min, followed by 25 cycles of denaturation at 94 °C for 45 s, annealing at 50 °C for 1 min, and extension at 72 °C for 75 s, followed by a final extension at 72 °C for 10 min. For the deadwood fungal ITS amplification, PCR cycling comprised an initial denaturation at 98 °C for 30 s, followed by 30 cycles of denaturation at 98 °C for 10 s, annealing at 56 °C for 30 s, and extension at 72 °C for 30 s. A final extension step was performed at 72 °C for 2 min. Amplification of the bacterial 16S rRNA gene consisted of an initial denaturation at 98 °C for 30 s, followed by 25 cycles of denaturation at 98 °C for 10 s, annealing at 50 °C for 1 min, and extension at 72 °C for 1 min, followed by a final extension at 72 °C for 2 min.

The three PCR replicates were pooled for each sample and purified using the MiniElute PCR Purification Kit (Qiagen, Hilden, Germany) in the case of the soil community, and with AMPure XP beads (Beckman Coulter Life Sciences), in the case of the deadwood community. Purified amplicons were quantified using a Qubit 2.0 fluorometer with the Qubit dsDNA BR Assay Kit (Life Technologies/Thermo Fisher Scientific). Amplicon libraries were subsequently prepared using the TruSeq DNA PCR-Free Kit (Illumina) and sequenced on an Illumina MiSeq platform using MiSeq Reagent Kit v2 chemistry with 2 × 250-bp paired-end sequencing.

#### Amplicon sequence processing, taxonomic assignment, data availability

Fungal ITS amplicon sequences were processed using SEED v2.1.3 (Větrovský et al., 2018). Paired-end reads were merged with fastq-join (Aronesty, 2013), followed by extraction of the ITS2 region using ITSx v1.0.8 (Nilsson et al., 2010) and removal of chimeric sequences. The resulting sequences were clustered into operational taxonomic units (OTUs) at a 97% sequence-similarity threshold using UPARSE (Edgar, 2013). Representative sequences were selected according to their abundance and taxonomically classified by BLASTn searches against the UNITE v10 database (Abarenkov et al., 2023) and GenBank. Non-fungal sequences were excluded from the final dataset. Bacterial sequences were processed using an edited version of the SILVA v138.1 database, in which mitochondrial and chloroplast sequences were retained to facilitate their identification during taxonomic assignment.

The raw sequencing data are available in SRA under the Bioproject Numbers: PRJNA1232009 (soil bacteria, 16S), PRJNA1232019 (soil fungi, ITS2), PRJNA1435076 (deadwood bacteria 16S) and PRJNA1435086 (deadwood fungi, ITS2).

#### Flies and hymenopterans

We deployed one Malaise trap (height front: 0.90 m; height rear: 0.60 m; length: 1.60 m) in the central area of each experimental patch. Traps were operated continuously from May to July. Collection bottles of the malaise traps were equipped with 70% ethanol as a preservative solution. Samples were collected fortnightly, resulting in six sampling periods per patch (n = 1403 samples; 234 patches across all sites × 6 sampling events, one sample lost due to a storm). Samples were sieved (8 mm mesh) and species were identified using COI-5P (mitochondrial cytochrome c oxidase subunit I) metabarcoding (for details see Rothacher et al. 2025).

#### Bats

Bat activity was recorded using one batcorder per patch, installed within 15 m of the patch center and suspended from a tree at 1.50–1.70 m height. Recorders were active for one night per sampling session, from one hour before sunset to one hour after sunrise, primarily in June and July and additional sampling in May and August. Sites in the University Forest and Bavarian Forest were sampled in 2022, and those in Saarland, Hunsrück, and Lübeck in 2023. Recordings were processed using bcAdmin4 (v1.2.6) and identified with batIdent (v1.5; ecoObs GmbH), using a minimum detection probability of 0.6. Only species-level identifications were retained, excluding species not known from the respective regions. Bat activity was quantified as minute counts and converted to species presence–absence per sampling night for subsequent analyses (Wild et al. 2026).

### Birds

Birds were sampled using one autonomous sound recorder (BAR or BAR-LT; Frontier Labs, Australia) per patch, installed within 15 m of the patch center at 1.50–1.70 m height. Recorders operated daily from March to June, recording for 2 min every 12 min around sunrise and sunset. Recordings from March 24 to June 15 were analyzed using BirdNET v2.4 with species-specific confidence thresholds based on expert-validated identifications. Implausible species records were manually verified and excluded when erroneous. Detections were aggregated to daily species presence–absence (Wild et al. 2026).

## Notes S3: Supplementary methods

### Contents of this document

This document describes the structure of the Bayesian hierarchical models used to model biodiversity data. Section **S3.1** describes the models for the multi-taxon diversity data, while Section **S3.2** focuses on the models with taxon-specific data. All models were fitted in R v.4.5.2 (R Core Team 2025) in a Bayesian framework based on R package brms v.2.23.0 (Bürkner 2018). Modelling was done in the framework of the tidyverse v.2.0.0 (Wickham et al. 2019).

### S3.1 Model for multi-taxon diversity

#### S3.1.1 Model structure

As the multi-taxon diversity index according to Allan et al. (2014) is based on averaging the values of the diversity indices of all taxa relative to their maximum value, it has a strict lower bound at 0 and a strict upper bound at 1. We therefore assumed the observed multi-taxon diversity to follow a beta distribution. We fitted three separate models for multi-taxon diversity indices calculated with Hill numbers of order q = 0, 1 and 2.

For the regression models, we used a reparameterization of the beta distribution with mean *µ* and precision *ϕ*, which can be derived from the shape parameters *α >* 0 and *β >* 0 as *µ* = *α/*(*α* + *β*) and *ϕ* = *α* + *β* (0 *< µ <* 1, *ϕ >* 0), resulting in the following density function:

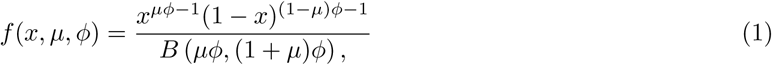

where *B*(*·*) is the beta function. The calculated multi-taxon diversity *M_i_* for plot *i* in (1*…I*) from district *j*in (1*…J*) in site *k* in (1*…K*) was modelled according to such a Beta distribution:

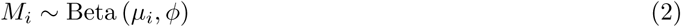

The expected value *µ_i_* of this Beta distribution was expressed with a linear predictor term using a logit link:

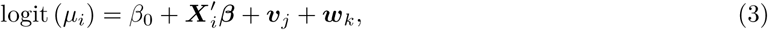

where *β*_0_ is an overall intercept, *X_i_’β* is a linear predictor term for the effects of the predictors in the model, and *v_j_* and *w_k_* are random intercepts on district-and site-level, respectively. Here, ***β*** is a vector of length *n* containing the main effects for the *N* predictor variables, and ***v*** and ***w*** are vectors of length *J* and *K*, respectively with the random district and site effects for each taxonomic group. ***X*** is a *I × N* model matrix without intercept containing columns for *N* predictor variables, namely canopy cover, deadwood volume, and macro-and microclimate as well as their interaction. All predictor variables (including the interaction term) were scaled and centred before inclusion into the model to permit comparing effect sizes across variables and simplify prior choices. Before scaling and centring, canopy closure was further transformed with a scaled logit transformation in the form *x_trans_* = logit(0.99*x/*100 + 0.005) to permit a more even spread of the canopy values and reduce the high leverage of a small number of plots with low canopy cover.

The district-and site-specific random intercepts ***v*** and ***w*** were assumed to be normally distributed with mean 0 and standard deviations *τ_v_* and *τ_w_*, respectively.

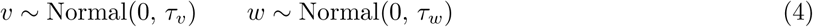

#### S3.1.2 Prior specifications

We used moderately informative priors for all model parameters. To simplify prior specification for the regression parameters, all predictor variables were scaled and centred before the analysis.

For the intercept *β*_0_, we used a normal prior centred around the mean *µ_logitM_* of the logit-transformed multi-taxon diversity numbers on the corresponding level of *q* with a standard deviation equal to the standard deviation *σ_logitM_* of the same data:

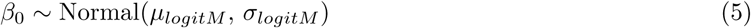

Due to the centred predictor variables, *β*_0_ can be interpreted as the grand mean of the data on the scale of the linear predictor and should thus be situated within the range specified by this prior.

For the remaining regression parameters *β*, we used a normal prior with a mean of 1.5:

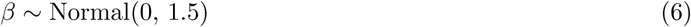

On the scale of the data, this prior places 95 % of probability mass on slopes that constrain the response to a range of approximately 0.05–0.95 when changing a predictor variable from *−*1 to +1 standard deviation, i.e., it permits a change of nearly the entire possible range of the response over the covered range of the predictor variable.

For the standard deviations of the random intercepts on district and site level (*τ_v_* and *τ_w_*, respectively), we used half-normal prior with a mean of zero and a standard deviation of *σ_logitM_*.

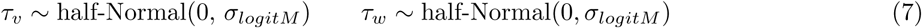

This prior constrains the standard deviations of the random intercepts to the range of the data on the scale of the linear predictor.

The precision parameter *ϕ* was assigned the brms default Gamma prior:

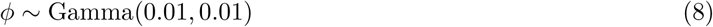

This prior has an expected value of 1, but a high variance of 100 that makes it weakly informative over a wide range of potential values.

#### S3.1.3 Model implementation and checking

Stan performs Markov Chain Monte Carlo (MCMC) sampling using Hamiltonian Monte Carlo (HMC) based on the no-U-Turn sampler (Hoffman and Gelman 2014). The sampling was run for 10000 iterations (5000 of which were discarded as a warmup interval) on 4 chains using a maximum tree depth of 12 and a target acceptance rate of adapt_delta = 0.99. None of the models showed any divergent transitions. We assessed model convergence based on the rank-based improved *R*^^^ convergence statistic (Vehtari et al. 2020), as well as visual inspection of trace plots of the most important model parameters.

The models were validated based on prior and posterior predictive checks (cf. Gabry et al. 2019), formal model validation based on R package DHARMa v.0.4.7 (Hartig 2024) via DHARMA.helpers v.0.0.2 (Rodríguez-Sánchez 2026). In addition, we performed approximate leave-one-out cross-validation based on Pareto-smoothed importance sampling (Vehtari et al. 2017) based on R package loo v.2.9.0 (Vehtari et al. 2025) using moment matching to identify influential observations and to identify potential model misspecification. For all values of q, none of the observations had pareto k values above 0.7 that are indicative of influential observations.

### S3.2 Structural model of multi-taxon diversity

To test whether the directed acyclic graph in Fig. 1 in the main text is consistent with the data, we used a path modelling approach using piecewise models building upon Shipley (2000). In this framework, we further tested whether there is an indirect effect of forest structure on biodiversity that is mediated by microclimate.

#### S3.2.1 Model structure

Besides the obvious effects of canopy cover, microclimate, macroclimate and deadwood on biodiversity, which are covered by the model in Eqn. 4, the DAG in Fig. 1 implies an effect of canopy cover on microclimatic temperature. We modelled this as a linear model with a variance function for canopy cover, as the variance in microclimate was higher in more open plots:

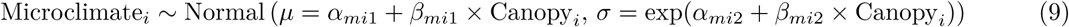

As we assumed this effect to purely result from the physical effect of forest structure on climate, we did not add site-specific random effects. A model with such random effects was tested, but had a ΔELPD of-1.7 (SE: 3.0) in a leave-one-out cross-validation based comparison with the model in Eqn. 9, indicating an equivalent fit.

Besides the relationships described by Eqn. 4 and Eqn. 9, the DAG in Fig. 1 implies the following conditional independence claims that constitute the basis set of that DAG: deadwood *⊥* macroclimate, deadwood *⊥* canopy cover, microclimate *⊥* deadwood | canopy cover, macroclimate *⊥* canopy cover, microclimate *⊥* macroclimate | canopy cover.

We tested these conditional independence claims via residual correlations in multivariate regressions.

As brms does not permit to specify pairs of variables *without* residual correlations when setting rescor(TRUE), we had to split the computation into separate models, as the alternative would have been to rewrite the model code manually in Stan. We fitted the following set of models:

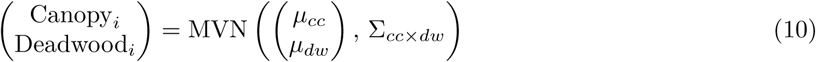

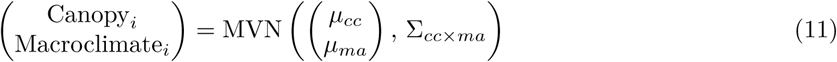

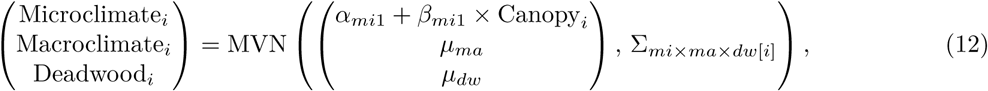

with MVN(*·*) denoting the multivariate normal distribution and Σ*_cc×dw_* and Σ*_cc×ma_* being 2 *×* 2 covariance matrices containing each two standard deviations *σ* and a residual correlation *ρ*:

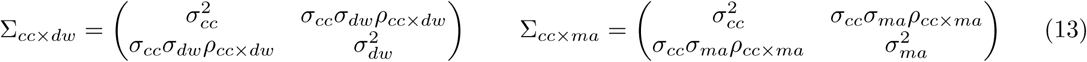

Due to the variance model in Eqn. 9, the expression for Σ*_mi×ma×dw_* has to be evaluated separately for every observation *i*, with *σ_mi_*_[*i*]_ = exp(*α_mi_*_2_ + *β_mi_*_2_ *×* Canopy*_i_*):

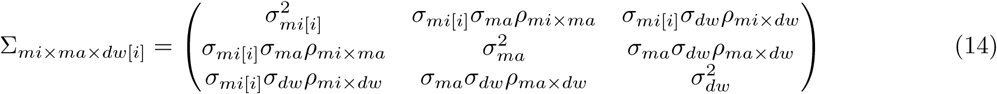

The models in Eqn.10, 11 and 12 permit inference on the microclimate-canopy cover relationship implied in Eqn. 9 while estimating the full set of residual correlations needed to test the conditional independence claims mentioned above. An inclusion of the model for multi-taxon diversity is not required in this context, as all of the other variables are already accounted for as predictor variables in this model.

#### S3.2.2 Prior specifications

For the models in Eqn.10, 11 and 12, we used the *brms* default priors, i.e., a Student-t(3, *x̄*, 2.5) prior on the intercept terms centred around the median *x̄* of the data, an improper flat prior onthe parameter vectors, a Student-t(3, 0, 2.5) prior on the intercept in the parameter regression for *σ_mi_*, and and LKJ(1) prior on all correlation matrices.

#### S3.2.3 Model implementation and checking

Models were fitted for 10000 iterations with a warmup interval of 5000 iterations on 4 chains using a target acceptance rate of adapt_delta = 0.9. None of the models showed any divergent transitions. Model validation was performed as described in Section S3.1.3.

#### S3.2.4 Tests of directional separation

Based on the residual correlations estimated in Eqn.10, 11 and 12, we tested whether the causal mode in Fig. 1 is an adequate representation of the data. As can be seen in the following table (where CC = canopy cover, DW = deadwood, MI = microclimate, MA = macroclimate), there was credible evidence for residual correlations for the deadwood-canopy cover relationship and the deadwood-macroclimate relationship:

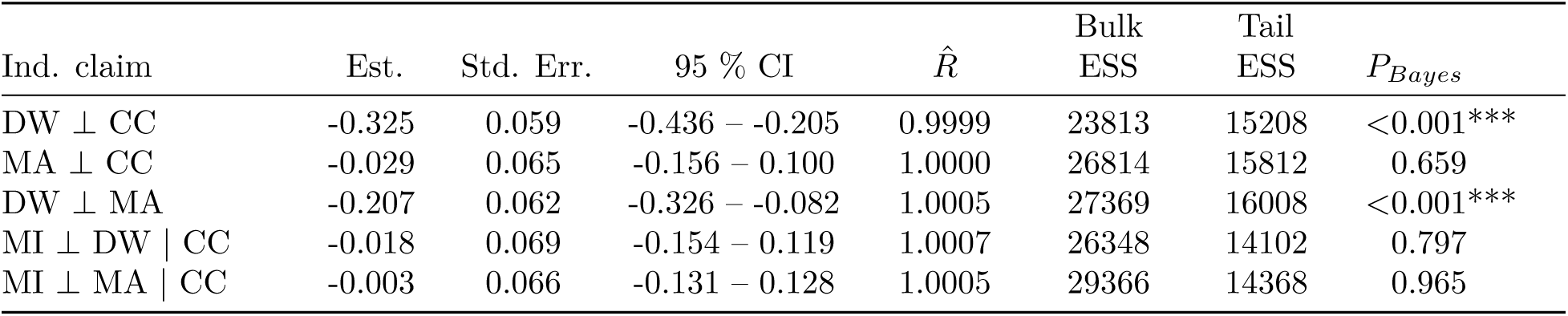

Here, *P_Bayes_* is a Bayesian pseudo-P-value based on the probability of direction. The probability of direction *PD* is the proportion of a posterior of a sampled parameter has the same sign as its median (Makowski et al. 2019), and *P_Bayes_*is defined as follows:

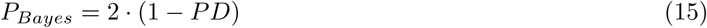

In a model with flat priors, *P_Bayes_* is equal to the two-sided frequentist p value.

As to our knowledge there is no Bayesian test for the consistency of a full structural model with a dataset, we then performed Fisher’s C test based on the *k* estimated values of *P_Bayes_* by computing the following test statistic:

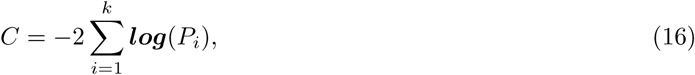

which according follows a chi-squared distribution with 2*k* degrees of freedom if the null hypothesis is correct (Shipley 2000).

For the full set of independence claims, this test was highly significant (P < 0.001) due to the strong correlations between deadwood and both macroclimate and canopy cover. After adjusting the original structural model and accounting for these residual correlations, the resulting model (compare Fig. 4 in the main text) was consistent with the data (*χ*^2^ = 1.362, df = 6, P = 0.968).

#### S3.2.5 Test of mediation

To conclude on the existence of an indirect effect of forest structure on biodiversity that is mediated by its effect on microclimate, it is necessary to demonstrate that a) there is an effect of canopy closure on microclimatic temperatures, and b) that there is an effect of microclimatic temperatures on biodiversity. To test a), it is sufficient to test whether the credible intervals of the parameter *β_mi_*_1_ Eqn. 9 exclude zero. To test b), it is necessary to compute the complete effect of microclimate on biodiversity, as it differs between sites due to the interaction with macroclimate. We therefore computed the total effect of microclimate for each site as the sum of the regression coefficient for microclimate and the product of the interaction coefficient and the macroclimatic temperature at that site. We then tested whether the credible intervals of this total effect exclude zero separately for each of the sites.

### S3.3 Model for taxonomic groups

For the biodiversity data of individual taxonomic groups, we fitted joint models for all taxonomic groups with a zero-altered gamma distribution with a flexible mean-variance relationship. Three separate models were fitted for Hill numbers of the order q = 1, 2 and 3.

#### S3.2.1 Modified gamma distribution

##### S3.2.1.1 Gamma distribution with custom dispersion model

The probability density function of the gamma distribution parameterized with shape *α >* 0 and rate *λ >* 0 is as follows:

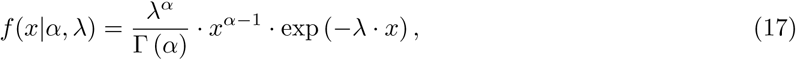

where Γ(*·*) denotes the gamma function.

For regression, Eqn. 17 is usually reparameterized with the expected value *µ* = *α/λ*:

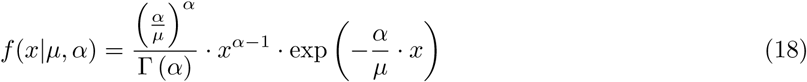

In gamma regression models, ***µ*** is then linked to a linear predictor ***Xβ*** using a logarithmic link function:

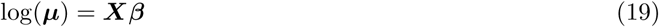

The variance *σ*^2^ of the gamma distribution is *α/λ*^2^, which implies a quadratic variance-mean relationship:

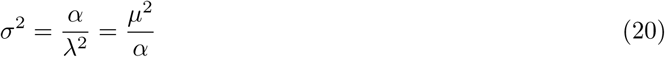

This increase in variance with the mean is adequate for a large number of strictly positive variables occurring in nature, but turned out problematic for our model based on coverage-standardized Hill numbers, which often showed a less pronounced (and, in some cases, even a stronger) increase in variance with the mean than expected by the gamma distribution. To account for these patterns, we added an additional parameter *b >* 0 that flexibilizes the variance-mean relationship in the following form:

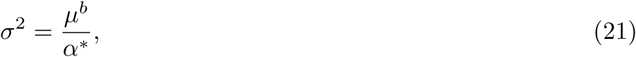

where *α^∗^* denotes the modified shape parameter, which is equal to the shape parameter *α* of a regular gamma distribution only for *b* = 2, under which conditions Eqn. 21 reduces to Eqn. 20.

Reparameterizing the probability density function of the gamma distribution with mean *µ* and variance *σ*^2^ by rearranging Eqn. 20 and substituting into Eqn. 18 results in the following expression:

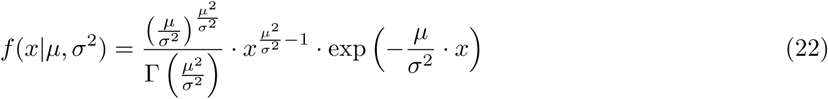

By substituting *σ*^2^ = *µ^b^/α^∗^* into Eqn. 22, the modified gamma density with additional parameter *b* can be derived as follows:

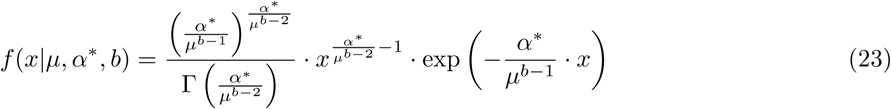

It can be seen that if *b* = 2, Eqn. 23 reduces to Eqn. 18. For values of *b >* 2, the variance increases with the mean more strongly and for values of *b <* 2 less strongly than expected by a regular gamma distribution. In the following, the density function of this modified gamma distribution with a custom dispersion model will be denoted as Gamma_disp_ (*x|µ, α^∗^, b*).

##### S3.2.1.2 Zero alteration model

Our biodiversity data contained a number of true zeros in cases where not a single organism of the corresponding taxonomic group was present in a plot. The probability density of the gamma distribution at zero is 0 for values of *α >* 1, 1 for *α* = 1 and infinite for *α <* 1. For that reason, data including zeros cannot be easily modelled with the gamma distribution.

One common way to model positive data including zeros is the hurdle gamma or zero-adjusted gamma model, which assigns a probability of *ν* to observing a value of zero. Using the gamma distribution with dispersion model as defined in Eqn. 23, this results in the following model:

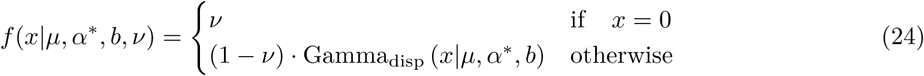

The parameters *µ* and *ν* are then assigned separate linear predictor terms using a log and logit link, respectively:

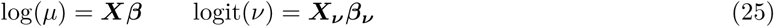

This is especially useful if the zeros in *x* are expected to come from a different data generating process than the rest of the data, for instance a sensor that measures concentrations of volatile compounds that gets shut off for safety reasons and thus returns values of zero when wind speed is too high.

In our case, since detectability issues would affect all observations equally and there are no obvious processes that could explain the observed zeros, it is more appropriate to assume them to come from the same process as the rest of the data. Using separate linear predictors for *µ* and *ν* with the same predictor terms in Eqn. 25 to account for this would rapidly lead to very complex models. In the other extreme, leaving *ν* as a constant without a predictor term is equivalent to estimating the proportion of zeros as *ν* and then fitting a model with the non-zero observations only.

To be able to use the zero observations to inform our inferences on the linear predictor for *µ*, we set up a model using the same linear predictor with an additional parameter *−∞ < c < ∞* analogous to a cutpoint value in ordered logistic regression:

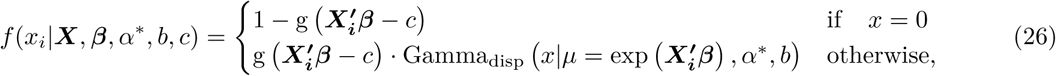

where g(*·*) denotes the inverse logit function. Expressed more generally in terms of the expected value *µ* of the Gamma_disp_ component, this is equivalent to:

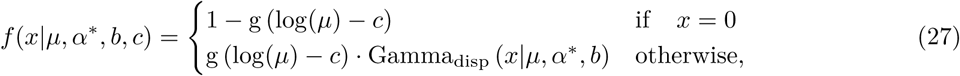

Due to the analogy to ordered logistic regression and in particular to the ordered beta model for zero/one-inflated beta regression (Kubinec 2022), in the following, this model will be referred to as the ordered gamma model or orderedGamma_disp_ (*x|µ, α^∗^, b, c*).

In this model, the probability of observing non-zero values is a function of the same linear predictor as expected value of the response, and is lower for lower expected values, such that for very low expected values, observations of zero become very likely. If the linear predictor of the model is equal to *c*, the probability of observing a 0 is 0.5. Higher values of *c* lead to a higher baseline probability of observing zeros and vice versa.

For the ordered gamma model, the expected value including zeros is simply the expectation of the gamma component *µ* times the probability of observing non-zero values:

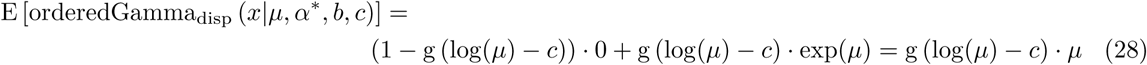

##### S3.2.1.3 Implementation of the modified gamma model

To fit the model in Eqn. 27 with brms, we implemented it as a custom response distribution according to vignette(“brms_customfamilies”) (cf. Bürkner 2018).

The Stan code for the log probability density function of the ordered gamma model with dispersion parameter *b* (here named bg – short for ‘b gamma’ – to avoid confusion with the regression coefficients) is as follows:

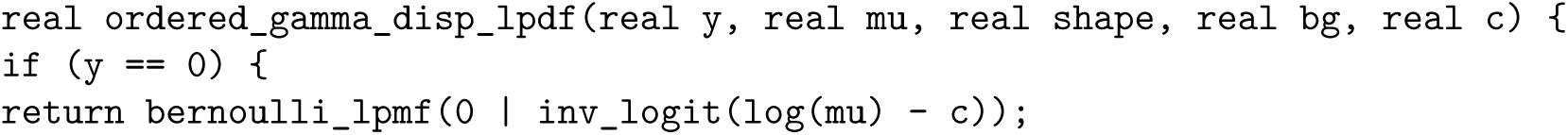
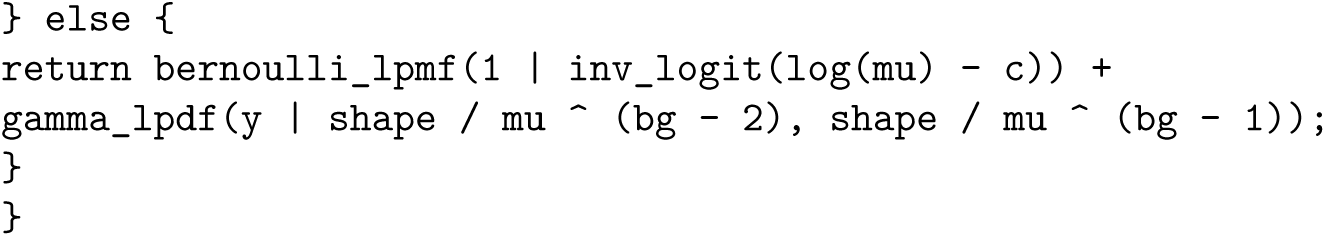

This code can be implemented as a custom family following the instructions in vignette(“brms_customfamilies”), and included in a brm call via the stanvars argument.

To be able to quantify whether the custom model structure actual is an improvement over standard model formulations, we further also fitted a model with the zero alteration structure, but without the dispersion model, as well as a model with the dispersion model, but a regular gamma hurdle formulation.

The zero-altered model without additional dispersion parameter can be implemented as follows:

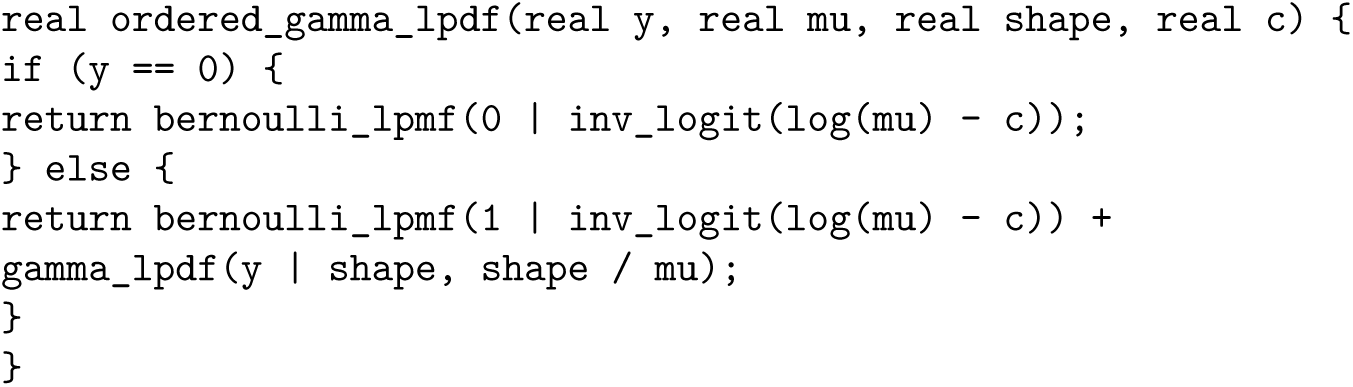

The hurdle model with modified dispersion structure can be implemented with the following Stan code:

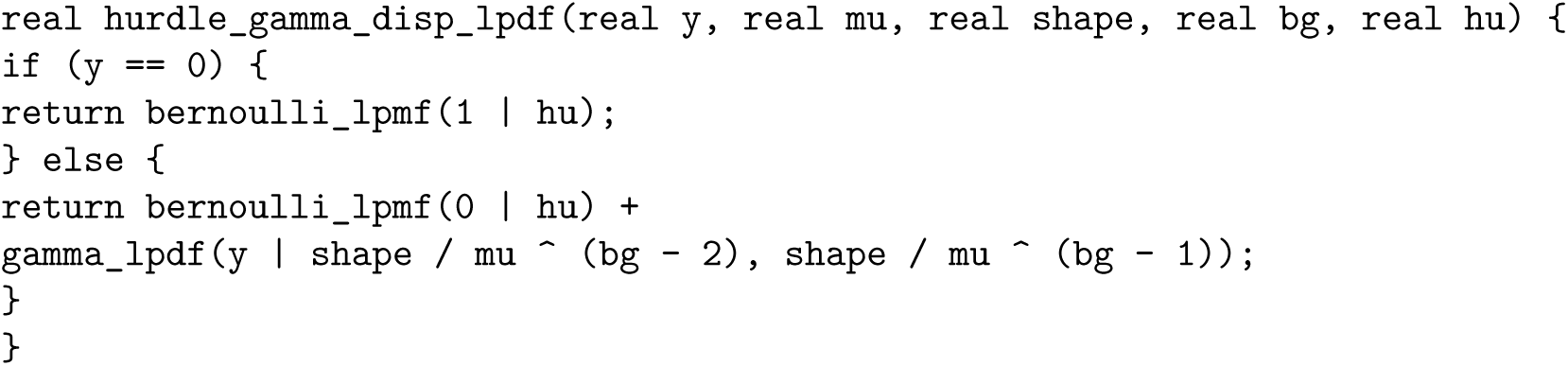

#### S3.2.2 Model fitting

##### S3.2.2.1 Model structure

We modelled the coverage-standardized Hill numbers for *q* = 0, 1 and 2 in three separate models. In our model, the Hill number H for the observation of plot *i* in (1*…I*) from district *j* in (1*…J*) in site *k* in (1*…K*) for taxonomic group *l* in (1*…L*) was modelled based on the zero-altered gamma distribution with modified dispersion structure (orderedGamma_disp_(*·*)) defined in Eqn. 27 with taxonomic-group-specific distributional parameters α_i_*, *b_l_* and *c_l_*:

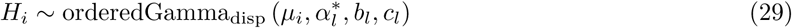

The logarithm of the expected value *µ_i_* of the Gamma component of the model was expressed as a linear function of an overall intercept *β*_0_, a linear predictor term ***X_i_*^’^***β* for the overall effects, a second linear predictor term *Z_i_***^’^***β* for taxonomic-group specific random intercepts and slopes, as well as random intercepts on district-and site-level (*v_jl_* and *w_kl_*, respectively).

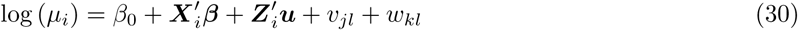

Here, ***β*** is a vector of length *n* containing the main effects for the *N* predictor variables, ***u*** is a *L ·* (*N* + 1) vector of taxonomic-group specific random intercepts and slopes, while ***v*** and ***w*** are vectors of length *J · L* and *K · L*, respectively with the random district and site effects for each taxonomic group. ***X*** is a *I × N* design matrix without intercept containing columns for *n* predictor variables, in our case canopy cover, deadwood volume, and macro-and microclimate as well as their interaction. Analogously, ***Z*** is a *I ×* ((*N* + 1) *· L*) design matrix with taxonomic-group specific terms for intercepts and all predictors in ***X***.

All predictor variables (including the interaction term) were scaled and centred before inclusion analogously to the model in **S3.1.1**.

The taxonomic-group-specific random effects ***u*** were assumed to be drawn from a multivariate normal distribution with mean 0 and covariance matrix **Σ***_u_*:

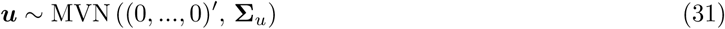

This covariance matrix can be decomposed into a correlation matrix **Ω*_u_*** and a vector of standard deviations

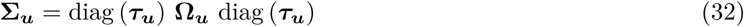

The district-and site-specific random intercepts *v* and *w* were assumed to be normally distributed with mean 0 and standard deviations *τ_v_* and *τ_w_*, respectively.

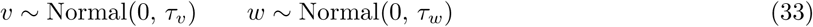

Finally, the taxonomic-group-specific distributional parameters α_i_*, *b_l_* and *c_l_* for taxonomic group *l* were described as follows:

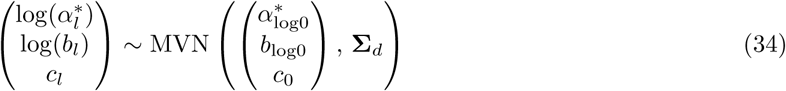

Accordingly, (α_i_*), log(*b_l_*) and *c_l_* were assumed to follow a multivariate normal distribution around their average values α_log0_*, *b*_log0_ and *c*_0_ with covariance matrix **Σ***_d_*. The shape α_i_*, and dispersion parameter *b_l_* were modelled with a log-link as they are constrained to be strictly positive.

Again, **Σ***_d_* can be decomposed into a correlation matrix **Ω*_d_*** and a vector of standard deviations ***τ_d_***:

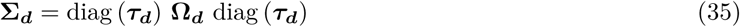

##### S3.2.2.2 Prior specifications

As in **S3.1.3**, we used moderately informative priors for all model parameters.

For the intercept *β*_0_, we used a relatively narrow normal prior centred around the mean *µ_logH_* of the log transformed non-zero Hill numbers on the corresponding level of *q* with a standard deviation of half of the standard deviation *σ_logH_* of these data:

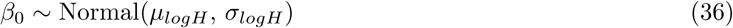

As the predictor variables are centred around their mean, *β*_0_ can be interpreted as the grand mean of the non-zero data on the scale of the linear predictor and should therefore squarely fall within the range of this prior.

For the remaining regression parameters *β*, we used a normal prior with a mean of *σ_logH_*:

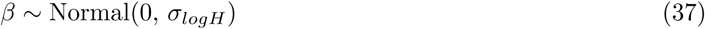

This prior places ca. 95 % of probability mass in the range of *±*2 standard deviations of the data on the scale of the linear predictor. Hence, this prior reflects the (realistic) a priori assumption that it is most likely that the effect of changing one of the (scaled) predictor variables by 1 standard deviation should correspond to a change in the transformed response of less than *±*2 standard deviation.

For the mean α*_log0_ of the shape parameter on the log scale, we used a normal prior centred around a value of log(4) with a standard deviation of 1:

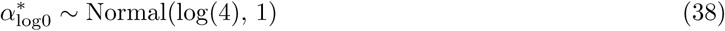

On the scale of *α^∗^*, this places 95 % of the prior probability mass on values between 0.6 and 28.4 and hence defines a broad range of potential average shape parameters.

For the mean *b*_log0_ of the additional dispersion parameter of our model on the log scale, we used a normal prior centred around a value of log(4) with a standard deviation of 1:

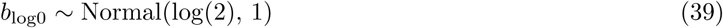

On the scale of *b*, this prior is centred around a value of 2, which, as mentioned above, corresponds to a regular gamma distribution. The prior places 95 % of the prior probability mass on values between 0.3 and 14.2 and hence allows a broad range of different average values for *b*.

The average cutpoint value *c*_0_ for zero alteration component of the model was assigned a normal prior centred around *µ_logH_ −* 4*σ_logH_* with with a standard deviation of 2*σ_logH_*:

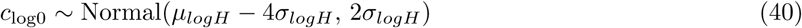

This places 95 % of the prior probability mass on values between the average of the data and *−*8*σ* below the average, and hence defines a broad range of values on the lower range of the data with a tendency towards lower values outside the range of the data. This is necessary, as for a majority of the taxonomic groups there are no zero observations, and the model relies on the prior containing *c* at low enough values to not result in any predicted values of zero for those groups.

For the standard deviations of the random intercepts on taxonomic group level (here denoted as *τ_u_*_[1]_), we used a half-normal prior with a mean of zero and a standard deviation of *σ_logH_*:

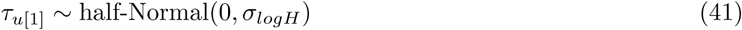

This prior contains the standard deviation of the random intercepts in the range of the standard deviation of the data on the scale of the linear predictor and hence constrains them to a realistic range without pushing them in any particular direction.

For the other taxonomic group level random effects *τ_u_*_[2*…*(*N*+1)]_, which denote the random slopes for the variables in *X*, we used a normal prior with mean 0 and a standard deviation of 0.5.

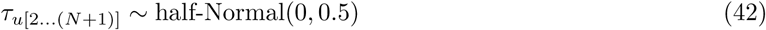

These relatively narrow priors on the random slopes constrain 95 % of the probability mass on the relative change in the linear predictor per standard deviation in the predictor variables within [exp(*−*1), exp(1)], i.e. between 0.4-fold and 2.7-fold changes compared to the main effect of the corresponding predictor.

The correlation matrix of the taxonomic-level random effects Ω*_u_*was assigned a LKJ(2) prior (Lewandowski et al. 2009).

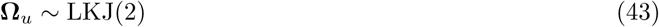

The LKJ(2) prior is a multivariate generalization of the boundary-avoiding Beta(2, 2) prior for correlation parameters (cf. Gelman 2014, p317) that rules out correlations of exactly *−*1 or 1 while not contradicting any other possible values.

For the standard deviations of the random intercepts on district and site level (*tau_v_* and *tau_w_*, respectively), we used half-normal prior with a mean of zero and a standard deviation of *σ_logH_* to constrain them to the range of the data (compare S3.1.3).

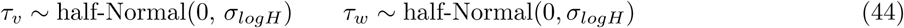

The random effects standard deviations for the distributional parameters *τ_d_* were assigned half-normal priors with a standard deviation of 1 (parameter *b*; *τ_d_*_[1]_) or 2 (parameters *α^∗^*, *c*; *τ_d_*_[2,3]_):

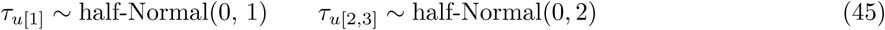

Again, the correlation matrix of the taxonomic-level random effects Ω*_d_* was assigned a LKJ(2) prior (Lewandowski et al. 2009).

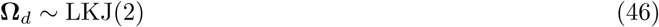

##### S3.2.2.3 Model implementation and checking

Sampling was run for 10000 iterations (5000 of which were discarded as a warmup interval) on 4 chains using a maximum tree depth of 12, a target acceptance rate of adapt_delta = 0.95 and an initial step size of 0.01. To avoid the sampler to move into regions of the posterior associated with very extreme values at the onset of sampling, we set init_r = 0.5 to initiate the chains with parameter values randomly sampled from a uniform distribution from [*−*0.5, 0.5] on the unconstrained scale (instead of the default value of init_r = 2, which initiates from [*−*2, 2]). None of the models showed any divergent transitions. We assessed model convergence based on the rank-based improved *R*^^^ convergence statistic (Vehtari et al. 2020), as well as visual inspection of trace plots of the most important model parameters.

Again, the models were validated based on prior and posterior predictive checks (cf. Gabry et al. 2019), residual diagnostics based on R package DHARMa v.0.4.7 (Hartig 2024) via DHARMA.helpers v.0.0.2 (Rodríguez-Sánchez 2026), and approximate leave-one-out cross-validation based on R package loo v.2.9.0 (Vehtari et al. 2025), computed with moment matching. For all models, only a very small percentage (*<* 0.4) of the observations had pareto k values above 0.7, indicative of influential observations. We checked with the data authors whether these observations are reliable or due to notation errors, but maintained in the dataset them unless there were clear reasons for removal and instead re-ran loo with the reloo = TRUE argument to refit the model (with a reduced number of 1000 iterations) for these observations to obtain reliable loo results for those observations.

We then performed a model comparison (Sivula et al. 2025) based on the theoretical expected log pointwise predictive density (ELPD) (Vehtari et al. 2017) obtained via leave-one-out cross-validation to test whether the modified model in Eqn. 27 performed better than a) an analogous model with a regular hurdle gamma formulation, but the same dispersion model (cf. Eqn. 24) and b) an analogous model with a zero alteration model as in Eqn. 27, but without the modified dispersion model. The regular hurdle model had a ΔELPD of-63.4 (SE: 7.6) compared to the full model, while the model without the dispersion model had a ΔELPD of-119.4 (SE: 19.9), which amounts to a difference of 8.3 and 6.0 standard errors, respectively. Accordingly, there is very strong evidence that both the zero alteration and the dispersion term considerably improved the fit of our model.

## References

1. Seibold, S. et al. Arthropod decline in grasslands and forests is associated with landscape-level drivers. Nature 574, 671–674 (2019).

2. Neff, F. et al. Different roles of concurring climate and regional land-use changes in past 40 years’ insect trends. Nat Commun 13, 7611 (2022).

3. Staab, M. et al. Insect decline in forests depends on species’ traits and may be mitigated by management. Commun Biol 6, 338 (2023).

4. Keck, F. et al. The global human impact on biodiversity. Nature 641, 395–400 (2025).

5. Aszalós, R. et al. Natural disturbance regimes as a guide for sustainable forest management in Europe. Ecological Applications 32, e2596 (2022).

6. Uhl, B., Schall, P. & Bässler, C. Achieving structural heterogeneity and high multi-taxon biodiversity in managed forest ecosystems: a European review. Biodivers Conserv 34, 3327–3358 (2025).

7. Stritih, A. et al. Can “Closer-to-Nature” Forest Management Sustain Biodiversity and Ecosystem Services in an Uncertain Future? Lessons from Central Europe. Curr. For. Rep. 12, 1 (2026).

8. Vitali, V. et al. Managing European forests for biodiversity conservation and global change adaptation: An assessment of current management concepts. Forest Ecology and Management 611, 123698 (2026).

9. Brunet, J., Fritz, Ö. & Richnau, G. Biodiversity in European beech forests - a review with recommendations for sustainable forest management. Ecological Bulletins 53, 77–94 (2010).

10. Paillet, Y. et al. Biodiversity Differences between Managed and Unmanaged Forests: Meta-Analysis of Species Richness in Europe. Conservation Biology 24, 101–112 (2010).

11. Gossner, M. M. et al. Forest management intensity measures as alternative to stand properties for quantifying effects on biodiversity. Ecosphere 5, art113 (2014).

12. Schall, P. et al. The impact of even-aged and uneven-aged forest management on regional biodiversity of multiple taxa in European beech forests. J Appl Ecol 55, 267–278 (2018).

13. Bae, S. et al. Radar vision in the mapping of forest biodiversity from space. Nat Commun 10, 4757 (2019).

14. Tinya, F. et al. Environmental drivers of forest biodiversity in temperate mixed forests – A multi-taxon approach. Science of The Total Environment 795, 148720 (2021).

15. Wagenaar, L. F., Olsson, O., Stjernman, M. & Smith, H. G. A systematic meta-review: The relationship between forest structures and biodiversity in deciduous forests. Forest Ecology and Management 596, 123072 (2025).

16. Seibold, S. et al. Experimental studies of dead-wood biodiversity — A review identifying global gaps in knowledge. Biological Conservation 191, 139–149 (2015).

17. Doerfler, I., Gossner, M. M., Müller, J., Seibold, S. & Weisser, W. W. Deadwood enrichment combining integrative and segregative conservation elements enhances biodiversity of multiple taxa in managed forests. Biological Conservation 228, 70–78 (2018).

18. Roth, N. et al. Decadal effects of landscape-wide enrichment of dead wood on saproxylic organisms in beech forests of different historic management intensity. Diversity and Distributions 25, 430–441 (2019).

19. Froidevaux, J. S. P., Zellweger, F., Bollmann, K., Jones, G. & Obrist, M. K. From field surveys to LiDAR: Shining a light on how bats respond to forest structure. Remote Sensing of Environment 175, 242–250 (2016).

20. Renner, S. C. et al. Forest structure, plants, arthropods, scale, or birds’ functional groups: What key factor are forest birds responding to? PLOS ONE 19, e0304421 (2024).

21. Hendel, A. et al. Disentangling direct and indirect effects of forest structure on biodiversity: Bottom-up and top-down effects between forestry, bats and their insect prey. Journal of Applied Ecology 62, 93–105 (2025).

22. Kim, H., McComb, B. C., Frey, S. J. K., Bell, D. M. & Betts, M. G. Forest microclimate and composition mediate long-term trends of breeding bird populations. Global Change Biology 28, 6180–6193 (2022).

23. Shipley, B. A New Inferential Test for Path Models Based on Directed Acyclic Graphs. Structural Equation Modeling: A Multidisciplinary Journal 7, 206–218 (2000).

24. Pierick, K. Spatial microclimatic heterogeneity is associated with forest structure and biodiversity. Ecological Indicators 187, 114874 (2026).

25. Yi, H. et al. Tree diversity and mycorrhizal type independently shape multitrophic biodiversity, with stronger effects belowground than aboveground. Nat Ecol Evol 10, 1628–1643 (2026).

26. Antão, L. H. et al. Temperature-related biodiversity change across temperate marine and terrestrial systems. Nat Ecol Evol 4, 927–933 (2020).

27. Loewen, C. J. G., Jackson, D. A. & Gilbert, B. Biodiversity patterns diverge along geographic temperature gradients. Global Change Biology 29, 603–617 (2023).

28. De Frenne, P. et al. Forest microclimates and climate change: Importance, drivers and future research agenda. Global Change Biology 27, 2279–2297 (2021).

29. Weißing, K. K., Pierick, K., Link, R. M., Köhler, M. & Ehbrecht, M. Impacts of forest management on the inter-annual variability of forest microclimate. Eur J Forest Res 145, 34 (2026).

30. Thom, D. et al. Effects of disturbance patterns and deadwood on the microclimate in European beech forests. Agricultural and Forest Meteorology 291, 108066 (2020).

31. Zellweger, F. et al. Forest microclimate dynamics drive plant responses to warming. Science 368, 772–775 (2020).

32. De Lombaerde, E. et al. Maintaining forest cover to enhance temperature buffering under future climate change. Science of The Total Environment 810, 151338 (2022).

33. Sanczuk, P. et al. Microclimate and forest density drive plant population dynamics under climate change. Nat. Clim. Chang. 13, 840–847 (2023).

34. Verdonck, S. et al. Managing canopy cover to preserve forest microclimate and diverse macroarthropod communities in times of drought. Journal of Applied Ecology 62, 2582–2593 (2025).

35. Hähn, G. J. A., et al. Protecting temperate old-growth forests as biotic microrefugia amid climate change. Conservat Sci and Prac 8, e70323 (2026).

36. Sebek, P. et al. Does a minimal intervention approach threaten the biodiversity of protected areas? A multi-taxa short-term response to intervention in temperate oak-dominated forests. Forest Ecology and Management 358, 80–89 (2015).

37. Penone, C. et al. Specialisation and diversity of multiple trophic groups are promoted by different forest features. Ecology Letters 22, 170–180 (2019).

38. Zeller, L. et al. What does literature tell us about the relationship between forest structural attributes and species richness in temperate forests? – A review. Ecological Indicators 153, 110383 (2023).

39. Rothacher, J. et al. The impact of tree mortality and post-disturbance management on insect diversity in temperate forests: Insights from a replicated experiment. Journal of Applied Ecology 62, 1878–1888 (2025).

40. Perlík, M. et al. Canopy openness as the main driver of aculeate Hymenoptera and saproxylic beetle diversity following natural disturbances and salvage logging. Forest Ecology and Management 540, 121033 (2023).

41. Kortmann, M. et al. The positive effects of canopy openness across post-disturbance management on insect diversity. Biological Conservation 313, 111647 (2026).

42. Hilmers, T. et al. Biodiversity along temperate forest succession. Journal of Applied Ecology 55, 2756–2766 (2018).

43. Thom, D. et al. The impacts of climate change and disturbance on spatio-temporal trajectories of biodiversity in a temperate forest landscape. Journal of Applied Ecology 54, 28–38 (2017).

44. Elek, Z. et al. Taxon-specific responses to different forestry treatments in a temperate forest. Sci Rep 8, 16990 (2018).

45. Viljur, M.-L. et al. The effect of natural disturbances on forest biodiversity: an ecological synthesis. Biological Reviews 97, 1930–1947 (2022).

46. Vigués Jorba, J., et al. Differential responses of taxonomic, functional and phylogenetic multi-taxa diversity to environmental factors in temperate forest ecosystems. Ecological Indicators 178, 113855 (2025).

47. Geres, L. S. et al. Macro-and microclimate interactively shape species diversity of multiple taxa in mountain landscapes. Ecography 2025, e07984 (2025).

48. König, S. et al. Micro-and macroclimate interactively shape diversity, niches and traits of Orthoptera communities along elevational gradients. Diversity and Distributions 30, e13810 (2024).

49. Richter, T. et al. Effects of climate and forest development on habitat specialization and biodiversity in Central European mountain forests. Commun Biol 7, 1518 (2024).

50. Richter, T. et al. Landscape context modulates the effect of local canopy cover on forest multidiversity across elevations. Journal of Applied Ecology 62, 3537–3549 (2025).

51. Goßmann, A., Öckinger, E., Schroeder, M., Lindman, L. & Ranius, T. Interaction between regional temperature and shade level shapes saproxylic beetle communities. Diversity and Distributions 30, e13836 (2024).

52. Müller, J. et al. Enhancing the structural diversity between forest patches—A concept and real-world experiment to study biodiversity, multifunctionality and forest resilience across spatial scales. Global Change Biology 29, 1437–1450 (2023).

53. Chao, A. et al. Measuring temporal change in alpha diversity: A framework integrating taxonomic, phylogenetic and functional diversity and the iNEXT.3D standardization. Methods in Ecology and Evolution 12, 1926–1940 (2021).

54. Allan, E. et al. Interannual variation in land-use intensity enhances grassland multidiversity. Proceedings of the National Academy of Sciences 111, 308–313 (2014).

55. Woods, H. A., Dillon, M. E. & Pincebourde, S. The roles of microclimatic diversity and of behavior in mediating the responses of ectotherms to climate change. Journal of Thermal Biology 54, 86–97 (2015).

56. Barahona-Segovia, R. M., Crespin, S. J., Grez, A. A. & Veloso, C. Anthropogenic thermal gradient in managed landscapes determines physiological performance and explains the edge-biased distribution of ectothermic arthropods. Forest Ecology and Management 440, 147–157 (2019).

57. Zellweger, F. et al. Seasonal drivers of understorey temperature buffering in temperate deciduous forests across Europe. Global Ecol Biogeogr 28, 1774–1786 (2019).

58. Decker, O. et al. Three-dimensional temperature variations drive arthropod communities in temperate forests. Forest Ecology and Management 618, 123969 (2026).

59. Müller, J. et al. Weather explains the decline and rise of insect biomass over 34 years. Nature 628, 349–354 (2024).

60. Wolda, H. Insect Seasonality: Why? Annual Review of Ecology and Systematics 19, 1– 18 (1988).

61. Ma, C.-S., Ma, G. & Pincebourde, S. Survive a Warming Climate: Insect Responses to Extreme High Temperatures. Annual Review of Entomology 66, 163–184 (2021).

62. Dormann, C. F. et al. Plant species richness increases with light availability, but not variability, in temperate forests understorey. BMC Ecol 20, 43 (2020).

63. Czyżewski, S. & Svenning, J.-C. Temperate forest plants are associated with heterogeneous semi-open canopy conditions shaped by large herbivores. Nat. Plants 11, 985–1000 (2025).

64. Bradler, P. M. et al. Enhancing structural heterogeneity in managed forest landscapes promotes gamma but not beta diversity in understorey plant communities. Journal of Applied Ecology 63, e70477 (2026).

65. Gossner, M. M. Light intensity affects spatial distribution of Heteroptera in deciduous forests. Eur. J. Entomol. 106, 241–252 (2009).

66. Scherber, C. et al. Bottom-up effects of plant diversity on multitrophic interactions in a biodiversity experiment. Nature 468, 553–556 (2010).

67. Schuldt, A. et al. Multiple plant diversity components drive consumer communities across ecosystems. Nat Commun 10, 1460 (2019).

68. Lettenmaier, L. et al. Light and ungulate browsing interact in shaping future woody plant diversity through natural regeneration. Journal of Applied Ecology 62, 3367–3380 (2025).

69. Pierick, K. et al. Artificial gaps but not thinning persistently enhance forest structural heterogeneity. Eur J Forest Res 145, 68 (2026).

70. Müller, J. et al. LiDAR-derived canopy structure supports the more-individuals hypothesis for arthropod diversity in temperate forests. Oikos 127, 814–824 (2018).

71. Knuff, A. K. et al. Insect abundance in managed forests benefits from multi-layered vegetation. Basic and Applied Ecology 48, 124–135 (2020).

72. Fei, S. et al. Coupling of plant and mycorrhizal fungal diversity: its occurrence, relevance, and possible implications under global change. New Phytologist 234, 1960–1966 (2022).

73. Bässler, C. et al. Enhancing experimentally the structural heterogeneity of forests increase soil fungal diversity but functional lifestyles in contrasting ways. Preprint at 10.1101/2025.08.29.673114 (2025).

74. Kovács, B. et al. The Short-Term Effects of Experimental Forestry Treatments on Site Conditions in an Oak–Hornbeam Forest. Forests 9, 406 (2018).

75. Schwarz, R. et al. Inconsistent short-term effects of enhanced structural complexity on soil microbial properties across German forests. Applied Soil Ecology 214, 106335 (2025).

76. Heidrich, L. et al. Heterogeneity–diversity relationships differ between and within trophic levels in temperate forests. Nat Ecol Evol 4, 1204–1212 (2020).

77. Cordeiro Pereira, J. M., et al. Disentangling the effects of multifunctional forestry practices on the abundances of birds and their invertebrate prey. Ecological Applications 36, e70198 (2026).

78. Farwig, N., Brandl, R., Siemann, S., Wiener, F. & Müller, J. Decomposition rate of carrion is dependent on composition not abundance of the assemblages of insect scavengers. Oecologia 175, 1291–1300 (2014).

79. Hoermann, C. von et al. Effects of abiotic environmental factors and land use on the diversity of carrion-visiting silphid beetles (Coleoptera: Silphidae): A large scale carrion study. PLOS ONE 13, e0196839 (2018).

80. Staab, M. et al. Negative effects of forest gaps on dung removal in a full-factorial experiment. Journal of Animal Ecology 91, 2113–2124 (2022).

81. Büchner, G. et al. Ecological drivers of carrion beetle (Staphylinidae: Silphinae) diversity on small to large mammals. Ecology and Evolution 14, e70203 (2024).

82. Asch, J. et al. Dung beetles do not profit from enhanced spatial heterogeneity in temperate production forests: A forest manipulation experiment. Journal of Applied Ecology 63, e70288 (2026).

83. Barton, P. S. & Evans, M. J. Insect biodiversity meets ecosystem function: differential effects of habitat and insects on carrion decomposition. Ecological Entomology 42, 364– 374 (2017).

84. Hertlein, M. L. et al. Canopy cover and forest management shape vertebrate scavenger assembly but not carrion removal rates. Forest Ecology and Management 593, 122920 (2025).

85. Pincebourde, S., Sinoquet, H., Combes, D. & Casas, J. Regional climate modulates the canopy mosaic of favourable and risky microclimates for insects. Journal of Animal Ecology 76, 424–438 (2007).

86. Bartholomée, O., Jie, V. W., Caplat, P., Smith, H. G. & Baird, E. Warm or bright – Temperature and light microhabitat use in insect pollinators. Ecological Entomology 50, 853–864 (2025).

87. Nowakowski, A. J., Frishkoff, L. O., Agha, M., Todd, B. D. & Scheffers, B. R. Changing Thermal Landscapes: Merging Climate Science and Landscape Ecology through Thermal Biology. Curr Landscape Ecol Rep 3, 57–72 (2018).

88. Mallick, S. et al. Elevation reverses the effects of forest structure on folivory and leaf asymmetry. Functional Ecology 40, 1061–1073 (2026).

89. Lunow, J. et al. Structural characteristics mediate forest mitigation potential against climate change and biodiversity loss. Ecological Applications 36, e70211 (2026).

90. Lettenmaier, L. et al. 12 years of assembly patterns in saproxylic beetles suggest early decay wood as ephemeral resource patch. Journal of Animal Ecology 95, 282–295 (2026).

91. Komonen, A. & Müller, J. Dispersal ecology of deadwood organisms and connectivity conservation. Conservation Biology 32, 535–545 (2018).

92. Rieder, J. S., et al. Canopy gap dynamics contribute to drought-induced defoliation in European beech. Forestry (Lond) 99, cpag060 (2026).

93. Thom, D. & Keeton, W. S. Disturbance-based silviculture for habitat diversification: Effects on forest structure, dynamics, and carbon storage. Forest Ecology and Management 469, 118132 (2020).

94. Hylander, K., Koelemeijer, I. A., Greiser, C. & Christiansen, D. M. Climate adaptation of biodiversity conservation in managed forest landscapes. Conservation Biology 36, e13847 (2022).

95. Seidl, R. et al. Forest disturbances under climate change. Nature Clim Change 7, 395– 402 (2017).

96. Borderieux, J., Gégout, J.-C. & Serra-Diaz, J. M. Extinction drives recent thermophilization but does not trigger homogenization in forest understorey. Nat Ecol Evol 8, 695–704 (2024).

97. Hagge, J. et al. Change in saproxylic beetle, fungi and bacteria assemblages along horizontal and vertical gradients of sun-exposure in forest. Biological Conservation 291, 110493 (2024).

98. Graser, A. et al. Increase in disturbance-induced canopy gaps leads to reorganization of Central European bird communities. Basic and Applied Ecology 83, 88–97 (2025).

99. Achury, R., Staab, M., Blüthgen, N. & Weisser, W. W. Forest gaps increase true bug diversity by recruiting open land species. Oecologia 202, 299–312 (2023).

100. Kortmann, M. et al. Bark-beetle disturbance severity only moderately alters forest affinity of arthropod communities. Insect Conservation and Diversity 17, 579–588 (2024).

101. Massó Estaje, C., et al. Experimental enhancement of structural heterogeneity in forest landscapes promotes multidimensional hoverfly diversity. Journal of Applied Ecology 63, e70252 (2026).

102. Mitesser, O. et al. Forest Heterogeneity by Chain Saw: How Between-Patch Variation in Old Growth Attributes Changes the Metacommunities of Beetles. Ecology Letters 29, e70355 (2026).

103. Schwarz, R. et al. Enhanced forest structural heterogeneity increases functional β-diversity but reduces α-and γ-diversity in soil nematodes. Soil Biology and Biochemistry 214, 110078 (2026).

104. Wild, C. et al. Restoring structural complexity in temperate forests increases bat and bird diversity. Current Biology 36, 2840–2852 (2026).

105. Hill, M. O. Diversity and Evenness: A Unifying Notation and Its Consequences. Ecology 54, 427–432 (1973).

106. Chao, A. & Jost, L. Coverage-based rarefaction and extrapolation: standardizing samples by completeness rather than size. Ecology 93, 2533–2547 (2012).

107. Kortmann, M. et al. A shortcut to sample coverage standardization in metabarcoding data provides new insights into land-use effects on insect diversity. Proceedings of the Royal Society B: Biological Sciences 292, 20242927 (2025).

108. Chiu, C.-H. & Chao, A. Estimating and comparing microbial diversity in the presence of sequencing errors. PeerJ 4, e1634 (2016).

109. Li, D. hillR: taxonomic, functional, and phylogenetic diversity and similarity through Hill Numbers. JOSS 3, 1041 (2018).

110. R Core Team. R: A language and environment for statistical computing. R Foundation for Statistical Computing, Vienna, Austria (2025).

111. Bürkner, P.-C. brms: An R package for Bayesian multilevel models using Stan. Journal of Statistical Software 80, 1–28 (2017).

112. Arif, S. & Massey, M. D. B. Reducing bias in experimental ecology through directed acyclic graphs. Ecology and Evolution 13, e9947 (2023).

113. Makowski, D., Ben-Shachar, M. S., Chen, S. H. A. & Lüdecke, D. Indices of Effect Existence and Significance in the Bayesian Framework. Front. Psychol. 10, (2019).

114. Vehtari, A., Gelman, A., Simpson, D., Carpenter, B. & Bürkner, P.-C. Rank-normalization, folding, and localization: An improved *R̂* for assessing convergence of MCMC. Bayesian Anal. 16, (2021).

115. Gabry, J., Simpson, D., Vehtari, A., Betancourt, M. & Gelman, A. Visualization in Bayesian workflow. J. R. Stat. Soc. A 182, 389–402 (2019).

116. Vehtari, A., Gelman, A. & Gabry, J. Practical Bayesian model evaluation using leave-one-out cross-validation and WAIC. Stat Comput 27, 1413–1432 (2017).

117. Vehtari, A. et. al. loo: Efficient leave-one-out cross-validation and WAIC for Bayesian models. R package version 2.9.0 (2024).

118. Hartig, F. DHARMa: Residual Diagnostics for Hierarchical (Multi-Level /Mixed) Regression Models. R package version 0.4.7 (2024).

119. Rodríguez-Sánchez, F. DHARMa.helpers: Helper functions to check models not (yet) directly supported by DHARMa. R package version 0.0.2 (2025).

## References

Abarenkov, K., Nilsson, R. H., Larsson, K. H., Taylor, A. F., May, T. W., Frøslev, T. G.,…& Kõljalg, U. (2024). The UNITE database for molecular identification and taxonomic communication of fungi and other eukaryotes: sequences, taxa and classifications reconsidered. Nucleic Acids Research 52, D791–D797. 10.1093/nar/gkad1039

Arbeitsgemeinschaft Forsteinrichtung, Arbeitskreis für Standortskartierung (2016). Forstliche Standortaufnahme. 7th edition. IHW-Verlag, Eching bei München.

Aronesty, E. (2013). Comparison of sequencing utility programs. The Open Bioinformatics Journal 7, TOBIOIJ-7-1. 10.2174/1875036201307010001

Bongers, T. (1994). De Nematoden Van Nederland [The Nematodes of the Netherlands], second edition. KNNV Uitgeverij, Zeist.

Caporaso, J. G., Lauber, C. L., Walters, W. A., Berg-Lyons, D., Lozupone, C. A., Turnbaugh, P. J.,…& Knight, R. (2011). Global patterns of 16S rRNA diversity at a depth of millions of sequences per sample. Proceedings of the National Academy of Sciences of the USA 108, 4516–4522. 10.1073/pnas.1000080107

Cesarz, S., Schulz, A.E., Beugnon, R., Eisenhauer, N. (2019). Testing soil nematode extraction efficiency using different variations of the Baermann funnel method. Soil Organisms 91, 61–72. 10.1101/318691

Edgar, R. C. (2013). UPARSE: highly accurate OTU sequences from microbial amplicon reads. Nature methods 10, 996–998. 10.1038/nmeth.2604

Ihrmark, K., Bödeker, I. T., Cruz-Martinez, K., Friberg, H., Kubartova, A., Schenck, J.,…& Lindahl, B.D. (2012). New primers to amplify the fungal ITS2 region–evaluation by 454-sequencing of artificial and natural communities. FEMS Microbiology Ecology 82, 666–677. 10.1111/j.1574-6941.2012.01437.x

Junger, W. (2025). mtsdi: Multivariate Time Series Data Imputation. 10.32614/CRAN.package.mtsdi R package version 0.3.7

Man, M., Kalčík, V., Macek, M., Brůna, J., Hederová, L., Wild, J., & Kopecký, M. (2023). myClim: Microclimate data handling and standardised analyses in R. Methods in Ecology and Evolution 14, 2308–2320. 10.1111/2041-210X.14192

Man, M., Kalčík, V., Macek, M., Brůna, J., Růžičková, A., Wild, J., Kopecký, M. (2025). myClimGui: GUI for myClim package. R package version 1.1.5

Nilsson, R. H., Veldre, V., Hartmann, M., Unterseher, M., Amend, A., Bergsten, J.,…& Abarenkov, K. (2010). An open source software package for automated extraction of ITS1 and ITS2 from fungal ITS sequences for use in high-throughput community assays and molecular ecology. Fungal Ecology 3, 284–287. 10.1016/j.funeco.2010.05.002

R Core Team. 2025. R: A Language and Environment for Statistical Computing. Manual V.4.5.2. R Foundation for Statistical Computing.

Rothacher, J., Seidl, R., Thom, D., Kortmann, M., Chao, A., Chiu,…& Müller, J. (2025). The impact of tree mortality and post-disturbance management on insect diversity in temperate forests: Insights from a replicated experiment. Journal of Applied Ecology 62, 1878–1888. 10.1111/1365-2664.70086

Sagova-Mareckova, M., Cermak, L., Novotna, J., Plhackova, K., Forstova, J., & Kopecky, J. (2008). Innovative methods for soil DNA purification tested in soils with widely differing characteristics. Applied and Environmental Microbiology 74, 2902–2907. 10.1128/AEM.02161-07

Schwarz, R., Bradler, P. M., Chao, A., Chuang, P.-Y., Ciobanu, M., Decker, O.,…& Cesarz, S. (2026). Enhanced forest structural heterogeneity increases functional β-diversity but reduces α-and γ-diversity in soil nematodes. Soil Biology and Biochemistry 214, 110078. 10.1016/j.soilbio.2025.110078

Szabó-Takács, B., Farda, A., Zahradníček, P., & Štěpánek, P. (2015). Continentality in Europe according to various resolution regional climate models with A1B scenario in the 21st century. Időjárás 119, 515–535

Větrovský, T., Baldrian, P., Morais, D. (2018). SEED 2: a user-friendly platform for amplicon high-throughput sequencing data analyses, Bioinformatics 34, 2292–2294. 10.1093/bioinformatics/bty071

Wild, C., Chao, A., Chuang, P.-Y., Cadotte, M., Daume, N., Decker, O.,…& Müller, J. (2026). Restoring structural complexity in temperate forests increases bat and bird diversity. Current Biology 36, 2840–2852.e4. 10.1016/j.cub.2026.04.058

Žifčáková, L., Větrovský, T., Howe, A., & Baldrian, P. (2016). Microbial activity in forest soil reflects the changes in ecosystem properties between summer and winter. Environmental Microbiology 18, 288–301. 10.1111/1462-2920.13026

## S3.3 Supplementary references

Allan, Eric, Oliver Bossdorf, Carsten F. Dormann, et al. 2014. “Interannual Variation in Land-Use Intensity Enhances Grassland Multidiversity.” Proceedings of the National Academy of Sciences 111 (1): 308–13. 10.1073/pnas.1312213111.

Bürkner, Paul-Christian. 2018. “Advanced Bayesian Multilevel Modeling with the R Package brms.” The R Journal 10 (1): 395–411. 10.32614/RJ-2018-017.

Gabry, Jonah, Daniel Simpson, Aki Vehtari, Michael Betancourt, and Andrew Gelman. 2019. “Visualization in Bayesian Workflow.” Journal of the Royal Statistical Society: Series A (Statistics in Society) 182 (2): 389–402. 10.1111/rssa.12378.

Gelman, A, J B Carlin, H S Stern, et al. 2014. Bayesian Data Analysis, Third Edition. 3rd ed. Taylor & Francis Group, LLC.

Hartig, Florian. 2024. DHARMa: Residual Diagnostics for Hierarchical (Multi-Level / Mixed) Regression Models. R Package Version 0.4.7. Manual. 10.32614/CRAN.package.DHARMa.

Hoffman, Matthew D, and Andrew Gelman. 2014. “The No-U-Turn Sampler: Adaptively Setting Path Lengths in Hamiltonian Monte Carlo.” Journal of Machine Learning Research 15 (1): 1593–623.

Kubinec, Robert. 2022. “Ordered Beta Regression: A Parsimonious, Well-Fitting Model for Continuous Data with Lower and Upper Bounds.” *Political Analysis*, July, 1–18. 10.1017/pan.2022.20.

Lewandowski, Daniel, Dorota Kurowicka, and Harry Joe. 2009. “Generating Random Correlation Matrices Based on Vines and Extended Onion Method.” Journal of Multivariate Analysis 100 (9): 1989–2001.

Makowski, Dominique, Mattan S. Ben-Shachar, S. H. Annabel Chen, and Daniel Lüdecke. 2019. “Indices of Effect Existence and Significance in the Bayesian Framework.” Frontiers in Psychology 10 (December). 10.3389/fpsyg.2019.02767.

Rodríguez-Sánchez, Francisco. 2026. DHARMa.helpers: Helper Functions to Check Models Not (yet) Directly Supported by DHARMa. R Package Version 0.0.2. Manual.

Shipley, Bill. 2000. “A New Inferential Test for Path Models Based on Directed Acyclic Graphs.” Structural Equation Modeling: A Multidisciplinary Journal 7 (2): 206–18. 10.1207/S15328007SEM0702_4.

Sivula, Tuomas, Måns Magnusson, Asael Alonzo Matamoros, and Aki Vehtari. 2025. “Uncertainty in Bayesian Leave-One-Out Cross-Validation Based Model Comparison.” *Bayesian Analysis*, ahead of print, January. 10.1214/25-BA1569.

Vehtari, Aki, Jonah Gabry, Måns Magnusson, et al. 2025. Loo: Efficient Leave-One-Out Cross-Validation and WAIC for Bayesian Models. R Package Version 2.9.0.

Vehtari, Aki, Andrew Gelman, and Jonah Gabry. 2017. “Practical Bayesian Model Evaluation Using Leave-One-Out Cross-Validation and WAIC.” Statistics and Computing 27 (5): 1413–32. 10.1007/s11222-016-9696-4.

Vehtari, Aki, Andrew Gelman, Daniel Simpson, Bob Carpenter, and Paul-Christian Bürkner. 2020. “Rank-Normalization, Folding, and Localization: An Improved Rhat for Assessing Convergence of MCMC.” Bayesian Analysis, ahead of print. 10.1214/20-BA1221.

Wickham, Hadley, Mara Averick, Jennifer Bryan, et al. 2019. “Welcome to the tidyverse.” Journal of Open Source Software 4 (43): 1686. 10.21105/joss.01686.

